# Hierarchical classification of immune cell transcriptomes at population-scale

**DOI:** 10.64898/2026.05.30.728980

**Authors:** Christopher Beltz, Zekai Qiu, Leon Sadowski, Joscha A. Kraske, Anmol Aggarwal, Alvaro Quintanal-Villalonga, Parvathy Manoj, Andrea Littbarski, Sunanjay Bajaj, Brigita Meškauskaitė, Shigeaki Umeda, Linas Mazutis, Samuel A. Rose, Joseph M. Chan, Tal Nawy, Juozas Nainys, Ronan Chaligne, Elisa de Stanchina, Katharina A. Kälber, Christiane S. Cussigh, Stefan M. Kallenberger, Anja Williams, Maximilian Jenzer, Tillmann Pompecki, Steffen Kahle, Nicolas Hohmann, Daniel P. Nussbaum, Nelson S. Moss, Etay Ziv, Anne K. Berger, Georg M. Haag, Christoph Springfeld, Stefanie Zschäbitz, Jessica C. Hassel, Jürgen Debus, Dirk Jäger, Christine A. Iacobuzio-Donahue, Karuna Ganesh, Dana Pe’er, Guy Ungerechts, Charles M. Rudin, Peter E. Huber, Thomas Walle

## Abstract

Accurate immune cell classification is essential for interpreting single-cell RNA sequencing (scRNA-seq) data. However, progress in automating cell type annotation is constrained by the lack of independent, high-resolution benchmarks, as routine data integration introduces statistical dependencies that inflate model generalizability. Here, we present the single-cell universal classification omnibus (Suco), a resource of independent, uniform expert annotations, and Compocyte, a modular hierarchical classifier. Together, they establish a framework that substantially outperforms existing classifiers while facilitating expert review of ambiguous annotations. Applying Compocyte across 50 studies, including three newly generated datasets, we classified 15.6 million leukocytes from 3,965 patients. Within this cohort, we identified a new tumor-associated resorptive macrophage phenotype, a non-canonical monocyte subtype in subclinical cytokine release syndrome, and the programmatic erosion of T cell memory stemness across metastatic sites. Suco and Compocyte thus provide a generalizable framework to uncover the principles governing human immunity at population scale.

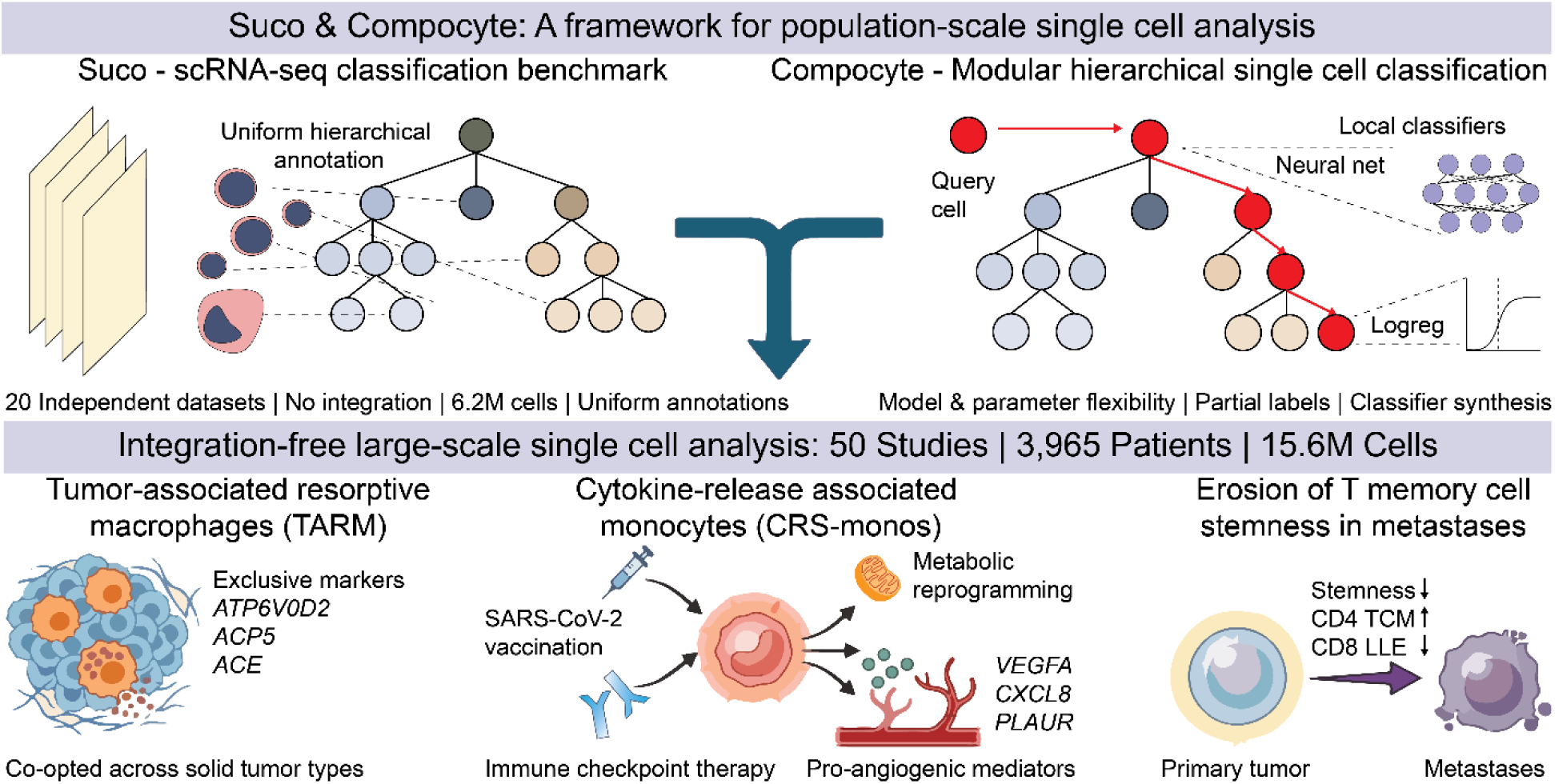

**In brief:** The single-cell universal classification omnibus and the modular hierarchical classifier Compocyte enable annotating single cell RNA sequencing data from 3,965 patients, revealing novel resorptive macrophage and vaccination-associated monocyte states, alongside the erosion of T cell memory stemness as a hallmark of solid tumor metastases.

**Highlights:**

- Suco, a benchmark enabling novel single cell artificial intelligence models
- Compocyte, a hierarchical cell type classifier outperforming current architectures
- Macrophages adopt osteoclast-like gene expression states across cancer types
- Stem-like programs erode in metastasis-infiltrating T memory cells across tumors

## Introduction

Developing robust (semi-)supervised machine learning (ML) models requires strict adherence to the steps of training, validation, and external testing^1^. Central to this process is the statistical independence of test data, which prevents models from relying on confounding patterns that fail to generalize. In computer vision, high-quality, independently labeled resources like ImageNet have been pivotal in driving major advances^2^. More recently, human-in-the-loop approaches have further accelerated this progress by focusing expert annotation on the instances most challenging for automated systems^3^.

In many areas of biology, comparable resources are conspicuously absent. This "benchmarking crisis"^4^ is particularly acute in single-cell RNA sequencing (scRNA-seq), where cell-type annotation requires navigating millions of cells across thousands of sparse features. Cell type annotation is a key initial step for biological interpretation and many downstream analyses such as cell type compositional analysis^5^, differential expression analysis^6^, or cell type aware factor analysis^7^. Given its complexity, cell type classification has also become an important benchmark for generative single cell genomics artificial intelligence (AI) models^8-10^. Because manual annotation is time-consuming and requires significant domain expertise^11^, researchers often resort to data integration to project multiple datasets into a shared latent space for joint clustering and annotation^12-14^. While efficient in manual labor, this practice introduces statistical dependencies between training and test data^1^, leading to an overestimation of a classifier’s ability to generalize to truly unseen biological contexts.

The challenge is compounded by the fluid and diverse nature of immune cell ontologies. Immunological ontologies are subject to frequent refinement, as seen in the expansion of T helper cell subsets^15^, the discovery of diverse innate lymphoid cell (ILC) populations^16^, and the transition to modular cell types^17^. Single-cell genomics has further accelerated this evolution by providing unprecedented phenotypic depth. However, existing automated classifiers often lack the flexibility to adapt to these shifting ontologies. Even advanced transfer learning models utilizing query adapters while accounting for batch effects struggle to formally integrate newly discovered cell states^18^. Ultimately, appending new labels to these architectures requires computationally intensive retraining that risks catastrophic forgetting or destabilizes performance on previously established categories^19^.

Here, we present a unified classification framework for population-scale human immunology comprised of three pillars: The single-cell universal classification omnibus (Suco)^20,21^, the first resource of independent but uniform expert annotations (10.5281/zenodo.13709549 & 10.5281/zenodo.15350417), Cytopus 2, a knowledge base for transparent, marker-based labeling of single cell ontologies (https://github.com/wallet-maker/cytopus), and Compocyte, a modular hierarchical cell type classifier that outperforms existing automated classification methods with open-source pretrained models for human immunobiology (https://github.com/WALL-E-Lab/Compocyte). Suco provides 6.2 million independently annotated leukocyte profiles from 776 individuals across 20 datasets, establishing a rigorous benchmark for generalizability. By utilizing Cytopus 2 to manually label over 2,500 cell clusters, we identified conserved immune states that are frequently lost during traditional data integration, including a tumor-enriched resorptive macrophage phenotype and Langerhans-like cells across multiple solid tumors.

After training Compocyte on the Suco resource, we deployed it across an expansive cohort of 50 studies, including the entire Human Tumor Atlas Network (HTAN) corpus^22^. This allowed us to map the immune landscape of 15.6 million cells from 3,965 patients with unprecedented resolution. This population-scale analysis revealed a non-canonical monocyte subtype associated with subclinical cytokine release syndrome and the erosion of T cell memory stemness in metastases across solid tumors. Together, Suco and Compocyte provide a generalizable machine learning framework and benchmark resource that enables the reliable, high-resolution annotation of human immunology at a global scale.

## Results

### Graph-based ontology annotations enable the single cell universal classification omnibus

Biological cell types exist within developmental and phenotypic hierarchies. However, standard single-cell analysis frameworks (e.g., AnnData^23^, SingleCellExperiment^24^) predominantly represent these relationships as flat, tabular data. This structural flattening risks annotation inconsistencies such as assigning disjoint lineage labels to the same cell and complicates lineage-aware queries. To resolve this, we developed *Cytopus 2* (https://github.com/wallet-maker/cytopus), a knowledge base that models immune ontologies as a graph, explicitly linking all cells to their precise coordinates within the hierarchy.

Furthermore, defining cell identities based on single marker genes remains uniquely vulnerable to single-cell artifacts like dropout, ambient RNA, and multiplet formation. *Cytopus 2* overcomes this by defining cell types through hierarchically contextualized functional marker gene signatures (**Table S1**). Rather than relying on absolute markers, the framework utilizes both positive markers (genes critical to identity) and negative markers (genes incompatible with the identity) that are evaluated against a cell type’s direct siblings within the ontology graph. In total, this knowledge base encompasses >2000 marker genes defining 161 human cell types including 131 leukocyte types (**Fig. 1a**, **Methods**, **Table S1**). Because the distinguishing power of a marker is fundamentally dependent on the reference population, this hierarchical anchoring provides a highly robust foundation for single-cell annotation.

**Fig. 1.**
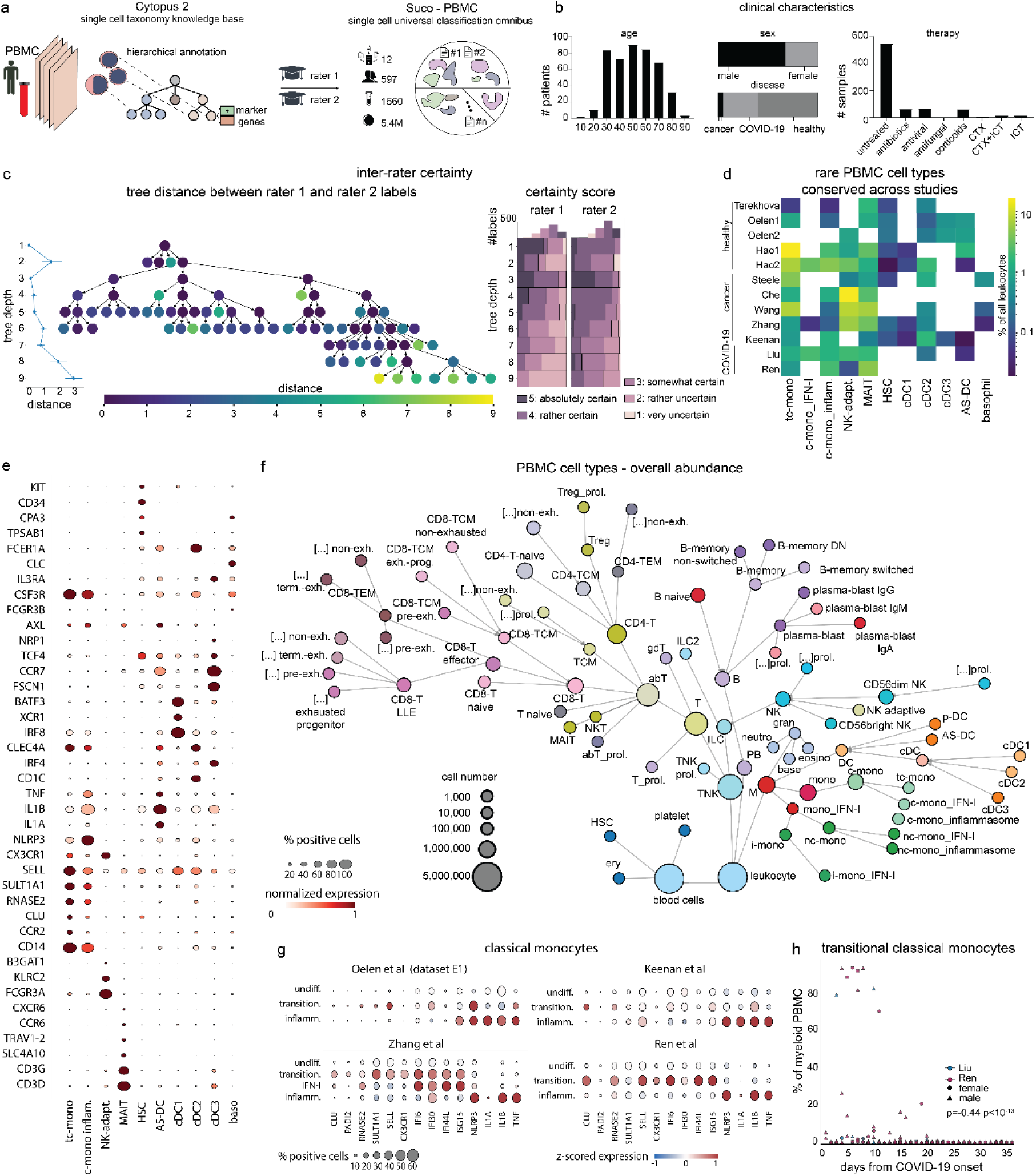
Suco standardizes human peripheral blood mononuclear cells for automated hieararchical single cell classification. **a,** To generate the single cell universal classification omnibus (Sue□), we individually preprocessed 12 human peripheral blood mononuclear cells (PBMC) scRNA-seq dataset. Two raters indepedently assigned hierachical cell type labels using marker genes from our Cytopus 2 knowledge base followed by homogenizing annotations by inter-rater agreenment. **b,** Patients in Suco show a broad age range and mostly untreated samples from various diseases. **c,** Annotations from independent raters differed more with increasing granularity by mean (+/- 95% confidence interval!) ontology tree distance (left panel) at each tree level, or each cell type label (middle panel). Perceived uncertainty as measured on a five-point Likert scale (right panel) increase with increasing cell type granularity. **d,** Suco identified several rare cell types conserved across studies which were not consistently annotated in the original studies. **e,** Marker genes for selected cell types identified in Suco across the 12 PBMC datasets. tc: transitional classical. mono: monocyte, NK: Natural Killer, MAIT: mucosa-associated invariant chain T cell, AS DC (AXL & SIGLEC6 positive dendritic cell), baso (basophil), cDC (conventional dendritic cell). HSC (hematopoietic stem cell), MAIT (mucosa associated invariant chain T cell), mono (monocyte), NK (Natural Killer). tc (transitional classical). **f,** Ontological relationship and abundance of cell types in Suco. The cell number of each parent node includes the sum of all cells of their child nodes. […] (add annotation from parent label), B (B cell), c mono (classical monocyte), DN (double negative), eosino (eosinophil), ery (erythrocyte), exh. (exhausted), gdT (gamma delta T cell). gran (granulocyte). IFN (interferon), ILC (innate lymphoid cell), i mono (intermediate monocyte), M (myeloid cell), nc mono (non classical monocyte), neutro (neutrophil), PB (plasma or B lineage), prol. (proliferating), T (T cell). TCM (T central memory). le-mono (lransitional classifcal monocyte) TEM (T effector memory). term (terminally). TNK (Tor innate lymphoid cell). Treg (regulatory T cell). g. Suco identified new subsets of classical monocytes with characteristic markers genes highlighted in the dotplot including transitional monocytes (transition.), inflammasome-activated monocytes (inflamm.), interferon activaled monocytes (IFN-I) and undifferentiated monocytes (undif.). **h,** transitional monocytes are associated with time from COVID-19 onset (by symptom or positive polymerase chain reaction test whichever came first) with p values calculated using Spearman rank.

We leveraged Cytopus 2 to construct the single-cell universal classification omnibus (Suco, **Fig. 1a**), an independently but uniformly annotated resource designed to bypass the statistical dependencies and loss of rare cell types inherent to joint data integration (e.g. loss of cDC1, **Fig. S1**)^25^. We focused initially on peripheral blood mononuclear cells (PBMCs) due to their phenotypic granularity, diagnostic accessibility, and relevance across diverse diseases. To rigorously test a model’s ability to generalize across diverse clinical and biological conditions, we aggregated 12 high-quality PBMC datasets totaling 5.4 million cells from 1,560 samples across 597 individuals (**Fig. 1a**). Suco spans a wide age demographic (6 to 92 years) and a broad clinical spectrum with healthy donors (60%), patients with solid tumors (5%), and patients with acute COVID-19 (30%, **Fig. 1b**). Suco also includes diverse therapeutic interventions ranging from corticosteroids (8%) and antibiotics (8%) to immune checkpoint inhibitors (4%). Reflecting the underlying disease prevalence of cancer and COVID-19, we observed a slight male predominance (67%, **Fig. 1b**).

To preserve statistical independence, each dataset was processed and clustered individually (**Methods**). Through iterative rounds of subsetting, re-normalization, and dimensionality reduction, we isolated 1,315 highly resolved PBMC clusters (**Table S3**). To benchmark human baseline accuracy, these clusters were labeled independently by two computational immunologists using the Cytopus 2 framework with discordant labels being harmonized through joint review and consensus. Systematic evaluation of these annotations revealed a critical bottleneck in manual single-cell analysis: Both inter-rater agreement and self-reported certainty scores (measured via a five-point Likert scale) decreased at lower levels of the cell type ontology (**Fig. 1c**). This divergence was particularly pronounced within the memory T cell compartment demonstrating a strict necessity for the standardized, hierarchical annotation framework established by Suco and Compocyte.

### Suco identifies novel conserved peripheral blood immune states across diseases

Leveraging the high resolution of the Suco reference, we next identified rare or previously uncharacterized immune states conserved across the 12 independent PBMC datasets which were not consistently annotated in the original publications^26-35^. These included adaptive NK cells^36^, mucosal-associated invariant T (MAIT) cells^37^, basophils, hematopoietic stem cells, and distinct dendritic cell (DC) subsets (conventional DCs type 1 (cDC1^38^) and 2 (cDC2^38^), migratory DCs (cDC3^38^), and AXL^+^ SIGLEC6^+^ DCs (AS-DCs)^39^) (**Figure 1d–f, S2a**). To interpret the functional significance of these conserved states, we systematically evaluated marker genes recorded by the independent raters (**Table S3**). This approach revealed previously unappreciated functional programs. AS-DCs exhibited a core cDC2 transcriptional phenotype superimposed with robust NLRP3 inflammasome activation, suggesting these cells represent cDC2s actively responding to danger-associated molecular patterns (**Figure 1e, S2a**).

Beyond identifying rare lineages, the granular resolution of Suco allowed us to systematically characterize exhaustion programs across the memory T cell compartment^17,40^. This high-resolution mapping revealed that exhaustion gene programs are not uniformly acquired but are instead preferentially enriched within specific memory subtypes (**Figure 1f**, **S2b**). CD8 long-lived effectors spanned the full spectrum from progenitor exhaustion^41^ (with high *TCF7*, *CTLA4, TOX*), over pre-exhaustion (also called intermediate exhaustion^42^ with high *GZMK*, *EOMES*, low *GZMB* & *PRF1*), to terminal exhaustion states^43,44^ (with high *TOX*, *PDCD1*, *ENTPD1*, and *LAG3,* **Fig. S2b**). By contrast, CD8 T central memory cells showed either pre-exhausted or progenitor-exhaustion states while the remaining CD4 & CD8 T cell memory types showed no transcriptomic sign of exhaustion highlighting a strong relationship between ontogeny and exhaustion trajectory. Hence, by linking established concepts of exhaustion with rigorous, large-scale data analysis, our results facilitate the targeting of distinct exhaustion programs within specific T cell types.

Within the myeloid compartment, this high-resolution mapping resolved multiple distinct functional states of classical monocytes, including inflammasome-activated and type-I interferon (IFN-I)-activated subsets (**Fig. 1g**). We also identified a highly conserved, non-canonical classical monocyte subpopulation we term "transitional classical monocytes" (tc-monos). Molecularly, tc-monos exhibit a unique transcriptional signature at the interface of inflammation and tissue repair (**Fig. 1g, S2a**). Rather than expressing canonical interferon (*IFI6, IFI30, IFI44L, ISG15*)^45^ or inflammasome (*NLRP3, IL1A, IL1B, TNF*)^46^ genes, they transcribe immune-modulating factors like *PADI2*^47^ and the alarmin *RNASE2*^48^, alongside tissue repair and detoxification genes (*CLU*^49,50^ & *SULT1A1*^51^, **Fig. 1g, S2a**). In line with their classical monocyte identity tc-monos lack *FCGR3A*, but selectively express *CX3CR1 (***Fig. 1g**), a chemokine receptor typically restricted to non-classical or intermediate monocytes^52^. The absence of key dendritic cell transcripts (*CD1A*, *NRP1, XCR1, CD1C*)^39^ definitively separates tc-monos from dendritic cells (**Fig. 1e**, **S2**), confirming them as a conserved myeloid state. Kinetic analysis of acute COVID-19 patients revealed that tc-monos peak around 7 days post-infection (**Fig. 1h**). Because this coincides with a critical inflection point in COVID-19 biology where systemic cytokine levels begin to recede^53-55^, we hypothesize these cells play a role in curbing acute inflammation. While predominant in COVID-19, we also found tc-monos in the blood of cancer patients and healthy donors suggesting that they represent an integral cell type rather than an artifact of extreme pathology (**Fig. S2c**).

### Suco identifies rare tumor infiltrating immune cell types conserved across solid tumors

We incorporated tumor infiltrating leukocytes (TIL) into Suco due to their central role in cancer and the inherent technical challenge of resolving subtle variations amidst strong technical batch effects which occur due to tissue dissociation. We included 8 independent high-quality scRNA-seq datasets^26,28,29,34,56-59^ across breast, colorectal, and pancreatic cancers which enabled the identification of both conserved and tumor specific cell populations (**Fig. 2a**). The cohort reflected a representative oncology demographic with a median age of 58 years ranging from 30 to 90 years (**Fig. 2b**). Most leukocytes derived from primary tumors (60%), but we also included metastases (25%) and adjacent normal tissues (15%, **Fig. 2b**). Samples were predominantly collected at baseline (62%) alongside cohorts receiving chemotherapy (21%), immune checkpoint therapy (ICT, 11%), or combination regimens (6%, **Fig. 2b**).

**Fig. 2.**
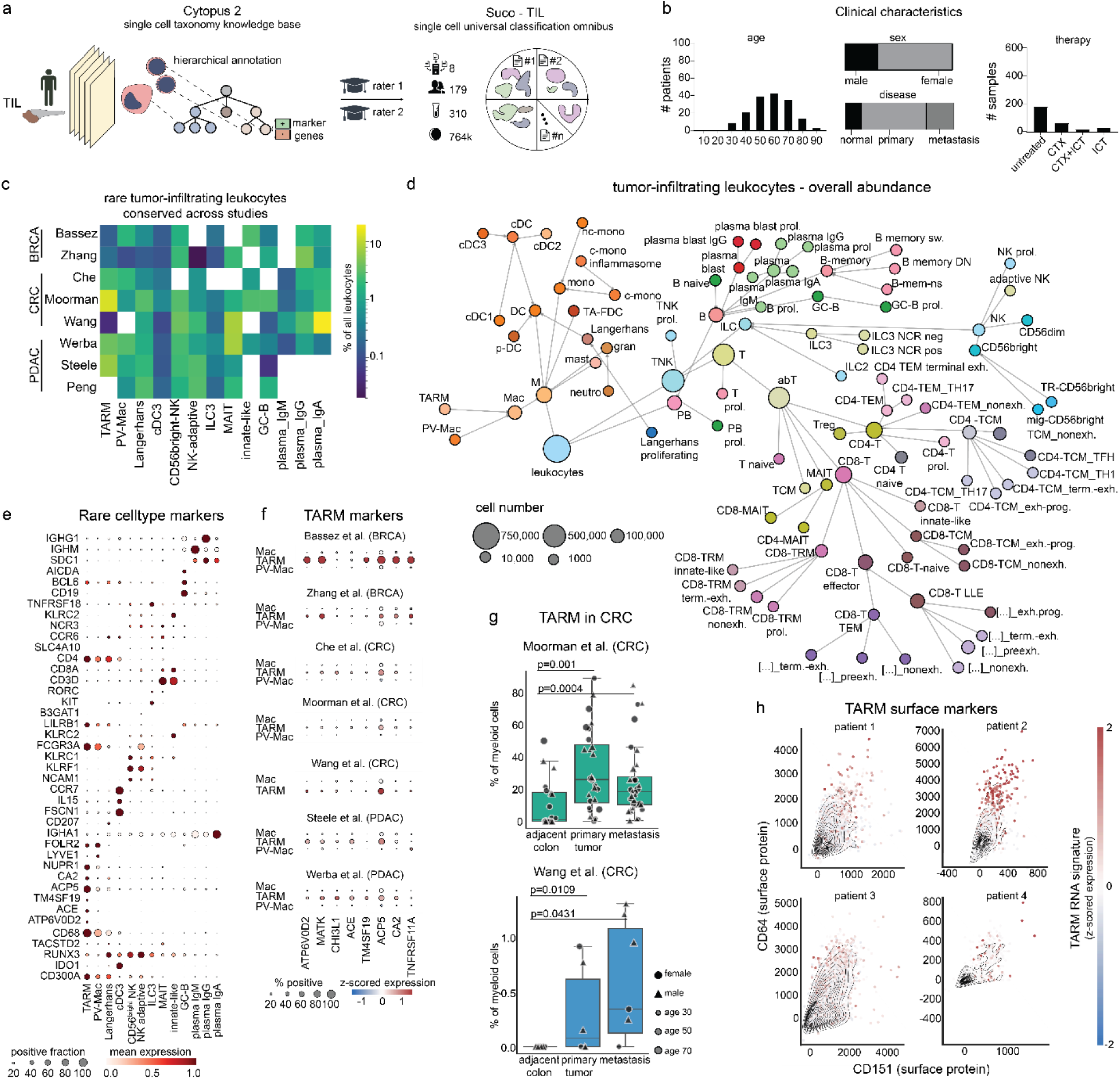
Suco identifies tumor-infiltrating leukocyte phenotypes conserved across cancer types. **a**, We individually preprocessed 8 scRNA-seq dataset from human tumor-infiltrating leukocytes (TIL). Two raters then indepedently assigned cell types hierarchically using marker genes from our Cytopus 2 knowledge base followed by homogenizing annotations by inter-rater agreenment generating the Suco TIL resource for benchmarking cell type annotation methods. **b,** Patients in Suco show a typical age range for cancer patients and samples from chemotherapy (CTX), immune checkpoint therapy (ICT) and untreated samples from various tumor types including metastatic samples. **c,** Suco identified several rare cell type conserved across studies and often missed by the original study authors. **d,** Hierachical relationship and cell numbers of different cell types in the Suco hierarchy. The cell number of each parent node includes the sum of all cells of their child nodes. c- (classical), cDC (conventional dendritic cell), DN (double negative), exh (exhausted), GC (germinal center B cells), gran (granulocyte), ILC (innate lymphoid cell), innate like (innate like T cells), LLE (long lived effector), M (myeloid), Mac (Macrophage), MAIT (mucosa associated invariant chain T cells), mem (memory), mono (monocyte), nc (non classical), neutro (neutrophil), NK (Natural Killer cell), ns (non switched), PB (plasma or B cell lineage), plasma (plasma cell), prog (progenitor), prol. (proliferating), PV (perivascular), sw. (switched), T (T cell), TA FDC (tumor associated follicular dendritic cell), TARM (tissue associated resorptive macrophages), TEM (T effector memory), TH17 (T helper 17), TNK (Tor ILC lineage). **e,** Marker genes for selected cell types identified in Suco across the 12 PBMC datasets. **f,** Suco identified TARM with characteristic markers genes as highlighted in the dotplot by average z-scored (across all cells) expression. **g,** TARM were enriched in colorectal cancer primaries and liver metastases as compared to adjacent normal tissue. P values were calculated using Wilcoxon-matched-pairs signed rank tests. **h,** TARM showed surface expression of CD64 and CD151 as assessed by independent CITE-seq data from n;4 primary pancreatic cancer patients.

Utilizing independent rater systematic annotation via Cytopus 2 without prior dataset integration ensured that the observed cell types reflected genuine biology rather than algorithmic artifacts (**Table S4**). This strategy successfully unmasked rare populations often obscured by integration noise such as type 2 innate lymphoid cells (ILC2, **Fig. 2c-e**, **S3**). Furthermore, CD56^bright^ NK cells frequently exhibited a tissue resident phenotype marked by *ITGAE* and *ZNF683* alongside an absence of SELL (**Fig. 2c-e**, **S4**). MAIT cells were universally present across primary and metastatic tumors despite not being consistently described in the original studies^59,60^ (**Fig. 2c**). We also reliably identified B cell subsets, Including germinal center B cells and isotype-specific (IgG, IgA, or IgM) plasma cells (**Fig. 2c**), suggesting that class-switch status is a primary driver of their transcriptomic profiles.

Moreover, dendritic cells exhibiting a cDC2 phenotype (high *CLEC4A, CLEC10A, IRF4, CD1C*, and *FCER1A*) frequently expressed genes of Langerhans identity (*CD207, CD1A*) which are typically found in the skin (**Fig. 2e, S4**)^38,61-63^. We show that these visceral Langerhans cells express *TACSTD2*, making them a potential target of standard of care TROP2 antibody drug conjugates with implications for immune checkpoint therapy combinations (**Fig. 2e**)^64^. Furthermore, their elevated *IDO1*, *CD300A*, and *RUNX3* expression partially align them with the previously identified cDC2B subset (**Fig. S2e, S4**)^65^. Within the T cell compartment, exhaustion states showed strong memory-cell preferences, with only CD8 LLEs spanning the full progenitor-to-terminal exhaustion spectrum we also described in PBMC (**Fig. 2d, S4, S5a**). However, exhaustion was significantly enriched in TILs as compared to PBMC and expanded beyond CD8 T cells to include CD4 TCMs and TEMs (**Fig. 2d**, **Fig. S5a**), likely driven by the immunosuppressive tumor microenvironment. While pre-exhausted CD8 LLEs dominated the CD8 compartment, the CD4 compartment primarily comprised progenitor-exhausted CD4 TCMs (**Fig. 2d**, **Fig. S5a**). Ultimately, the tumor microenvironment drives both the expansion and diversification of exhaustion states across memory T cell subsets.

### Adoption of osteoclast programs in macrophages across tumor types

Shifting focus to the myeloid compartment, hierarchical annotation refined macrophage categorization into three major phenotypic states across tumor types: undifferentiated macrophages (which did not show consistent marker expression patterns), perivascular macrophages^66^, and tumor-associated resorptive macrophages (TARM) (**Fig. 2e,f**). Examining these subsets in detail, we showed that perivascular macrophages correlated positively with patient age in female but not male cohorts possibly driven by the age-dependent hormonal changes and effects on macrophage extravasation^67^ (**Fig. S5b**). Furthermore, this cross-cancer analysis unveiled TARM as a novel, osteoclast-like phenotype spanning eight of nine colorectal (CRC), pancreatic, and breast cancer datasets (**Fig. 2c**). TARM were significantly enriched in neoplastic tissue relative to adjacent normal regions in CRC suggesting disease-specific roles in cancer (**Fig. 2g**).

Because standard differential expression analyses often struggle to resolve phenotypic divergence and frequently highlight broadly expressed genes, we utilized an entropy-based exclusivity metric (**Methods**) to determine truly distinctive gene expression patterns characterizing TARM. This approach successfully identified TARM-exclusive functional drivers essential for bone resorption and extracellular matrix remodeling, including *ATP6V0D2^68^*, *ACP5^69^, CA2^70^, ACE^71^,* and *TM4SF19*^72^ (**Fig. 2f**). In breast cancer, these cells also express RANK (*TNFRSF11A*) revealing a potential therapeutic vulnerability (**Fig. 2f**)^73^. The cell type fidelity of these markers (Gini coefficients 0.97) contrasted with the more homogeneous expression of classical macrophage markers such as *SPP1* (Gini=0.68) and *TREM2* (Gini=0.72) (**Fig. S5c**). The reproducibility of TARM marker gene expression patterns contrasted with the broad range of macrophage labels assigned by original authors’ data, reflecting the poor definition of macrophage types in human tumors^74^ and underlining the need for rigorous marker identification in independent studies. Furthermore, TARM entirely lacked canonical dendritic cell signatures (*CD1C, CD1A, BATF3, IRF7, IRF4, IRF8,* **Fig. S4**) and were identified by the concurrent surface expression of CD64 and CD151 in CITE-seq data (**Fig. 2h**), thus establishing TARM as a separate cell type identity.

To integrate TARM into the broader cancer ecosystem, we mapped their co-occurrence with other immune cell types, revealing four conserved tumor immune microenvironments (TIME) (**Fig. S6**) including a recirculating TIME (lymphoid-homing naive and memory T/NK cells), an adaptive resistance niche (regulatory/exhausted T cells and dendritic cells), and an innate-like TIME (MAIT, resident NK, and exhausted progenitor T cells). TARM correlated with a fourth lymphoid constructor TIME comprising T follicular helper cells, NCR2^+^ ILC3s, and Th17 cells (**Fig. S6**). This reveals an intriguing ecological link, where the accumulation of matrix remodeling TARM closely parallels the local assembly of the cellular building blocks for tertiary lymphoid structure formation.

### Compocyte enables hierarchical classification across modifiable ontologies

Assembling multi-dataset atlases without the statistical confounders of latent-space integration requires classification frameworks capable of adapting to fluid immunological ontologies. Because standard models generally cannot combine profound taxonomy modifications with hierarchical annotation, we developed Compocyte (Composite classification of cytological taxonomies) (**Methods**). Compocyte utilizes a supervised, hierarchical composite architecture mapped to a user-defined ontology. At each ontology branching point, an independent local classifier partitions cells into subsets and passes them to downstream nodes (**Fig. 3a**). Because these local classifiers function independently, individual hierarchy components can be exchanged or updated without disturbing the global model. Furthermore, Compocyte assigns the optimal machine learning model to each node based on the available training data via an integrated hyperparameter optimization module. Classical models (e.g., logistic regression, boosted trees) are usually prioritized for rare cell types with limited data, whereas neural networks are deployed for data-abundant nodes to maximize classification accuracy (**Methods**).

**Fig. 3.**
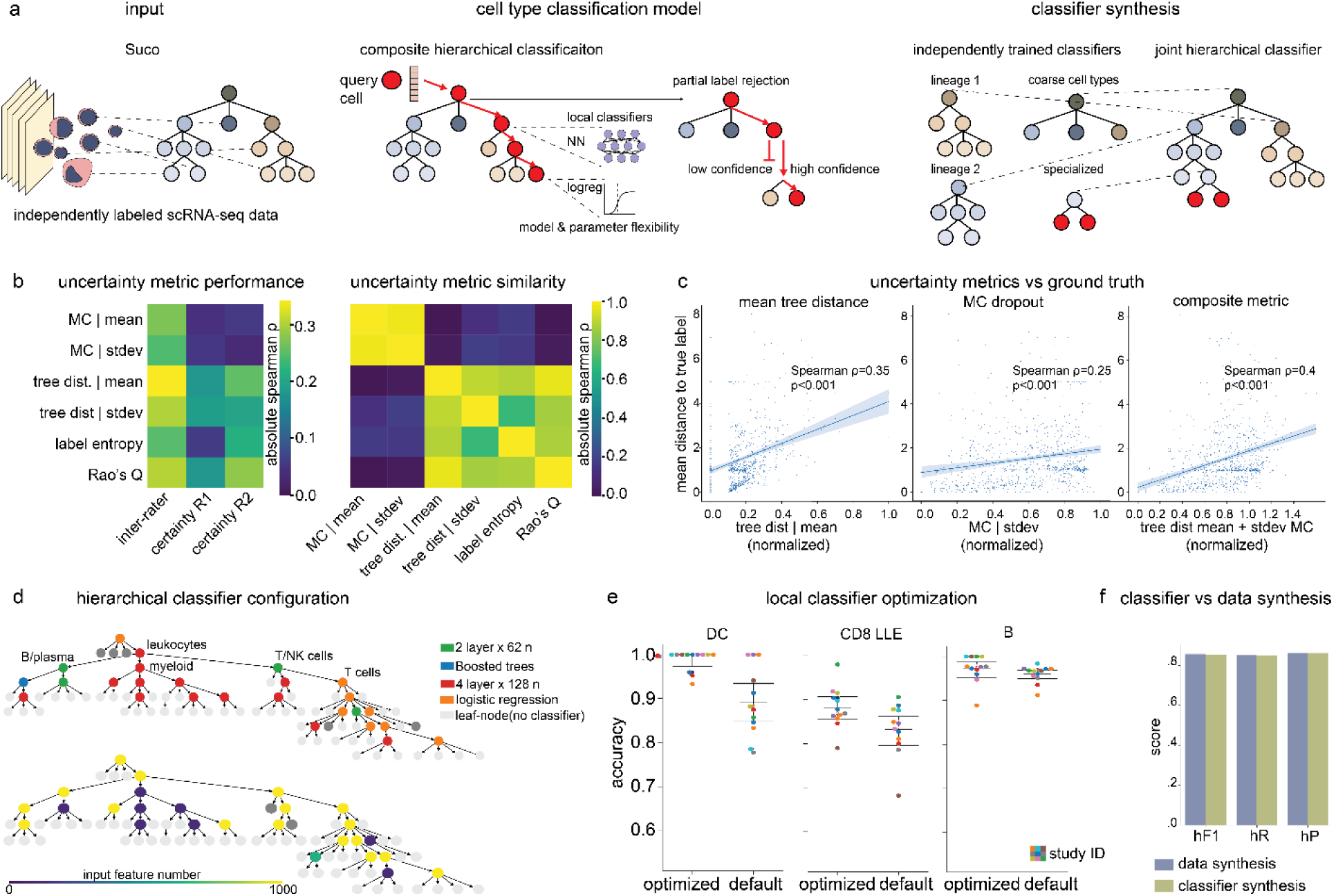
Compocyte enables modular hierarchical cell type classification. **a,** Compocyte modularizes cell type annotation through local classifiers at taxonomy branching points allowing for flexible choice of the most performant classifiers in the sklearn, catboost and pytorch packages. Integrated uncertainty quantification avoids misclassification, stopping classification early at coarser labels. Classifier hierarchies can be interchanged to modify user-defined cell type taxonomies and synthesize knowledge from different sources (classifier synthesis), **b,** Correlation of uncertainty metrics with subjective certainty of rater 1 (R1) and rater 2 (R2) on a 5-point Likert scale or tree-distance between rater labels (inter-rater). Uncertainty metrics include Monte-Carlo dropout (MC), Rao’s Q, and tree distances (tree dist) using their standard deviation (stdev) or mean within metacells (SEACells, left panel). Correlation patterns of uncertainty metrics (right, panel), **c,** A sum of MC and the average tree distance in SEAcells shows the higest correlation with mean tree distance from predicted to ground truth labels, **d,** Local classifier architectures can be exchanged to include neural networks (n=neurons), tree-based models or logistic regression (upper panel) using local classifiers feature spaces (lower panel), **e,** Local model optimization can increase the perfomance of difficult-to-classify cell types such as dendritic cells (DC) or CDS LLE (long lived effector T cells, color code indicating study), **f,** Classifier synthesis allows merging independent classifiers of overlapping sublineages, yielding virtually identical performance as compared to training the classifier on the combined data. hF1: hierarchical F1 score, hR: hierarchical recall, hP: hierarchical precision.

To reduce misclassification during automated annotation, Compocyte employs partial label rejection. Rather than assigning a highly granular but incorrect label, the model halts classification at a predefined certainty threshold to output a broader, biologically precise parent label (**Fig. 3a**, **Methods**). To validate these annotations, we developed an uncertainty quantification metric calibrated against Suco’s ground-truth measurements (subjective rater certainty and hierarchical inter-rater label distances, **Fig. 1c, Methods**). Because cell-level uncertainty metrics often obscure phenotypic consistency, we quantified uncertainty across transcriptomically homogeneous groups of cells (metacells) using the using SEACells algorithm^75^. Within each metacell, we calculated the average ontology tree-distance of the assigned labels. Unlike standard entropy, tree-distance inherently penalizes biologically distant misclassifications and correlated strongly with rater certainty (**Fig. 3b**). Additionally, average tree-distance did not correlate with Monte-Carlo dropout estimates of epistemic model uncertainty, indicating these metrics capture distinct aspects of model stability and label correctness. We therefore integrated normalized Monte-Carlo dropout standard deviations with normalized average tree-distance, which maximized correlation with ground-truth confidence (**Fig. 3c, Methods**).

To demonstrate the practical advantages of this structural flexibility, we highlight two key capabilities enabled by Compocyte’s modularity. First, Compocyte permits targeted retraining of specific subtrees, allowing users to optimize classification by choosing the most adequate model architectures and feature spaces (**Fig. 3d**). This local optimization helps identify phenotypically subtle or data-sparse cell types without disrupting the global architecture (**Fig. 3e**). Furthermore, it enables the synthesis of a single hierarchical classifier from independent models trained on separate lineages (**Fig. 3a**). We validated this synthesis approach by training Compocyte modules independently on published tumor-infiltrating myeloid^61^ and T cell atlases^42^ (**Methods**). We then trained a coarse classifier on the Suco resource to separate major T and myeloid lineages, using it to link the independent lineage-specific models. We evaluated this synthesized classifier on a held-out breast cancer dataset (Bassez et al.^57^). The synthesized model yielded performance virtually equivalent to a classifier trained jointly on all three datasets simultaneously (**Fig. 3f, Table S7**). By facilitating robust performance without relying on global data integration, Compocyte provides a framework for the federated training of broad, cross-lineage classifiers.

### A framework for benchmarking automated hierarchical single cell annotation

Accurately assessing the generalizability of any machine learning classifier requires strict statistical independence between training, validation, and test data^1^. Suco provides such a resource to evaluate single-cell machine learning models with independently processed and annotated datasets. Using Suco, we benchmarked Compocyte against the most frequently utilized cell-type annotation tools across Python and R (**Fig. 4a**, **Methods**). These included variational autoencoder (VAE)-based models (scANVI^12^, scPoli^76^, treeArches^77^), regression-based approaches (CellTypist^78^), and similarity-based classifiers (transferData/Azimuth^32^, CHETAH^79^). To rigorously assess generalizability, we employed a 12-fold hold-one-dataset-out cross-validation strategy, training the models on a subset of 500,000 cells across all but one dataset, which was retained exclusively for hierarchical performance testing (**Methods**). Compocyte achieved the highest overall performance scores despite its modular, hierarchical architecture (**Fig. 4b**). Misclassifications predominantly involved cells where raters assigned broad partial annotations rather than maximally granular ontology terms, indicating that these discrepancies represent genuine aleatoric data ambiguity rather than epistemic model limitations (**Fig. 4c**). Notably, all models except CHETAH demonstrated robust classification performance, supporting the notion that a lack of classifier generalizability is primarily a data limitation rather than a model deficit (**Fig. 4b**). Furthermore, discriminative machine learning models (Compocyte, CellTypist, Seurat) outperformed complex deep generative models (scANVI, scPoli, treeArches) even without prior batch correction (**Fig. 4b**). These findings suggest that learning discriminative boundaries directly on unintegrated expression profiles preserves critical biological variance that generative models may inadvertently remove during latent space alignment, establishing frameworks like Compocyte as superior, integration-free foundations for single-cell annotation.

**Fig. 4.**
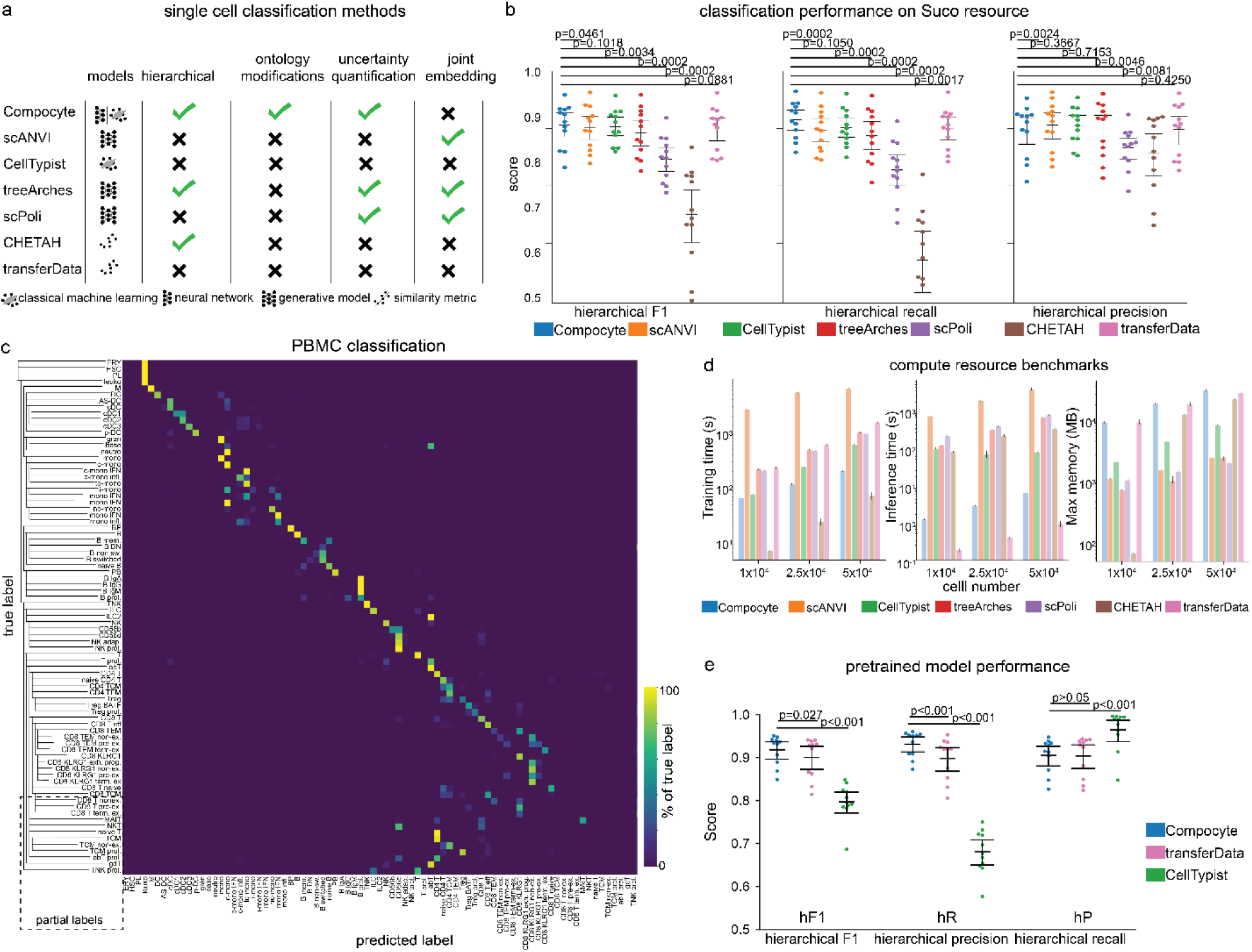
Compocyte outperforms reference mapping methods in immune cell annotations. **a,** Compocyte enables hierarchical classification allowing flexible choice of the most performant classifiers in the sklearn, Catboost and Pytorch packages which can be interchanged to modify user-defined cell type taxonomies with uncertainty quantification avoiding misclassification, **b,** Compocyte shows state-of-the-art performance in a leave-one-dataset out approach on the datasets in Suco PBMC (n= 12, subset of 500,000 cells per run), **c,** Misclassifications generally occur within lineage or with ambiguous cells which are assigned partial labels (meaning non-leaf-node) by independent raters, **d,** Compocyte has ordinary resource requirements and training and inference time as compared to other methods. **e,** Compocyte outperforms available pretrained scRNA-seq cell typing methods.

Our hold-one-dataset-out strategy provides a more realistic assessment of real-world performance than conventional random-split holdouts, which remain the frequent standard in reference atlas construction and benchmarking^9,80-83^. Even in rigorous atlasing initiatives, external testing is often limited to small numbers of unseen datasets or remains susceptible to information leakage inherent to integrated data structures^13,84^. Using Suco, we show how random holdouts inadvertently leak dataset-specific technical artifacts between training and test sets, predictably leading to overestimation of classification capabilities (**Fig. S7**). While scANVI achieved near-perfect accuracy on random holdout data, its performance dropped to mid-range under the stricter hold-one-dataset-out paradigm (**Fig. S7**). By contrast, Compocyte’s performance remained relatively stable (**Fig. S7**), largely due to its resistance to overfitting conferred by high dropout rates^85^, and batch-normalization^86^ at default settings. Furthermore, Compocyte achieved superior recall (**Fig. 4b**) through its partial label rejection mechanism, prioritizing the assignment of a biologically sensible, less granular parent label over risking a forced misclassification at the terminal nodes (**Methods**).

Beyond classification performance, we systematically benchmarked compute resource requirements. Compocyte demonstrated highly competitive training and inference times, surpassed only by CHETAH and transferData, respectively (**Fig. 4d**). Moreover, Compocyte’s native batching strategy maintained moderate, stable memory footprints (**Fig. 4d**). For population-scale applications, Compocyte seamlessly integrates with Dask for out-of-core computation, enabling the processing of datasets containing millions of cells without memory constraints. Leveraging this scalability, we trained a global Compocyte model on a corpus of 5.4 million PBMCs from Suco, to provide a granular highly accurate open-source hierarchical classification model to the research community (10.5281/zenodo.19708295). To ensure an unbiased evaluation of this population-scale model, we tested it against 12 entirely independent external PBMC datasets (**Table S2, S5**). This pretrained Compocyte classifier robustly outperformed the most popular pretrained models from the runners-up CellTypist^78^, and transferData (Azimuth)^32^, particularly in label precision (**Fig. 4e**). Together, these results demonstrate that Compocyte resolves the computational and statistical bottlenecks that previously precluded the application of top-performing, hierarchical classification to population-scale single-cell data.

### A vaccination-induced inflammatory monocyte state is preserved across diseases

In-depth profiling of specific cell lineages in defined contexts by scRNA-seq has led to the discovery of novel inflammation-associated cell types^26,87^. However, limited quantity of the reference data often precludes training accurate cell type classifiers to map these cells to query data containing multiple other cell lineages. Compocyte enables us to dissect this problem by leveraging a large generalizable pretrained model to perform coarse subsetting and using the reference data only to retrain and exchange specific subtype classifiers (**Methods**). We demonstrate the power of this modular approach in inflammatory responses induced by SARS-CoV-2 vaccination in immune checkpoint therapy (ICT) treated patients (**Fig. 5a**). We previously showed that this combined treatment triggers the release of cytokines typically observed in cytokine release syndrome (CRS), a severe adverse event, even if clinical symptoms are absent (subclinical CRS). We further reported that these patients showed prolonged overall survival^88^ – an association later validated by others^89^. However, the cellular sources of inflammatory responses in subclinical CRS remained unclear.

**Fig. 5.**
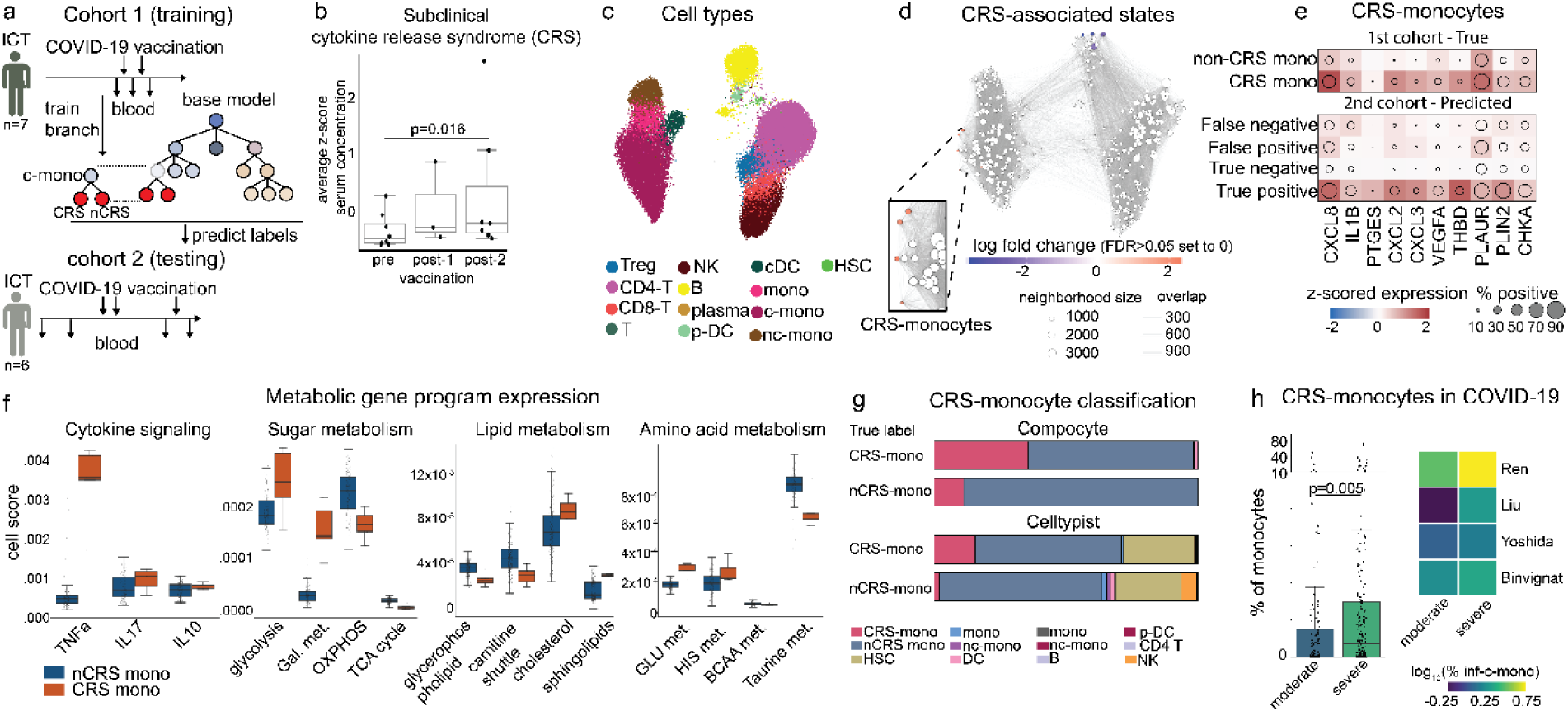
Compocyte identifies subclinical cytokine-release syndrome associated monocytes after vaccination and severe SARS-CoV-2 infection. **a**, We generated peripheral blood mononuclear cell (PBMC) scRNA-seq data before and after SARS-CoV-2 vaccination (Comirnaty, BionTech/Pfizer) from two independent cohorts of cancer patients undergoing immune checkpoint therapy (ICT). Cohort 1 patients (n=7) were sampled before receiving the first, after the first (<6 weeks), and after the second (<6 weeks) Comirnaty dose (n=16 samples). Cohort 2 patients (n=6) were sampled before starting ICT, 1-7 weeks after starting ICT but before Comirnaty, and before and after (<6 weeks) the third Comirnaty dose (n=23). CRS-associated monocytes (CRS-monos) were identified using Milo to train a CRS-mono classifier with cohort 1 which was synthesized with the Compocyte classifier trained on the Suco resource and apiied on Cohort 2. **b,** Average z-scored serum concentration of CRS-associated cytokines (IL-6, CXCL8, IL-2, CCL2, slL1-RA) pre-vaccination (pre), after the first {post-1) and second (post-2) Comirnaty dose with p values calculated using Mann-Whitney U tests (n=17). **c,** UMAP of 35,028 PBMC with color code indicating cell types, **d,** Specific monocyte neighborhoods (CRS-monos) were significantly associated with high CRS-associated cytokine levels as revealed by Milo (false discovery rate (FDR)=0.05). **e,** Dot-plot showing relevant differentially-expressed genes in CRS- and other monocytes, **f,** Mean Spectra program expression in cellular neighborhoods (n=428) of CRS-monos and nCRS-monos. Gal. (galactose), met. (metabolism) **g,** Compocyte shows higher accuracy than Celltypist (without majority voting) in the unseen cohort 2 data (n=15,634 nCRS-monos and n=997 CRS-monos) with stacked bars indicating predicted cell types, **h,** Predicted proportion of CRS-monos of all monocytes per patient sample in patients with mild or moderate (moderate, n=181) vs severe or critical COVID-19 (severe, n=120, left panel). Heatmap showing the proportion of CRS-monos in each study (right panel).

To address this question, we selected 16 samples from 8 patients with subclinical CRS and analyzed their PBMCs before and after SARS-CoV-2 vaccination using the Atrandi Biosciences scRNA-seq platform (cohort 1, **Fig. 5a,b**). We used our Cytopus-based workflow to hierarchically annotate this data (**Fig. 5c**) and performed differential abundance analysis using Milo^90^ (**Fig. 5d**). Milo revealed several monocyte neighborhoods positively correlated with CRS cytokines (**Fig. 5d**, Cxcl7, IL-6, Ccl2, IL-2, sIL2-Ra)^88^. These monocytes showed significantly higher expression of *CXCL8*, *VEGFA*, *THBD* and *PLAUR*^91^, suggesting an inflammatory, pro-angiogenic phenotype (**Fig. 5e**). To identify the cellular processes characterizing these subclinical cytokine release syndrome associated monocytes (CRS-monos) we used Spectra^7^ to identify interpretable gene programs from our scRNA-seq data leveraging our knowledge base Cytopus 2. In line with the differentially regulated genes observed, Spectra revealed a strong transcriptomic program of TNF-α signaling (**Fig. 5f**). Moreover, we found a high degree of metabolic reprogramming compatible with an activated phenotype including higher glycolysis, galactose and cholesterol metabolism but lower tricarboxylic acid (Krebs) cycle activity and reduced oxidative phosphorylation (**Fig. 5f**). Thus, while SARS-CoV-2 vaccination induced angiogenic monocyte phenotypes classically linked to cancer progression^91^, it remains associated with longer survival^89,92^, likely because the transient nature of this inflammatory boost.

We next asked whether Compocyte could map CRS-monos in additional datasets. This task was challenging because our SARS-CoV-2 vaccination dataset (cohort 1) was small and used a different single cell platform (Atrandi Biosciences). We therefore used our cohort 1 only to train a subclassifier dividing classical monocytes (c-monos) into CRS-monos and non-CRS-monos (nCRS-monos) which we then synthesized with our Compocyte base model pretrained on our Suco PBMC resource. Testing this combined classifier on a prospectively collected cohort of 23 samples from 6 ICT patients undergoing SARS-CoV-2 vaccination (**Fig. 5a**, **S8**, cohort 2) we showed an accuracy of 86% for nCRS/CRS-mono classification (**Fig. 5g**, **S8d, Methods**). The predicted CRS-monos in our second cohorts showed identical angiogenic marker expression patterns thus confirming their identity (**Fig. 5e**). Notably, the false-negative cells in cohort 2 lacked these markers, suggesting they represent incorrect annotations in the test set rather than an epistemic model error by Compocyte (**Fig. 5e**). Using the base model virtually eradicated cross-lineage contamination with incorrect annotations almost being almost exclusively within classical monocytes (e.g. nCRS-monos, **Fig. 5g**). By contrast, Celltypist frequently mislabeled CRS-monocytes as hematopoietic stem cells (HSC) or NK cells, resulting in an accuracy of 60% (without majority vote) or 71% (with majority vote) (**Fig. 5g, Fig. S8d**). Importantly, with majority voting Celltypist did not find any CRS-monos likely due to their low frequency (**Fig. S8d**).

Using our classifier we explored CRS-monos across diseases using 7 external PBMC datasets from 389 patients with cancer, autoimmune and acute infectious disease (**Table S2**, **Fig. S8c**). While we found CRS-monos in all these diseases (**Fig. S8c**), we noticed an overwhelming abundance in patients with severe and critical COVID-19 as compared to moderate or mild cases – an effect conserved in all 4 COVID-19 datasets analyzed (**Fig. 5h**). Collectively our results show that Compocyte can accurately annotate rare cell populations across large patient cohorts even with small reference datasets, effectively abrogating cross-lineage misclassification. Furthermore, we establish CRS-monocytes as a novel angiogenic classical monocyte phenotype conserved across diverse pathologies, and associated with adverse outcomes in COVID-19.

### Divergent gene expression programs in human memory T cells across cancer types

The characterization of memory T cell (Tmem) subsets has historically relied on surface protein expression. While scRNA-seq has provided unprecedented transcriptomic breadth, it often fails to resolve specific Tmem phenotypes (**Fig. S9**). Key lineage markers are frequently sparse at the transcript level (e.g., *FAS*) or exist as isoforms (e.g., *CD45RA* vs. *CD45RO*) that standard short-read sequencing cannot distinguish. This resolution gap is particularly acute in human solid tumor metastases, where paired RNA and protein datasets (CITE-seq) remain scarce. Consequently, how metastases rewire gene expression programs within <u>Tmem</u> remains poorly understood.

To address this, we assembled a comprehensive cohort of 39 tumor-infiltrating leukocyte (TIL) samples, totaling 150,059 cells (**Table S2, S8, S9**). This cohort included primary tumors and various metastatic sites from pancreatic (PDAC) and non-small-cell lung cancer (NSCLC) patients (**Fig. 6a**, left). These clinical samples represented a broad demographic (age range 29-90) and seven distinct metastatic sites, obtained using surgical resections (54%) or core needle biopsies (64%) (**Fig. 6b**).To ensure precise separation of conventional Tmem cells from unconventional lineages such as γδT cells, we designed a custom CITE-seq panel (**Table S6**) and performed sample-level manual gating (**Fig. S9d**) which allowed for the identification of subsets that are typically masked on the RNA level (**Fig. S9c,d**).

**Fig. 6.**
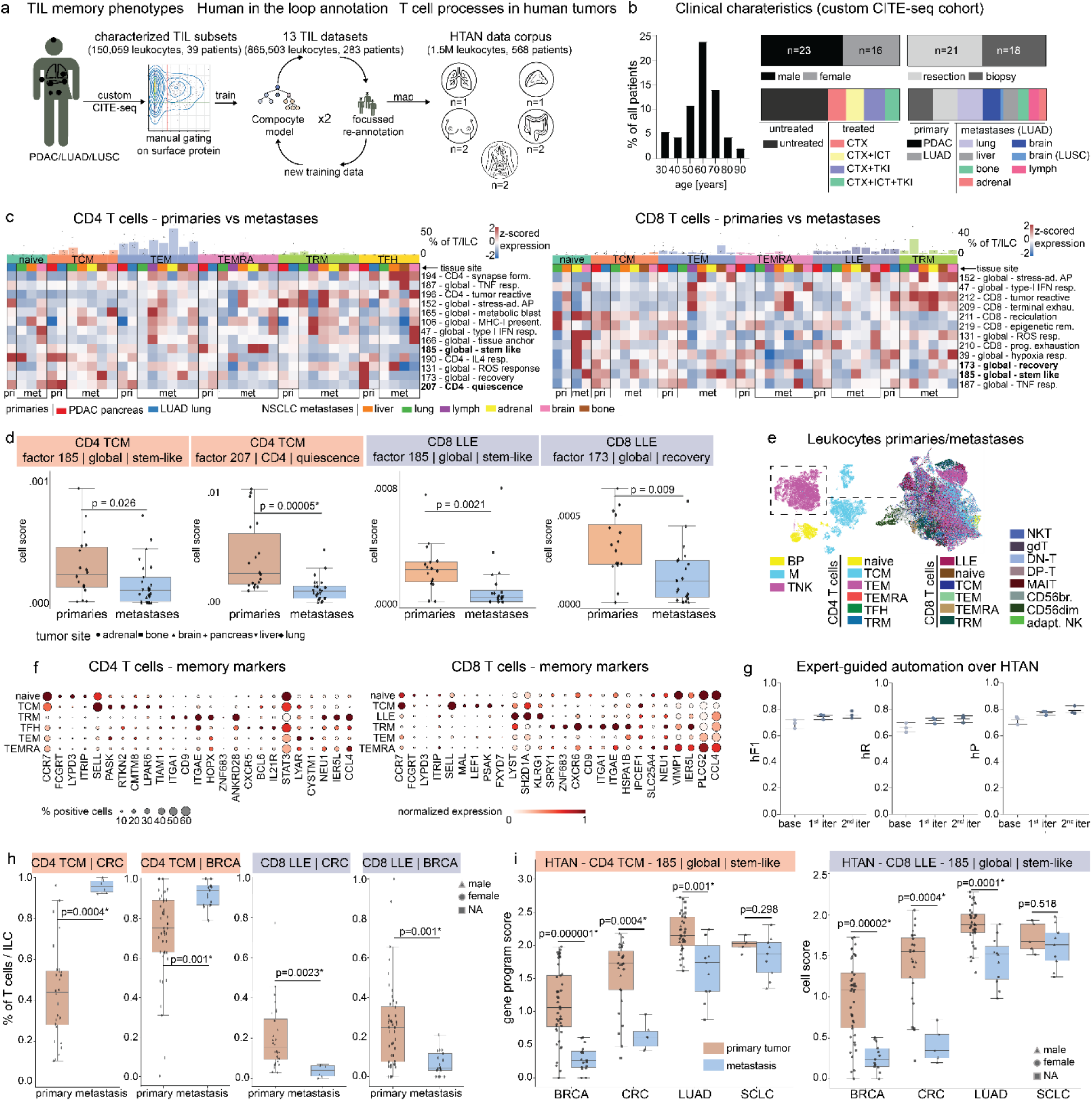
Compocyte reveals metastatic-reprogramming of T memory cells across cancer types. **a,** Primary and metastatic lung adenocarcinomas (LUAD), lung squamous cell carcinomas (LUSC), and pancreatic adenocarcinoma (PDAC) samples were profiled via CITE-seq, with memory T cells identified by surface-marker gating. We further optimized the model by iteratively training Compocyte on independent lung cancer datasets (Salcher et al.), and manually refining the predicted labels. The resulting classifier was applied to the complete HTAN scRNA-seq data release (Nov. 2025). **b,** Clinical characteristics from our CITE-seq samples from n=39 patients. CTX: chemotherapy, ICT: immune checkpoint therapy, TKI: tyrosine kinase inhibitor. **c,** Heatmaps indicating z-scored (over rows) mean cell scores per sample and cell type of differentially expressed Spectra gene programs in metastases (met) and primary tumors (pri) with cell types indicated in the color code on the top and bars indicating cell type proportions of naive T (naive), T central memory (TCM), T effector memory (TEM), T effector memory with CD45RA expression (TEMRA), tissue resident memory (TRM), long-lived effector (LLE), and T follicular helper (TFH) cells **d,** Mean Spectra cell scores for selected gene programs and cell types of n=46 samples from our CITE-seq LUAD, LUSC, PDAC cohort. P values were calculated using Mann-Withney-U tests with signficant comparisons highlighted with an asterisk as determined by the Ben1amini-Hochberg (BH) method **e,** Uniform Manifold Approximation and Pro1ection for Dimension Reduction (UMAP) of n=150,059 tumor-infiltrating leukocytes from n=39 tumors and UMAP of n=84,222 tumor infiltrating T and innate-lymphoid cells from n=39 tumors. **f,** Marker genes for T memory subtypes (Tmem) in our CITE-seq cohort help validating Compocyte’s annotation on RNA alone. **g,** Compocyte classification performance increases with iterations (iter) of human-in-the-loop refinement as indicated in a (middle panel). **h,** Cell type frequencies of Tmem subtypes defined by Compocyte in the HTAN data release from all T / innate lymphoid (ILC) cells in colorectal cancer (CRC) and breast cancer (BRCA) patients. **i,** Average per sample Spectra gene program cell scores in CD4 TCM and CD8 LLE in primary tumors and metastases across BRCA, CRC, LUAD and small-cell lung cancer (SCLC) patients (n=160) with p values obtained using Mann-Whitney-U tests, significant comparisons are indicated with an asterisk as determined by the BH method

We analyzed the abundance of the resulting high-quality Tmem annotations across primary tumors and metastases (**Fig. 6c**). In the CD4 compartment, recirculating populations, specifically T central memory (TCM) and T effector memory (TEM) cells dominated across all tumor sites. Conversely, the CD8 compartment was characterized by tissue-bound effectors, including tissue-resident memory (TRM) cells and KLRG1^+^ long-lived effector (LLE) cells (**Fig. 6c**). To determine if the primary and metastatic environments drive divergent cellular processes within <u>Tmem</u> subsets, we utilized Spectra^7^ for gene program inference. This analysis revealed multiple previously undescribed gene programs in T cells differentially expressed in primary tumors and metastases (**Fig. 6c, Supplementary Note**).

We observed a profound transcriptional shift, particularly regarding cellular stress, quiescence, and stemness. In CD4 TCM and TEM cells, primary tumors showed significantly higher expression of stem-like (factor (F) 185) and quiescence (F207) programs (**Fig. 6d, Fig. S10a**). While both programs share core survival factors such as *IL7R*, *JUNB*, and *CREM*, they represent distinct regulatory modes. The stem-like program (F185) utilizes potent mRNA decay and signaling regulators (*ZFP36^93^, ZFP36L2, TNFAIP3^94^, PDE4B^95^*) to enforce a resting, anti-inflammatory state (**Table S10**). In contrast, the quiescence program (F207) enforces suppression through limiting mTOR activation (*PIK3IP1^96^*), anti-apoptotic mediators (*RTKN2^97^*), and co-inhibitory receptors like *TIGIT^98^* (**Table S10**). The CD8 compartment mirrored these changes (**Table S10**): Primary tumor-infiltrating CD8 LLEs and TRM were highly enriched for the F185 stem-like program and a unique tolerogenic recovery program (F173, **Fig. 6d**). The latter is characterized by genomic and ER stress responses (*TP53^99^, EDEM2^100^*) and TGF-β signaling (*SMAD2^101^*). In metastases, CD8 LLEs and TRM adapted to an apparently hostile metastatic niche, showing higher terminal exhaustion (F209), stress-adapted activation (F152), and type I interferon signaling (F47, **Fig. S10**).

### Erosion of conserved memory T cell programs across solid tumors metastases

To validate these findings at scale, we refined our Compocyte TIL classifier using a human-in-the-loop strategy which integrates manual feedback into automated annotation to allow for continuous ontology refinement (**Fig. 6a**, middle panel).

We utilized a lung cancer meta-atlas comprising 865,503 leukocytes from 283 patients across 13 studies. By dividing this data into independent training and test sets, we iteratively refined the model using our uncertainty quantification framework: Compocyte was applied to the data followed by manual correction of the most uncertain annotations (**Methods**). To facilitate manual validation, we defined new biologically plausible RNA markers for Tmem subtypes using our entropy-based marker gene identification method described above (**Fig. 6e,f, Methods**). This human-in-the-loop strategy steadily improved hierarchical precision and recall (**Fig. 6g**), with the model converging toward maximum performance at high annotation granularity. Finally, we deployed this validated classifier across the entire HTAN data corpus, encompassing 1.5 million tumor infiltrating leukocytes (TIL) from 568 patients with breast, colorectal, and lung cancers and other tumors (**Fig. 6a**, right panel). The resulting dataset enabled us to explore Tmem distribution and gene programs across tumor types. We found that while CD4 TCM populations numerically increased, CD8 LLE numbers decreased in breast and colorectal cancer metastases (**Fig. 6h, S10b**). Across multiple cancer types, metastatic Tmem gene expression programs appeared severely altered, thus confirming the results from our NSCLC and PDAC cohort. Metastatic CD4 TCM and CD8 LLE exhibited significantly lower expression of the stem-like (F185), quiescence (F207), and tolerogenic recovery (F173) programs compared to their primary tumor counterparts (**Fig. 6h, Fig. S10c**). Because these programs were initially inferred from our NSCLC and PDAC cohort, their consistent downregulation across independent pan-cancer datasets suggests a universal functional shift. Rather than acquiring a single, unified metastatic program, T cells in the metastatic niche appear to lose the highly conserved stem and quiescence programs that maintain stable memory states in primary tumors. This suggests that T cell self-renewal is fundamentally impaired in metastatic microenvironments across cancer types.

## Discussion

The development of single-cell transcriptomics machine learning methods has historically been limited by the absence of granular, uniform, and independently annotated benchmarks^4^. The single cell universal classification omnibus (Suco) overcomes this hurdle by providing ground-truth for any type of single cell transcriptome representation learning. Rather than simply engineering more complex generative algorithms for data integration, our benchmarking demonstrates that high-resolution, consistently labeled training data remains the fundamental prerequisite for accurate cell type classification. By establishing this baseline, Suco shifts the field’s focus from algorithmic architecture toward data fidelity.

Suco enabled us to develop Compocyte providing a blueprint for developing future cell type classifiers. Compocyte resolves the computational and biological rigidities of reference mapping. By using ontology branching points to modularize annotations, Compocyte focuses and expedites model retraining and enables greater modifications than typical transfer learning architectures^18^. Modularizing training tasks also circumvents challenges in continuous learning such as catastrophic forgetting^19^ and model instabilities which even with sophisticated continual learning architectures can only be mitigated^14,102^. Combined with uncertainty quantification that facilitates human-in-the-loop expert review, this approach transitions population-scale annotation from a static, computationally-prohibitive task into an iterative, biologically faithful framework capable of mapping global cohorts.

Applying this framework at scale exposes previously unappreciated but highly conserved immune cell types. In the tumor microenvironment, the emergence of a conserved, tumor-associated resorptive macrophage (TARM) phenotype across diverse solid tumors indicates that neoplastic tissues systematically co-opt physiological, osteoclast-like invasive programs independent of the bone marrow niche. This convergence suggests that tissue invasion and metastasis rely on the selective enrichment of highly specific tissue-remodeling myeloid states. Importantly, the expression of *ACE* within this population exposes actionable therapeutic vulnerabilities and a mechanistic rationale for the retrospective clinical benefits associated with ACE-inhibitors in pancreatic^103^ and colorectal cancer^104^ where TARM were particularly abundant. Mechanistically, the ACE product angiotensin II is a well-established driver of cancer cell invasion and metastasis in vivo^105^. Because of their RANK expression, TARM present a compelling therapeutic target for the clinical antibody Denosumab in breast cancer^73^.

Beyond the tumor bed, circulating monocytes exhibited previously uncharacterized adaptations to therapeutic perturbations. The subclinical cytokine-release syndrome-associated monocyte (CRS-mono) is defined by an inflammatory cytokine signature (CXCL2, *CXCL3*, *CXCL8, IL1B*), pro-angiogenic (*VEGFA*, *PLAUR*^91^) mediators, and distinct metabolic rewiring with upregulated glycolysis, galactose metabolism, and cholesterol synthesis. This adaptation, likely required to sustain acute pro-inflammatory secretion profiles, is diametrically opposed to the tolerogenic states induced by influenza vaccines^106^. Because acute systemic inflammation can reinforce anti-tumor immunity via IL-1b^107^ or by CXCR1/2 ligands^108^, this specific myeloid polarization offers a compelling biological mechanism for the temporally enhanced efficacy of immune checkpoint therapies observed following specific vaccinations^88,89,109^.

Finally, projecting high-resolution cellular ontologies across the pan-cancer Human Tumor Atlas Network (HTAN) cohort revealed the erosion of core regulatory programs governing stemness and quiescence as the dominant, metastasis-induced features conserved across tumor types. We hypothesize that this programmatic loss is driven by a time-dependent failure of niche formation. Primary tumors often co-evolve with the immune system over years, forming complex immune architectures such as tertiary lymphoid structures. In contrast, the rapid expansion of metastatic lesions precludes the development of these slow-building, supportive niches often reflected in the emergence of immature tertiary lymphoid structures^110-112^. Consequently, this functional collapse occurs because the rapidly evolving metastatic environment lacks time and structural stability needed to sustain memory T cell stemness and withstand harsh microenvironments.

In summary, Suco and Compocyte establish a unified machine-learning framework that functionally maps the immune system at an unprecedented scale. By providing a trusted benchmark, Suco enables building the next generation of artificial intelligence methods for single cell genomics. While Compocyte and Suco are currently limited to supervised training and human leukocytes, modularity enables their iterative expansion and inclusion of other lineages. Moving beyond static data integration with an architecture built for federated refinement, this approach facilitates the systematic discovery of systemic and microenvironmental principles governing human immunity.

## Methods

### Clinical cohorts overview

#### Generating CITE-seq data for tumor infiltrating leukocytes

##### Single cell isolation & cryopreservation

Human lung and pancreatic tissue samples were obtained by core biopsy or surgical resection at the Memorial Sloan Kettering Cancer Center (MSKCC, U.S.A.) from patients who provided written informed consent under an institutional review board approved protocol (14-091) and following the Declaration of Helsinki in its current form. Tissue was processed within 0.5h after resection and transferred to the sample processing laboratory on ice. Single cell suspensions were obtained following a validated standard protocol^113,114^. Briefly, samples were digested using the Miltenyi GentleMACS (Miltenyi Biotech, Germany) system, followed by red blood cell lysis, sorted for CD45 positive and live cells by flow cytometry (FACS, BD Biosciences, NJ, U.S.A.) or magnetic cell sorting (Levicell, LevitasBio, CA, U.S.A.) and cryopreserved using Bambanker freezing media until further analysis (Nippon Genetics, Japan). We performed side-by-side comparisons to determine the effects of cell sorting before or after freezing on cell type composition, library size and complexity, were unable to find relevant differences, and sorted most samples before freezing.

##### CITE-seq data generation

We screened 278 oligonucleotide-tagged antibodies (Totalseq-A, Revvity, U.S.A.) to identify a custom CITE-seq panel to characterize tumor-infiltrating leukocytes across metastatic and primary tumors in pancreatic and lung cancer patients (**Table S6**). Antibodies were assessed for specific staining in cell type populations of interest, their aberrant staining and additional information for cell typing. Using these criteria we arrived at a custom set of 169 antibodies (**Table S4**) which included 7 antibodies not contained in standard commercial CITEseq panels (TotalSeq-A Universal Cocktails v1.0, Revvity, U.S.A.). The additional antibodies allowed for more granular distinction of myeloid (CD15, CD66b, c-kit) and unconventional T cell subsets (GITR, TCR-γ/δ, TCR-Vγ9, TCR Vα24-Jα18). Cryopreserved tumor-infiltrating leukocytes were thawed and stained using the CITE-seq panel above. Samples were multiplexed using Totalseq A hashtag antibodies directed against β2-microglobulin and CD298 (Revvity, U.S.A.). Samples were carried forward if cell viability was >70% and were otherwise discarded. Samples were then processed using the 10x Genomics Chromium 3’ sequencing v3.0 or v3.1 Kit (10X Genomics, U.S.A.). We made side-to-side comparison of the v3.0 and v3.1 kits and did not see any relevant changes in cell type distribution. The manufacturer’s protocol (user guide CG000315, 10x Genomics) was modified in that separate sequencing libraries were generated for gene expression (>300bp) and antibody-derived tags (<180bp) by separating them with SPRI select beads (BeckmanCoulter, U.S.A.) from the obtained cDNA. Sequencing was performed on Illumina NovaSeq S4 or NovaSeq X Plus sequencers at 150 base pairs paired-end reads.

##### CITE-seq specific preprocessing - RNA

The gene expression library derived FASTQ files were aligned using SEQC^115^ and the GRCh38-2020-A human genome reference. Standard SEQC quality control thresholds were applied including removal of cells with >20% mitochondrial gene content. The output data was processed using the scanpy (v1.11.1) and AnnData (v0.11.4) software packages. We filtered out low quality cells by filtering out cells with less than 200 genes and genes with less than 10 cells. Mitochondrial and ribosomal genes were removed from downstream analyses since these are heavily driven by technical effects. The data was preprocessed in two independent batches and unified after subsetting the leukocyte population.

To remove residual empty droplets, we applied EmptyDrops with a reference droplet population containing less than 30 genes (lower=30) and standard parameters (retain=Inf, test.ambient=T, alpha=0.05). This algorithm identified droplets containing genuine cellular profiles through their statistically significant divergence from a reference of empty droplets containing only ambient RNA. To validate that only high-quality cells were retained, we confirmed that PhenoGraph derived cell type clusters (k=10) preserved structural gene-gene covariance patterns.

We then removed doublets using the DoubletDetection package (v4.2) per sample (10.5281/zenodo.2658729) using the PhenoGraph algorithm (verbose=True, standard_scaling=False), a p value threshold of 1e^-7^, and a voter threshold of 0.8. We verified that all flagged doublets with a false-discovery rate (FDR) lower-bounded by the number of algorithmic iterations had an FDR<0.05. We reasoned that doublets show similar transcriptomic patterns as compared to singlets. We therefore used PhenoGraph clustering (k=10) to identify and remove entire clusters showing significantly higher numbers (>2 fold) of doublets as expected by uniform distribution using Chi-square test corrected for multiple comparisons using the Benjamini-Hochberg (BH) method (α=0.05). We chose high cluster resolution (k=10) which reduced the number of cells not flagged as doublets by DoubletDetection to avoid discarding genuine high-quality cells. Moreover, we removed all residual doublet calls. We also removed clusters showing conflicting lineage signatures (for example, concurrent B-lymphocyte alongside T-lymphocyte markers) representing heterotypic multiplets.

Gene expression was normalized to median library size and log1p transformed (pseudocount 1, natural logarithm) and then iteratively annotated: To focus on the most informative genes we took the top 10,000 highly variable genes identified using the scanpy implementation of the Seurat v3 method and added a custom list of cell type specific genes of interest (**Table S11**) to our highly variable (hv) genes. We selected an optimal number of principal components (PCs) as the lowest number explaining at least 25% of the total variance, or at least 50 PCs whichever number was higher. We clustered the data using PhenoGraph. To select an optimal k parameter we calculated clusterings for different parameters (k=10,20,30,40,50,60,70,80,90,100) and compared their pairwise rand-indices as well as their ability to distinguish small cell type populations (e.g. innate lymphoid cells, dendritic cells). We then set k as low as possible while still achieving robust clustering with rand indices in the neighboring parameters >0.8. Clusters were annotated using curated lists of marker genes included in our Cytopus 2 resource (https://github.com/wallet-maker/cytopus) and cell types of interest were carried forward for further subtyping by repeating the steps 1 to 4 above.

We first clustered the data [k=40 (first cohort), k=30 (second cohort)] to dissect coarse cell types (endothelial cells, (myo)fibroblasts/mesenchymal cells, epithelial cells including cancer cells, neuroendocrine cells, neuronal/glial and other cells, and immune cells). We carried forward 150,059 immune cells followed by scran normalization and another round of clustering (k=40, 5000 hv genes) to dissect coarse immune cell subsets (T/NK cells:‘TNK’, B/plasma cells: ‘PB’, myeloid cells: ‘M’). We next annotated the cells using the CITE-seq based gating strategy outlined below. We ran the Spectra factor analysis method on 55,850 TNK cells to obtain gene programs for cellular processes. We set 20 cell type specific factors for each CD8 T cells, CD4 T cells and NK cells, and 189 global factors as a number close to the 199 input gene sets obtained from our Cytopus knowledge base (https://github.com/wallet-maker/cytopus v2.1). We chose a lambda parameter of 0.01 which keeps the factors close enough to the input gene sets to allow for good interpretability, while also enabling discovery of new genes in known gene programs as well as overall new gene programs. T cells were renormalized, highly variables were computed using the scanpy.pp.highly_variable_genes function (with the Seurat method and min_disp=0.01, min_mean=0.001) and clustered using PhenoGraph with a k parameter of 30 followed by manual annotation of the resulting clusters using our Cytopus 2 cell type identity gene sets and removal of Treg cells before all downstream analyses.

##### CITE-seq specific preprocessing – antibody-derived tags

We aligned antibody-derived tags (ADTs) using CITE-seq-count (https://github.com/hoohm/cite-seq-count, v1.4.0). We next normalized the ADT using DSB (with empty_counts_range=(1,2) per sample and with all isotype controls in our CITE-seq panel (**Table S6**). DSB accounts for both cell and sample-specific noise in CITEseq data^116^. We observed in our data that DSB allowed us to optimally distinguish immune cell populations as compared to other normalization techniques (**Fig. S11**). We then gated immune cell types in each sample individually using FlowKit (v1.3.2) and the gating strategy outlined in **Fig. S9d**. Gating thresholds were set for each sample individually by inspecting biaxial density and contour plots and isotype controls so that generally

<1% positive cells were classified as positive in our isotype controls. Methodological reporting aligns with the BRISQ (Biospecimen Reporting for Improved Study Quality) guidelines.

### Generating PBMC scRNAseq data

For the first cohort, blood from patients in the ANTICIPATE trial exploration cohort^88^ undergoing immune checkpoint therapy was obtained before (≤6 months) and after the first or second COVID-19 vaccination (≤6 weeks) via venipuncture or central port catheter. PBMC were isolated by density gradient centrifugation by layering them on top of isotonic polysucrose solution (1.077g/ml) followed by careful centrifugation at 400g for 30min. PBMC were cryopreserved in fetal calf serum containing 10% dimethyl sulfoxide in liquid nitrogen after a cool down of -1°C/min to -80°C. Written informed consent was obtained from all participants before any sample collection or data analysis. All clinical studies have been performed under the Declaration of Helsinki in its current form. Ethics approval was obtained from the Medical Faculty institutional review board at Heidelberg University (Ethikkomission I, S-373/2020). All PBMC samples were processed and cryopreserved <24h after blood sampling. After thawing all biospecimens were directly processed for scRNA-seq.

For the first cohort, single cell genomics data was generated from 5000 cells per sample using the DropletGenomics inDrop platform (now Atrandi Biosciences) following the manufacturer’s instructions and sequenced on Illumina NextSeq 550 read 1 (16 cycles), read 2 (62 cycles). FASTQ files were demultiplexed and aligned using the Solo-in-drops NextFlow pipeline (v1.0, https://github.com/jsimonas/solo-in-drops). We filtered out low quality cells by filtering out cells with less than 200 genes and genes with less than 10 cells. Moreover, we removed mitochondrial and ribosomal genes from downstream analyses. Empty droplets and doublets were removed as indicated above and 35,329 singlets were carried forward to downstream analysis. We selected all genes because highly variable gene selection did not improve our resolution to identify granular cell types. We clustered the data using PhenoGraph and a k parameter of 20 defined by the parameter optimization strategy outlined above.

For the second cohort of 6 patients we obtained PBMC samples before as well as 1 to 7 weeks after starting of immune checkpoint therapy, as well as samples obtained before (<6 months) or after (≤6 weeks) the third vaccination dose (booster vaccination). Samples were stained with Totalseq B (Revvity, U.S.A.) oligonucleotide-conjugated (‘hashtag’) antibodies directed against β2-microglobulin and CD298 for 30min on ice followed by thorough washing of unbound antibody and processing using the 10x Genomics Chromium 3’ v4 single cell gene expression kit with feature barcoding following manufacturer’s instructions (user guide CG000732). Gene expression and cell surface protein (‘hashtag’) libraries were sequenced together on a Novaseq X 25B flow cell (Illumina, U.S.A.). We filtered out low quality cells by excluding cells with less than 200 genes and genes with less than 10 cells. Moreover, we removed mitochondrial and ribosomal genes from downstream analyses. Cell types were identified by clustering using Cytopus marker genes as described for cohort 1.

#### Milo differential abundance analysis

To evaluate shifts in the cellular composition of peripheral blood mononuclear cells (PBMCs) associated with cytokine release syndrome (CRS) markers, we employed the Milo framework. Rather than comparing transcriptomic profiles, Milo detects variations in cellular abundance by evaluating cell density within a k-nearest neighbor (KNN) similarity graph. Following standard protocols, 10% of the cells were sampled as index points to establish representative, partially overlapping neighborhoods. Abundance differences were determined by quantifying the cells from each sample within these neighborhoods and fitting the data to a negative binomial generalized linear model. To correct for multiple testing, spatial false discovery rates (FDR) were calculated.

For our downstream analysis, independent KNN graphs were generated for each patient cohort using a neighborhood size of k = 30. To quantify population shifts during PD-1 blockade, we applied the regression model y_ns_ ∼ cytokines, where y_ns_ denotes the cell count for sample s in neighborhood n. The cytokines variable represents the mean z-scored serum levels of a specific marker panel (IL-6, CCL2, CXCL8, IL-2, and sIL-1RA). Neighborhoods significantly enriched in non-responding patients were isolated using a spatial FDR threshold of < 0.05 containing over 50% classical monocytes were defined as cytokine release syndrome induced monocytes (CRS-monos). All classical monocyte neighborhoods with an FDR ≥0.05 were defined as nCRS-monos. We next compared the mean Spectra factor expression per neighborhood between CRS-mono and nCRS-mono neighborhoods. We also used the cells contained in these neighborhoods in cohort 1 to train a Compocyte classifiers differentiating between CRS-monos and nCRS-monos which we appended to the classical monocyte node in our Compocyte base model trained on Suco. Because Celltypist (v1.7.1) does not allow for hierarchical labels, we trained it with all labels from cohort 1 using default parameters either with or without majority voting.

### Measuring serum cytokine concentrations

Serum samples of patients from the ANTICIPATE trial with available scRNA-seq PBMC data (see above) were obtained at the same timepoints as the PBMC samples using collection tubes equipped with a coagulation matrix (Sarstedt, cat. no. 01.1602). Prior to processing, all blood was held at ambient temperature for a maximum of 24 hours, although most samples were processed within 6 hours. Serum isolation took place at the NCT Liquidbank under established standard operating procedures. Briefly, the blood tubes were centrifuged at room temperature for 10 minutes at 2,500×g. The resulting serum supernatant was then separated into 500-µL aliquots and cryopreserved at -80 °C.

For multiplex quantification, cryopreserved sera were thawed and assayed in duplicate utilizing BioLegend’s Legendplex Cytokine Storm Panel 1 (741091; RRID: AB_2895549) and Panel 2 (741142; RRID: AB_2895550) following the manufacturer’s protocols. Samples were diluted 1:2 for the cytokine multiplex arrays in 96-well V-bottom microtiter plates by combining the diluted sera, standards, assay buffer, matrix, capture beads, and biotin-conjugated detection antibodies. Following the required incubation and wash cycles, streptavidin-PE was introduced. The samples were subsequently transferred to flow tubes for data acquisition on a BD FACS Canto II flow cytometer running the BD FACS DIVA software (v.8.0). Final analyte concentrations were derived from standard curves fitted with a five-parameter logistic regression model via the Legendplex Data Analysis Software (v.2021.07.01). Any measurements falling beneath the assay’s lower limit of detection were recorded as zero.

### General statistics

To compare between patient groups, we generally used non-parametric tests (Mann-Whitney-U test for unpaired, Wilcoxon matched-pairs-signed-rank-test for paired samples) and corrected for multiple comparisons unless indicated otherwise. The alpha for all tests was set to 0.05 within each experiment. Specifics on statistical testing are indicated for each individual analysis in the methods section. For the here-described exploratory endpoints no sample size estimation was performed. Sample-size estimations for the primary endpoint of the ANTICIPATE trial are described in the study protocol (German Clinical Trials Register, DRKS00022890).

### Cytopus 2 Overview

Cytopus 2 builds on our previously introduced single cell gene program knowledge base Cytopus^7^ a lightweight single cell genomics knowledge base built on the NetworkX python package. Cytopus models cell type hierarchies as a graph in the *KnowledgeBase* class. Attached to these cell types are gene sets representative of their *cellular identities* as compared to their sibling cell types in the graph and gene sets representative of *cellular processes* which can for example be used for the gene program inference method Spectra^7^. These gene sets can be retrieved by the *cytopus.KnowledgeBase.processes* and *cytopus.KnowledgeBase.identities* methods, respectively.

In Cytopus 2 we greatly expand the number of identities gene sets to include 164 gene sets indicative of cellular identities (*positive identities*) and 164 gene sets precluding the cellular identity (*negative identities*). Cytopus 2 thereby aligns more with our current semantic models that the presence and the absence of certain molecules are required to establish the identity of a cell type. E.g. for a full phenotype non-exhausted cytotoxic effector T cell phenotype a T cell must acquire *CD8A*, *CD8B* and cytotoxic marker expression such as *PRF1* and *GZMB*. However, it must not express *CD4* or markers of exhaustion such as *TOX*. Within cellular identities, we distinguish cell types which represent a single established entity with representative cellular identity gene sets and cell states, differentiations of cell types which might occur similarly across cell types. While these cell states might alternatively be modeled by gene programs, we believe that depending on the use case, discretization is possible and helpful for gene programs with almost bimodal expression patterns. For example, proliferation represents a strong transcriptional signal which in clustering analysis usually results in a separate cluster. To reduce complexity of the KnowledgeBase object we chose to not separately model these states from cell types but rather indicate the state by separating it from the cell type with an underscore (‘_’) in the name of the cell type.

Cytopus 2 also includes tools to hierarchically query single cell data to facilitate manual annotation and working with cell type ontologies. The *cytopus.Hierarchy* class is used to link cell barcodes to the cell type graph interfacing with the popular anndata^23^ object for single cell data analysis. This helps ensure label consistency and to efficiently label and query cell types in a hierarchical manner. For example, using Cytopus 2 all CD4 T cells (‘CD4-T’) can be collected, which will retrieve all downstream cell types in the ontology for example T regulatory cells (‘Treg’), CD4 central memory T cells (‘CD4-TCM’) or T follicular helper cells (‘TFH’). It also allows quickly analyzing cell type composition at different granularity ensuring a complete subdivision of the data. The hierarchical graph-based approach also ensures that labels show internal consistency. For example, a T follicular helper cell label, a subtype of CD4 T cells, cannot be co-assigned with a CD8 T cell label.

Cytopus 2 interfaces with Compocyte enabling efficient training of hierarchical cell type classification models. For example, training data can be labeled using the *cellular identity* gene sets, attached to the *hierarchy* object and passed to *Compocyte* to train the cell type classifier. Like Compocyte, Cytopus 2 is meant as a lightweight tool which can quickly evolve and be customized to the user’s needs by building or modifying *KnowledgeBase* objects rather than providing a consensus cellular taxonomy. This directly addresses the quickly evolving and ever-increasing granularity of cellular identities uncovered by large-scale single cell genomics data.

### Single cell universal classification omnibus (Suco)

#### Data selection

For the peripheral blood mononuclear cell (PBMC) resource, we selected a total of 12 high quality PBMC datasets spanning 5.39x10^6^ cells from 597 individuals including healthy donors, COVID-19 patients, and cancer patients (colorectal, pancreatic, biliary, breast, **Table S2**). For our tumor infiltrating leukocyte (TIL) resource, we selected 8 datasets totaling 766,000 cells from 179 patients with breast, pancreatic or colorectal cancer (**Table S2)**. Both the PBMC^21^ and TIL^20^ Suco resource are available to the research community through Zenodo (10.5281/zenodo.15350417 and 10.5281/zenodo.13709549, respectively).

For the curation of our core Suco resource we re-processed and re-labelled published datasets. Authors provided aligned count data in h5/h5ad format or as count matrices with sample and feature labels as well as metadata provided separately. To enable better separation of cell type identities by improving signal-to-noise ratio we removed mitochondrial, as well as ribosomal and *PRPS* genes. We also removed cells with less than 200 unique genes and genes that were expressed in fewer than 2 cells. We then normalized gene expression to the median across cells and log1p transformed (natural logarithm, pseudocount=1) our data before running PhenoGraph^117^ with k=10, re-running doublet detection using the python package DoubletDetection (v4.2, PhenoGraph clustering algorithm probability limit below 10^-16^, voting threshold of 0.5; http://doi.org/10.5281/zenodo.2678041) and removing single doublets as well as Phenograph clusters containing 5 times more doublets or ambiguous cells compared to the average across all clusters. We generally set gene expression to raw counts again and re-normalized the data using scran^118^ followed by log1p transformation (natural logarithm, pseudocount=1). Due to scran’s increased memory use at scale, we were unable to run the algorithm for larger datasets (**Table S3**) and in these cases used total-count normalization (to median library size) and log1p-transformation (natural logarithm, pseudocount=1) instead which performs favorably on most datasets^119^.

#### Cell type annotation

We iteratively clustered the data using PhenoGraph or Leiden clustering for larger datasets due to its improved scalability as indicated above in principal component space using the parameters outlined in **Table S2**. We first annotated all leukocytes jointly to identify coarse lineages. We then divided the data into myeloid, B/plasma cell lineage and T/innate lymphoid cell lineage subsets and repeated the normalization, highly variable gene calculation, principal component analysis and clustering as indicated above. Cell type annotation was performed by two investigators independently using the cell type identity genes from Cytopus 2. Raters also documented the positive and negative marker genes supporting their annotations (**Table S3, S4**). Disagreements in rater labels were jointly reviewed and harmonized.

To semi-quantitatively assess subjective rater certainty we introduce a 5-point Likert scale of annotation certainty:

*How certain are you about your annotation?*

*1: very uncertain 2: rather uncertain 3: somewhat certain 4: rather certain 5: absolutely certain*

To quantify differences in rater certainty, we limited our analysis to clusters that had certainty scores assigned by both raters (78%) and which had not been labelled as doublets or low-quality cells. We then calculated the tree distance between rater labels for each cluster using our pre-defined Cytopus hierarchy and NetworkX’ shortest_path function. This function yields a list of all nodes in the path between two labels. This includes both labels, so we subtracted 1 to calculate the number of steps/edges between the two labels.

#### Dataset-specific information

Terekhova et al^35^ (PBMC) provided preprocessed count data in h5ad format filtered to cells with ≥ 512 genes detected per cell, ≤ 6 % mitochondrial counts detected per cell, ≤ 6,654 UMIs and ≥ 1,024 UMIs per cell. Data were aligned by the original study’s authors using Cell Ranger v7.0.0 and the GRCh38 reference genome. Doublets had been removed with Scrublet^120^ and by manually removing clusters expressing markers from different major cell type lineages. Moreover, red blood cells and platelets had been removed. Given this preprocessing, we did not rerun doubletdetection on this dataset.

Ren et al^31^ (PBMC) provided sample identifiers, sample metadata, preprocessed count data in .mtx format, as well as tabular barcodes, features, which was already filtered to include cells with between 500 and 5,000 genes per cell, ≤10 % mitochondrial gene count, and between 1000 UMIs and 25,000 UMIs per cell. The original study’s authors had aligned reads to the GRCh38 genome reference using kallisto/bustools^121^ and had removed doublets with DoubletDetection and Scrublet^120^. Given this preprocessing, we did not DoubletDetection or EmptyDrops on this data. After initial annotation of the T cell and ILC subset we found a relevant number of clusters displaying mixed T cell and NK cell expression patterns. We therefore re-clustered these populations allowed for the separation of CD8 T cell and NK cell substypes (**Table S3**).

Oelen et al^33^ (PBMC) also provided preprocessed count data in the form of an .mtx file, as well as tabular data for features, barcodes, sample identifiers and sample metadata. The study’s authors had aligned reads to the hg19 genome reference using Cell Ranger v3.0.2. Data had been filtered to include cells with ≥ 200 genes, ≤ 9 UMIs registering to erythroid cell marker HBB and ≤ 8/15 % mitochondrial gene count for samples run with 10x Genomics Chromium 3’ single cell gene expression kits v2 and v3 respectively. The authors had also removed doublets in this data with Souporcell^122^. However, we observed large batch effects between cells captured using 10x Chromium v2 compared with v3 reagents, so we separated the data into two groups labelled E1 and E2. We further applied DoubletDetection as outlined above separately to E1 and E2.

Liu et al^30^ (PBMC) provided preprocessed count data in the form of an .rds file filtered to cells with ≥ 200 genes detected per cell, ≤ 4,000 genes detected per cell, ≤ 30 % mitochondrial reads, ≤ 15,000 cell surface protein tags or mRNA reads and ≤ 5,000 hashtag antibody counts. The original study’s authors had aligned reads to an unspecified reference genome using STAR (https://github.com/alexdobin/STAR). The authors had also removed doublets based on demuxlet^123^ output. We converted the .rds file to .h5ad using Seurat and then proceeded as outlined above including DoubletDetection.

Zhang et al^26^ (PBMC) provided preprocessed count data in the form of an .mtx file, as well as tabular data for feature, sample identifiers and sample metadata. We used these to create an AnnData object. After alignment to the GRCh38 reference genome using Cell Ranger v3.1.0, the original study’s authors had filtered out genes present in less than 10 cells and restricted cells to those containing ≥ 400 genes and ≤ 8,000 genes, < 10 % mitochondrial gene count, ≤ 120,000 UMIs and ≥ 600 UMIs per cell. He authors had also removed doublets with Scrublet^120^. Because our initial data analysis showed evidence for residual doublets, we reran DoubletDetection as outlined above which revealed some residual doublets we removed.

Keenan et al^27^ (PBMC) provided count data in the form of an .h5ad file. The original study’s authors had aligned reads to the GRCh38 reference genome using Cell Ranger v3.1.0. The authors had retained cells expressing between 100 and 2500 genes per cell, and < 20 % mitochondrial genes. The authors had also removed red blood cells and platelets. Although they reported having removed doublets, the doublet detection algorithm was not specified.

Hao et al^32^ (PBMC) provided preprocessed count data in .h5seurat format which included only cells expressing between 500 and 6000 genes per cell, and between 10,000 and 50,000 ADT reads. The original study’s authors had aligned reads to the GRCh38 reference using Cell Ranger v3.1.0. They had also removed doublets using hashtag antibodies and HTODemux^124^ and in a second step, removed all cells with ≥ 20 % doublet neighbors after KNN graph calculation on ADT as marked by HTODemux^124^. We converted data to an .h5ad file using Seurat and attached the metadata provided by the authors. We re-applied DoubletDetection as outlined above. We separated the data into two groups sequenced on different lanes labelled E1 and E2 and processed them separately, because they showed strong batch effects.

For each sample, Steele et al^34^ (PBMC, TIL) provided pre-processed count data in .mtx and .csv files containing barcodes, features, and sample metadata. The original study’s authors had aligned reads to the hg19 reference genome using Cell Ranger. The authors had retained only cells with ≥ 200 genes and genes found in > 3 cells. We were unable to retrieve information about empty droplet or doublet removal steps. However, we did not find any evidence of residual droplets and opted to not run EmptyDrops^125^ and proceeded with DoubletDetection and downstream analysis as indicated above.

Che et al^28^ (PBMC, TIL) provided preprocessed count data in .mtx as well as features and barcodes in .tsv format. After aligning the data to the GRCh38 reference genome using Cell Ranger v2.0, the authors had already filtered out cells with < 500 UMIs and removed cells which had a UMI count outside the range of *μ* + 3 *σ* to *μ* − 3 *σ* after log10-transformation per batch or “showed an unusually high or low number of genes”^28^, and removed cells with > 15 % mitochondrial genes. Hence, we did not perform empty droplet removal and we proceeded with DoubletDetection as outlined above.

Wang et al^29^ (PBMC) provided preprocessed gene expression counts, features, barcodes and metadata in .txt format, which we assembled into an AnnData object. The original study’s authors had aligned reads to the GRCh38 reference genome using CellRanger v3 and retained non-immune cells expressing between 200 and 6000 genes as well as ≤ 25 % mitochondrial counts and immune cells expressing between 200 and 4000 genes as well as ≤ 25 % mitochondrial genes. The authors had removed doublets using Scrublet^120^ and genes expressed in ≤ 50 cells.

Werba et al^56^ (TIL) provided filtered CellRanger outputs in .mtx format, as well as tabular barcodes and features. Sample identifiers representing each patient were provided alongside accession numbers on GEO. Sample metadata was available as supplementary files. Information from individual sample was concatenated to an .h5ad object. We observed a minimum UMI count per cell of 500, suggesting previous removal of empty droplets. We therefore did not rerun EmptyDrops and proceeded with DoubletDetection as outlined above.

### Extended PBMC resource

We obtained a total of 89 human PBMC datasets constituting all such datasets on the Chan Zuckerberg Initiative CellxGene data portal on September 21^st^ 2024 ([CZ CELLxGENE Discover - Cellular Visualization Tool](https://cellxgene.cziscience.com/)). We retained 17 datasets^126-142^ with a minimum of 25,000 cells and 5 patients per datasets (**Table S2**). We normalized cells to 10 000 counts per cell and log1p transformed the data (natural logarithm, pseudocount=1). We used our pretrained Compocyte model^143^ to infer PBMC cell types in each dataset and confirmed their lineage identity by inspecting coarse cell type markers from Compocyte (CD4 T, CD8 T, NK, cDC, p-DC, c-mono, nc-mono, B, plasma, erythrocyte, platelet).

### Human Tumor Atlas Network (HTAN) datasets

We obtained the scRNA-seq count matrices and corresponding metadata from the HTAN data portal (https://humantumoratlas.org/data-access, download date April 12^th^ 2026, data release 26.11.2025)^58,114,144-160^. We obtained filtered single-cell RNA-seq data which generally did not contain empty droplets as .mtx, .csv or .h5 files and converted them into AnnData objects. Gene identifiers were converted from Ensembl IDs to HGNC symbols. We then performed a simplified but robust preprocessing accounting for the large quantity and size of the datasets. To remove low quality cells and residual droplets, we removed cells with fewer than 200 detected genes, fewer than 500 total transcript counts, or more than 25% mitochondrial transcripts. The WUSTL dataset was provided without prior empty droplet removal. We therefore used the DropletUtils R package (‘emptydrops’) using a UMI count below 100 to define the reference droplet population and retained cells with a false-discovery rate below 0.05. Any sample with less than 500 retained cells was removed to avoid including samples with subpar preprocessing and skew cell type proportions by sampling effects.

To separate the leukocytes in each dataset, gene expression values were normalized to median library size using the scanpy.pp.normalize_total function, and log1p-transformed (natural logarithm, pseudocount=1) with scanpy.pp.log1p. The 10,000 most highly variable genes were selected using the seurat_v3 method for downstream principal component analysis. We selected the top 50 PCs consistently across datasets given that kneepoints estimation generally yielded lower PCs numbers not sufficient to identify granular cell populations. Importantly, the top 50 PCs explained a large amount of the total variance per dataset and were thus determined sufficient for downstream clustering and embedding. The cumulative variance explained by the top 50 PCs in each dataset was: 48.39% (HTAPP), 44.68% (MSK), 23.43% (Stanford), 47.28% (Duke), 36.42% (DFCI), 33.56% (CHOP), 45.20% (WUSTL), 34.93% (BU), and 33.6% (Vanderbilt). Next, we clustered the data with PhenoGraph using the k selection strategy indicated above (MSK: k=60, Duke: k=80, Stanford: k=60, Vanderbilt: k=90). The size of the WUSTL, CHOP, and HTAPP datasets led to prohibitive runtimes for PhenoGraph which is why we used Louvain clustering for these datasets instead. Similar to our k selection strategy, we selected the optimal Louvain resolution parameter (res) from a set of parameters (0.2, 0.4, 0.6, 0.8, 1.0, 1.2, 1.4, 1.6, 1.8, 2.0) by inspecting the pairwise rand indices (WUSTL: res=0.8, CHOP res=0.4, HTAPP: res=1.0) which allowed for stable separation of the leukocyte numbers indicated in **Table S2** as assessed by manual annotations using our Cytopus 2 markers. Subsequently, we predicted leukocyte subsets using our pretrained Compocyte classifier followed by manual validation of basic lineage markers in SEACell metacells (see: Data-driven uncertainty quantification). We then quantified the expression of Spectra factors defined on our CITE-seq dataset (see: Generating CITE-seq data for tumor infiltrating leukocytes) by taking the average expression of the top 10 marker genes for each Spectra factor in library-size normalized (10 000 counts per cell) and log1p-transformed (pseudocount=1, natural logarithm) scRNA-seq data. For each factor, at least 5 of the top 10 marker genes were retrieved ensuring that the resulting scores are representative of the original factors. We then calculated the average expression of each Spectra factor in each samples and cell type and explored their expression across metastatic and primary tumors.

### Entropy based cell type marker identification

Cell type markers are generally identified using differential expression tests such as Mann-Whitney U tests or classification approaches [logistic regression, area under the receiver operator curve (AUROC)]. While these methods are well suited for feature selection for cell type classification, they are not ideal for manual cell type annotation which mainly relies on qualitative expression patterns, given their lack in cell type exclusivity. For example, a gene broadly expressed in all cells with only slightly but consistently higher expression in the cell type of interest would get a low p value/high AUROC. Although filtering on fold changes reduces weak differential signals it does not distinguish broadly expressed genes from truly exclusive markers. To identify the most characteristic genes reflecting more exclusive biological features we therefore developed an entropy-based marker gene identification strategy.

To define robust cell type markers, for every independent dataset, we calculated differentially expressed genes using Benjamini-Hochberg corrected Mann-Whitney U tests. Next, we carried forward the overexpressed genes in the condition of interest (positive fold change) and calculated a normalized Shannon entropy per gene defined by:

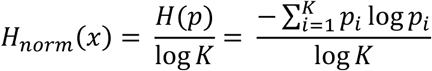

with i indicating one of K cell types and p being a probability vector of the gene expression across cell types with every p_i_ defined by:

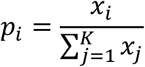

With x_i_ being the average transcript counts (UMIs) of gene x in cell type i.

The normalized Shannon entropy was then used to rank differentially-expressed genes (as determined by Mann-Whitney U tests) in every dataset and the genes with the lowest entropy were carried forward. We then counted the how often every gene appeared in the lowest entropy markers across all the datasets and considered the ones found in at least half of the datasets characteristic. To identify TARM markers, the number of top genes to focus on was empirically defined as the lowest number of genes where this analysis strategy yielded genes detected in at least 5 of 7 datasets (top 100 entropy markers, fdr=0.05, minimum log_2_ fold change=0.5). To identify markers for CITE-seq-defined memory T cell populations we relaxed these criteria given the lower cell numbers (top 100 entropy markers, fdr=0.3, minimum log_2_ fold change=0.5). For surface marker identification from CITE-seq data, we modified this strategy given the strong inter-sample variation in surface protein staining: We calculated the differentially-expressed DSB-normalized CITE-seq antibody derived tag (ADT) counts within each sample and performed and the entropy ranking described above which was performed per sample. To deal with spurious background expression not accounted for by DSB we focused on highly overexpressed genes (log_2_-fold change>0.5, fdr=0.05, top 100 entropy genes) which identified CD64 and CD151 in the top 25 genes found in 4 representative NSCLC CITE-seq samples with sufficient cell numbers (>3500 cells).

### Tumor immune microenvironment identification

To evaluate coordinated cellular dynamics across datasets, we first calculated the pairwise correlation of all cell type pairs using their frequencies from our TIL Suco atlas. To prevent artifacts arising from the hierarchical nature of the immunophenotyping panel, correlations between directly connected parent-child cellular lineages were explicitly excluded from the analysis. For all remaining valid cell type pairs, the Spearman rank correlation coefficients and corresponding asymptotic p values were calculated. To eliminate spurious correlations, raw p values were globally adjusted using the Benjamini-Hochberg false discovery rate procedure. Gene-gene pairs failing to meet a strict significance threshold (FDR<0.001) were masked.

To identify conserved cell type modules we performed agglomerative hierarchical clustering using the average linkage method (UPGMA). Discrete cell type modules were generated via flat clustering by pruning the dendrogram at a fixed cophenetic distance (0.75). To isolate robustly correlated cell types, we performed a rigorous two-step filtration of the resulting clusters. First, a cluster was immediately discarded if it contained fewer than four cell type members. Second, the remaining clusters were assessed for their average absolute pairwise correlation coefficients (only using off-diagonal correlations). Clusters with lower than 0.4 average absolute correlation were rejected resulting in our final list of tumor immune microenvironment (TIME) programs. Each cell type belonging to any rejected cluster was assigned a unique cluster identifier to not bias the adjusted rand index calculation below We then evaluated the stability and reliance on individual datasets of the filtered cell type TIME programs using a leave-one-dataset-out cross-validation method. For each validation fold, one unique dataset was entirely withheld, and the entire analytical pipeline was re-executed from raw frequencies using only the remaining training cohorts. This re-execution included pairwise Spearman correlation, fold-specific false discovery rate correction, average linkage tree construction, flat clustering tree cuts, and post-clustering module purification using the minimum average correlation approach described above. To quantify clustering stability, we then calculated the adjusted rand index between cross-validation folds and the baseline which was consistently >0.6 for all iterations. Additionally, we confirmed the presence of the major cell type patterns representing the TIME programs in each validation fold.

### Compocyte

We built our cell type annotation package Compocyte in Python, as a wrapper around three existing machine learning packages: CatBoost^161^, PyTorch^162^, and Scikit-learn^163^ to adopt them for modular, hierarchical classification of single cell data. Compocyte hierarchically assembles classifiers along a user-defined ontology. This modularity allows classifiers to be flexibly exchanged based on a user-defined tree-like cell type hierarchy in nested dictionary format, it constructs a directed NetworkX^164^ graph to store cell type relationships and information on each cell type label. Cell type ontologies can be directly imported from our newly developed single cell knowledge base Cytopus 2 to facilitate using, storing and sharing single cell ontologies. The single cell data is provided in AnnData^23^ format as input for training and inference. Compocyte automatically infers one label column for each level of the hierarchy using the provided hierarchy graph. Although this manuscript explores applications of Compocyte on single cell RNA sequencing (scRNA-seq) data, it can theoretically be adopted with other single cell measurements in tabular format (e.g. CITE-Seq, mass cytometry) although different data normalization and model parameterization may be required to achieve optimal performance.

#### Hierarchical classifier

Standard single-cell classifiers typically employ flat architecture, assigning a single terminal label to each input feature vector. In contrast, Compocyte utilizes a top-down hierarchical architecture, implementing a local-classifier-per-parent-node strategy^165^. In this framework, the classification ontology is structured as a directed graph. At each parent node, an independent local classifier evaluates incoming cells and routes them into progressively higher-resolution categories. This sequential partitioning continues down specific hierarchical branches until the cell reaches a terminal leaf node lacking further subdivisions.

The ontology defined within Compocyte enforces a strict phenotypic subsetting constraint that is every child node must represent a more granular transcriptomic sub-population of its immediate parent. For example, a broad ‘T cell’ parent node validly branches into ‘αβ T cell’ and ‘γδ T cell’ child subsets. By strictly requiring that child labels remain phenotypically nested within the boundaries of their parent nodes, every cell passed through the classifier follows a biologically logical annotation path, categorized strictly by its observed transcriptomic state.

#### Data preprocessing

For node-specific predictions, Compocyte dynamically instantiates local classifiers as either PyTorch-based neural networks, PyTorch- or scikit-learn-based logistic regression models, or CatBoost-based gradient-boosted trees. As standard input, the framework requires transcriptomic profiles to be normalized to 10,000 counts per cell and log1p transformed (natural logarithm, pseudocount=1). If unnormalized data is encountered, Compocyte automatically extracts raw counts (via anndata.raw or anndata.layers) to apply this transformation natively. Furthermore, across all benchmarking experiments, input dimensions were uniformly restricted to 5,000 highly variable genes prior to local-classifier-specific feature selection to maximize the signal-to-noise ratio.

#### Balancing ontological complexity and error propagation

The structural depth of the user-defined ontology directly determines Compocyte’s adaptability, as highly complex taxonomies allow for the highly selective isolation and retraining of specific subtrees. However, deeper hierarchical topologies inherently carry a theoretical risk of multiplicative error propagation. An initial misclassification at an upstream parent node irreversibly routes the cell down an incorrect lineage. Consequently, defining the optimal ontology represents a context-dependent trade-off between annotation resolution and algorithmic robustness. Notably, despite utilizing a highly complex, multi-level hierarchy that maximizes the potential for such error propagation, Compocyte’s architecture systematically mitigated this risk, successfully outperforming standard flat classification models in our benchmarks (**Fig. 4b**).

To prevent forced misclassification, Compocyte implements partial label rejection via non-mandatory leaf node prediction (NMLNP)^165^. During inference, if the softmax probability at a given branching point fails to exceed a predefined confidence threshold, sequential routing halts. Instead of forcing assignment to a terminal leaf node, the cell inherits the label of its most granular, confidently predicted parent node. Unlike flat classifiers, which often yield biologically erroneous annotations when prediction confidence is low, this hierarchical mechanism ensures that cells with ambiguous transcriptomic profiles receive a broader, but strictly accurate, annotation.

#### Feature selection and training

To train individual local classifiers, input cells are partitioned strictly according to the defined ontology. A given parent node is trained using all cells annotated with its immediate child labels, as well as any cells possessing downstream terminal descendants of those children (e.g., a node separating αβ T cells into CD4 and CD8 subsets is trained using all cells with any of the downstream labels). Prior to both training and inference, local transcriptomic features are standardized to a unit variance (standard deviation of 1). Zero-mean centring is explicitly omitted to retain the inherent sparsity of the single-cell expression matrix, thereby preventing prohibitive memory scaling during computation. Following variance standardization, the most discriminative genes for each local classifier are selected based on their ANOVA F-value. The absolute number of selected features is dynamically scaled according to the available sample size, utilizing a heuristic adapted from Google’s Rules of Machine Learning to determine the number of features (*Martin Zinkevich*, accessed May 17^th^ 2026, https://developers.google.com/machine-learning/guides/rules-of-ml):

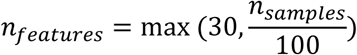

#### Default neural network architecture and training regime

Compocyte’s default local classifier is a shallow, highly efficient neural network, which empirically achieved the highest F1 scores across our benchmarks (**Fig. 4b**). The architecture consists of two 64-neuron hidden layers, each followed by batch normalization and a 40% dropout layer to heavily penalize overfitting, terminating in a softmax output layer. Network weights are assigned via Glorot initialization^166^, with biases initialized to zero. Models are trained for a fixed duration of 40 epochs using stochastic gradient descent (batch size = 64, momentum = 0.5). To ensure rapid and stable convergence within this fixed training window, learning rates are dynamically modulated using the OneCycle scheduler^167^ (initial learning rate = 0.01, maximum = 0.1). Because hierarchical single-cell datasets exhibit profound taxonomic imbalances, we mitigate majority-class dominance by applying a modified class-balanced focal loss function^168^ (β=0.8, γ=2.0). Although the training loop does not utilize early stopping, optimal convergence is ensured by systematically extracting the final model parameters from the epoch that achieved the lowest validation loss. Because this network is intentionally constrained in width and depth the architecture achieves competitive training and inference runtimes natively on standard CPUs without requiring GPU acceleration (Fig. 4e). Moreover, increasing parameter numbers yielded no corresponding gains in classification performance.

#### Hierarchical inference

During inference, query datasets are automatically standardized to match the training data format (normalized to 10,000 counts and log1p-transformed). To perfectly align feature spaces, any classifier-specific training genes missing from the query matrix are appended and zero-padded. Unlike the independent node sampling utilized during training, inference executes a strict sequential traversal of the classification hierarchy. Cells are evaluated at the root node and iteratively routed to subsequent downstream classifiers based on their predicted subset. To prevent forced misclassification, Compocyte dynamically applies partial label rejection at each branching point. A cell advances to a child node only if the maximal softmax activation of the output layer exceeds a predefined confidence threshold. If this threshold is not met, hierarchical routing halts, and the cell retains the most granular, confidently assigned parent label. As is standard for neural network inference, dropout and batch normalization layers are strictly disabled.

#### Decentralized, node-specific hyperparameter optimization

Because Compocyte relies on an architecture of independent local classifiers, each with distinct sample sizes, feature subsets, and class imbalances, global optimization frameworks (e.g., Ray Tune) and manual feature engineering are incompatible. To maximize out-of-dataset generalizability, we developed a custom, decentralized grid-search framework (the Tuner class) designed to optimize hyperparameters independently for every node in the taxonomy. For each trial run, the Tuner evaluates a comprehensive hyperparameter grid encompassing network architecture (hidden layers, neurons), regularization (dropout), training dynamics (epochs, batch size, momentum, OneCycle learning rate boundaries), and class-balanced loss parameters (β, γ). To maintain computational efficiency during optimization, trials can be executed on a downsampled data fraction (dictated by a user-specified test factor). Model generalizability is rigorously estimated via k-fold cross-validation, guided by a predefined metadata key containing the cross-validation folds (adata.obs). Performance metrics from all cross-validation runs including correctness at varying softmax activation thresholds are systematically logged into an SQLite database. During final model compilation (train_from_tuner), Compocyte automatically queries this database to assign the empirically optimal hyperparameter combination for each local classifier. Selection is strictly prioritized by local accuracy, followed by the derivation of the optimal softmax threshold to maximize overall accuracy.

#### Parallelized training

Because the classification ontology operates as a directed graph of independent local classifiers, each node can be trained entirely asynchronously. This structural autonomy is possible because each classifier executes node-specific feature selection and requires only the subset of training data corresponding to its downstream descendants, remaining strictly decoupled from upstream routing dependencies. Consequently, Compocyte natively supports highly parallelized computation, which is implemented via the Python *multiprocessing.Pool* module to concurrently orchestrate independent training processes across available CPU cores. To eliminate the severe synchronization bottlenecks inherent to shared-memory architectures, these spawned processes execute in strict memory isolation. While this architecture inherently increases peak RAM utilization, as each parallel worker must load an independent data copy, it strategically bypasses the need for a centralized, inter-process manager. Eliminating this manager removes the massive computational and memory overhead associated with continuous cross-process locking and data synchronization, ultimately maximizing training throughput and efficiency in multi-core environments.

#### Out-of-core computation and resource management

Current automated classification frameworks are frequently limited by prohibitive memory requirements, as they often require computation on the entire expression matrix. To overcome this limitation and enable population-scale training, Compocyte implements a highly optimized, out-of-core data management architecture. By default, the framework preserves expression matrix sparsity utilizing the compressed sparse row (CSR) format until algorithmic steps explicitly require densification. To further conserve memory, global feature selection is algorithmically capped at a representative subset of 100,000 cells. When training a neural network-based local classifier on large datasets (default threshold: >1,000,000 cells), the sparse expression matrix of the training subset is converted into a Dask array. Dask enables out-of-core processing by partitioning the global matrix into manageable chunks. During training, individual chunks are sequentially mapped to dense NumPy ndarrays strictly at runtime for each mini-batch. Because only the current mini-batch is densified and transferred to the hardware device, this architecture effectively circumvents both system RAM and GPU VRAM saturation, enabling hardware-accelerated training on datasets of virtually unlimited size, though of course at higher input/output cost. To offset the high CPU overhead inherently associated with runtime densification, Compocyte dynamically assigns parallel data-loading workers to prevent GPU/CPU starvation.

#### Mitigating class imbalances in cellular ontologies

Biological ontologies inherently exhibit profound class imbalances, primarily driven by the low frequency of rare cell states. Within the Suco resource, this translates into highly skewed label distributions at nearly every branching point of the ontology. Without algorithmic intervention, standard classifiers typically minimize validation loss trivially by defaulting to majority-class predictions, entirely failing to learn the discriminatory transcriptomic features of rare subsets. Because conventional re-sampling techniques (e.g., synthetic data augmentation or aggressive downsampling) introduce prohibitive runtime overhead and risk severe information loss in single-cell data, we implemented a class-sensitive learning strategy. Specifically, local classifiers in Compocyte are optimized utilizing a class-balanced formulation of focal loss ^168^, integrated via the balanced-loss Python package. This dual-penalty mathematical framework acts on two fronts: focal loss dynamically scales the gradient based on prediction confidence (assigning exponentially higher penalty weights to hard-to-predict samples), while the class-balancing term simultaneously applies a static corrective weight inversely proportional to the true label frequency. Together, this approach forces the local classifiers to penalize misclassifications of rare cell types severely, ensuring robust feature extraction across the entire biological hierarchy.

### Metrics

Because traditional flat classification metrics fail to capture the topology of ontological trees, evaluating hierarchical classifiers requires specialized mathematical approaches. Specifically, an optimal hierarchical evaluation framework must satisfy the three core criteria established by Kiritchenko et al.^169^. First, the metric must account for partial classification correctness by proportionately reducing penalties for errors that remain within the true ontological subgraph. Second, penalties must scale strictly with topological distance: traversing deeper into the hierarchy must be mathematically favored if it approaches the true category, but disproportionately penalized if it diverges further away compared to halting at the previous parent node. Finally, the framework must impose the strictest penalties for errors occurring at higher, fundamental levels of the hierarchy (e.g., major lineage misclassifications) compared to minor errors at terminal leaf nodes. To fulfill these requirements, we use modified precision, recall and F-Scores which use a set-based approach^165,169^. To define the sets:

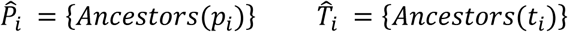

with *p*_*i*_ and *t*_*i*_ being the predicted and true leaf-label of instance (i.e. cell) *i*, respectively. In other words, start at the predicted/true label and move toward the root of tree, collecting each label one comes across. The modified precision, recall and F-score are defined as follows with *β* determining the weight of hR and hP. We used *β*=1 for our benchmarking.

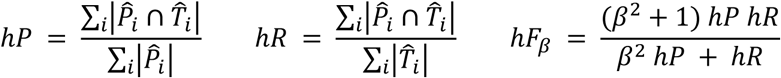

In cases of over-specialization, a potentially desirable trait^165^, we further modified hP to avoid penalizing over-specialization. Over-specialization occurs when |*P̂*_*i*_ ∩ *T̂*_*i*_| = |*T̂*_*i*_| *and* |*P̂*_*i*_| > |*T̂*_*i*_|. In other words, over-specialization occurs when prediction provides labels that can be neither confirmed nor falsified due to lack of a ground truth label at the corresponding hierarchy level beyond the last matching label. In these cases, there is total overlap at the hierarchy levels that have a ground truth label provided. To account for these cases, we adjust hP in the following way in cases where |*P̂*_*i*_ ∩ *T̂*_*i*_| = |*T̂*_*i*_|:

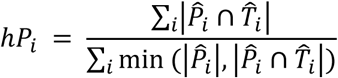

For every sample *i* with |*P̂*_*i*_| > |*P̂*_*i*_ ∩ *T̂*_*i*_|, this sets single-prediction hP to 1 and drives the aggregate hP toward 1 as well.

### Data-driven uncertainty quantification

To streamline the manual verification of automated annotations, we developed a data-driven uncertainty metric designed to actively direct expert raters toward predictions that are most challenging for automated classification. To validate this approach, we systematically evaluated candidate uncertainty metrics by benchmarking them against two ground-truth uncertainties: the human raters’ subjective uncertainty (quantified via a five-point Likert scale) and the topological tree-distance between the model’s predicted labels and the final expert annotations.

Conventional single-cell uncertainty metrics typically rely on isolated, cell-level evaluations or raw output layer activations (e.g., softmax probabilities). Because these raw activations often fail to represent the true error probability of deep neural networks, we quantified uncertainty based on label consistency across transcriptomically homogeneous groups of cells. We derived these granular phenotypic cell groups using the SEACells metacell algorithm^75^. To balance computational granularity with the structural constraints of our ontology, we initialized the algorithm to compute 90 metacells per dataset, directly mirroring the approximate expected number of distinct cell states in our classification hierarchy. The SEACells model was monitored for convergence by tracking the squared error across iterations, followed by the hard assignment of individual cells to metacells based on maximum membership values. By evaluating predictions within these defined SEACell boundaries, we established a localized label consistency metric (see below). This metacell-derived consistency score demonstrated a robust correlation with both the raters’ perceived uncertainty and the average tree-distance of the raters’ annotations within each metacell, establishing it as a highly reliable heuristic for targeted human review.

#### Average topological tree-distance

We calculated the pairwise hierarchical tree-distance between all single-cell predictions within a given SEACell. Both the average and standard deviation of these pairwise distances were evaluated as candidate uncertainty metrics. Average tree-distance exhibited a superior correlation with both ground-truth label accuracy and perceived human uncertainty compared to standard Shannon entropy (calculated via scipy v1.15.2).

#### Epistemic uncertainty estimation via Monte Carlo dropout

While dropout is routinely employed during training to penalize feature co-adaptation and mitigate overfitting^85^, it is conventionally disabled during inference. However, by retaining stochastic dropout during forward passes at inference, a model’s epistemic uncertainty can be robustly approximated^170^, as highly confident predictions should remain stable despite the random ablation of input data. To ensure methodological compatibility across our diverse local architectures including neural networks, boosted trees, and logistic regression classifiers we engineered Monte Carlo (MC) dropout specifically as feature dropout applied directly to the input layer. For a user-specified number of inference iterations (n), the classifier evaluates the input matrix with 50% of the transcriptomic features randomly masked during each pass. The resulting softmax output activations are then aggregated across iterations, allowing both the mean and standard deviation of the outputs to serve as estimators of prediction stability.

#### Metric evaluation and composite uncertainty formulation

To empirically validate these candidate metrics, we applied the respective classifiers to the PBMC cross-validation cohort utilizing 10 MC dropout iterations, subsequently deriving SEACell metacells as described above. To ensure dimensional comparability and facilitate the additive combination of independent metrics without undue weighting, all per-SEACell uncertainty values were strictly min-max scaled. We systematically calculated the Spearman correlation coefficients between all individual metrics, as well as their correlations with the true topological tree-distance between ground-truth and predicted labels. We observed that the MC-derived metrics did not correlate with the label distance-based metrics, indicating that they capture distinct, non-redundant dimensions of model uncertainty. Through systematic combinatorial testing, we determined that the most robust composite metric which yielded the highest Spearman correlation with ground-truth accuracy, was the unweighted sum of the normalized average pairwise tree-distance and the normalized MC dropout standard deviation of output activations, averaged per SEACell.

### Benchmarks

#### Classification performance benchmarks

To evaluate out of sample prediction accuracy across the twelve independent PBMC datasets in the Suco resource, we implemented a twelve-fold leave one dataset out cross validation strategy. Under this paradigm, each distinct dataset served sequentially as an entirely independent test set, while the remaining datasets were utilized to form the training and validation cohorts. Because several other classification methods reached prohibitive memory limitations when processing the unmanipulated Suco resource, we utilized a partially stratified downsampling approach. We constructed a standardized global benchmarking pool capped at 500,000 cells which enabled us to compare Compocyte’s performance to methods otherwise limited by prohibitive compute resource usage.

For our benchmark we retained 41,667 cells per dataset, which represents the 500,000 total cells divided by the 12 datasets. For the two datasets whose total cell counts fell below this threshold, all available cells were retained. To reach the target of 500,000 cells, we randomly sampled from the remaining ten datasets without further stratification. This workflow effectively minimized dataset specific composition biases in the training matrix while preserving the strict statistical independence required for the leave one dataset out evaluation.

##### Benchmarked model selection

To ensure an unbiased evaluation across disparate algorithmic architectures where exhaustive hyperparameter optimization for every tool is computationally prohibitive, all benchmarked methods, including Compocyte, were evaluated using their default parameter configurations. We also used central processing units (CPU) only to benchmark the models because not all of them (particularly TransferData and CHETAH) supported graphics processing unit (GPU) computation. Preprocessing pipelines were executed strictly according to the documentation and tutorials provided by the respective authors. We selected six of the most widely adopted single cell classifiers across the Python and R ecosystems: CHETAH^79^, TransferData^171^, scANVI^12^, scPoli^76^, treeArches^77^, and CellTypist^78^. Because these baseline methods require standard, one-dimensional label inputs and do not natively accommodate multi-level ontological hierarchies, they were trained using the most granular available annotation layer per cell, in line with standard benchmarking conventions.

Several candidate methodologies were excluded from the final benchmarking based on strict computational or structural criteria. The methods SingleR^172^, SingleCellNet^173^, and scID^174^ were excluded due to training times exceeding one week on our benchmarking dataset. The scPred^175^ algorithm could not be evaluated due to a critical exception within its trainModel function that remains unaddressed in the official repository (Issue 30, as of April 30^th^ 2026, https://github.com/powellgenomicslab/scPred/issues/30). Finally, Garnett^176^, scAnnotatR^177^, and HierarchicalFit^178^ were omitted because they depend fundamentally on user curated cell type marker lists rather than operating as purely data driven, reference based classifiers.

##### Information leakage and random holdout control simulation

To quantify the impact of dataset specific technical batch effects on generalizability, and to demonstrate how conventional validation strategies can artificially inflate performance metrics, we compared our leave-one-dataset-out paradigm against a randomized holdout baseline. To maintain identical cell numbers, class proportions, and training dimensions across both evaluation strategies, we permuted the dataset batch identifiers across the global pool using a NumPy random number generator.

By shuffling these dataset indices prior to splitting the cross-validation folds, we intentionally broke the isolation between technical batches. This control experiment effectively simulated a standard intra dataset random split, wherein cells from the same technical batch are uniformly distributed across both training and testing sets, allowing us to measure the degree to which information leakage masks overfitted model architectures.

###### scANVI

We used scANVI with 5,000 highly variable genes obtained using the seurat_v3 algorithm in scanpy. Layer normalization was enabled for encoder and decoder. Batch normalization was disabled. Dropout rate was set to 0.2, and covariate encoding was enabled. The number of hidden layers for encoder and decoder was set to 2. The resulting autoencoder was trained for the default number of maximum epochs given by 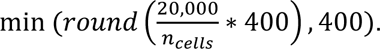 The scANVI model was then constructed from this pretrained autoencoder, trained for a maximum of 20 epochs with default ZINB loss and n_samples_per_label set to None (no subsampling). Following the loading of query data, we train unsupervised for another 100 epochs with weight_decay set to 0 before predicting cell type labels.

###### scPoli

We subset data to 3,000 highly variable genes, again using seurat_v3. We provided the dataset as condition keys. We set scPoli up to use three embedding dimensions and then trained for 50 epochs and 40 pretraining epochs, with eta=5, validation prototype loss as early stopping metric, early stopping set to min mode, with threshold 0 and patience 20. We had also enabled learning rate reduction with patience set to 13 and the reduction factor set to 0.1. After loading the query data, the model is trained unsupervised for another 50 epochs, 40 pretraining epochs with eta set to 10 before predicting cell types

###### treeArches

Training data was subset to 2000 highly variable genes using seurat_v3. We supplied the dataset key as covariate key, ran SCVI (with 2 layers in encoder and decoder, covariate encoding set to True, covariate deep injection set to True, layer normalization for encoder and decoder and without batch normalization) for a maximum of 80 epochs. We manually created a classifier tree following the hierarchical structure we used for labelling and training Compocyte, trained using a KNN classifier, on the embedding data generated from scVI and enabled cell rejection based on reconstruction loss with an allowed false negative rate of 0.5 for determination of reconstruction error threshold. We used 50 nearest neighbors but set dynamic_neighbors to *True* allowing for flexibility with small cell populations. For inference, we once again loaded the query data and trained the scVI model unsupervised for 80 epochs before retrieving the embedding and predicting cell types using the treeArches model.

###### CellTypist

Training data genes were subset to 5,000 highly variable genes using seurat_v3. Training data and test data were separately total-count normalized to 10,000 counts per cell then log1p-transformed. The CellTypist model was trained utilizing the two-step feature selection but without standardizing to mean 0, because setting CellTypist’s with_mean argument to *True* lead to excessive RAM usage, that would have precluded the experiment with the significant resources available (up to 1 TB of RAM). For inference, we used CellTypist’s majority voting feature.

###### CHETAH

Count data, observation and variable metadata were exported from scanpy into .csv format. Preprocessing for this method was performed in R. Briefly, we generated SingleCellExperiments for our training and test data from.csv files. We normalized and log-transformed our training count data using logNormCounts and subset to 5,000 highly variable genes using getTopHVGs. We then applied logNormCounts on our test data, generated 30 principal components using prcomp and from this calculated a TSNE embedding with reducedDim. A SingleCellExperiment class containing count and TSNE data for our test set and a SingleCellExperiment containing count data and cell type metadata for our training set were provided to CHETAH. CHETAH then conducted hierarchical reference mapping to provide cell type predictions for our test data.

###### TransferData

Preprocessing for this method was also performed in R. We created a Seurat object and normalized and log-transformed the count data using NormalizeData’s default “LogNormalize” method separately for every dataset in the training set. We then reduced to 2000 highly variable genes using FindVariableFeatures with the vst method. We built integration anchors using FindIntegrationAnchors and 30 dimensions of the default canonical correlation analysis. We integrated our training using these anchors, scaled using ScaleData and ran principal component analysis with 30 dimensions. After this, we normalized and feature-reduced our test data as above with our training data. Finally, we applied FindTransferAnchors using the reference data’s PCA reduction and transferred cell type labels according to these anchors.

###### Resource benchmarks

To compare resource requirements between methods, we simulated training and testing on datasets of 10,000, 25,000, and 50,000 cells. These datasets were sampled from our PBMC training data using a numpy random number generator, saved and reused across methods to ensure comparability. We conducted these tests on our compute cluster running IBM Spectrum LSF for resource allocation. We requested 8 cores of compute on the same cluster node for every benchmarking job across all methods. The memory_profiler and Rprof provided unrealistically high maximum memory estimates, in many cases exceeding the maximum memory requested from and allocated to the job by LSF. For this reason, we used the maximum memory provided per job by the LSF summary output. This included the entire job including data loading, training/preprocessing and inference. The start time for the training part of the process was measured after data loading was complete. The start time for the prediction part and end time for the training part of the process was measured as soon as the prediction process on test data was able to start. We repeated measurements 10 times for every method and cell count. Parameters for training and inference were the same as described for the performance benchmarking above.

For Compocyte, we used the more complex model hyperparameters that we also used for the training in our pretrained PBMC model to provide a higher boundary for resource usage.

### Pretrained models

To maximize out of sample prediction performance for the pretrained PBMC model, we conducted hyperparameter and architecture optimization at the local classifier level using the custom Tuner class described above. We systematically evaluated 982 distinct hyperparameter combinations for PyTorch based neural networks, varying feature counts, hidden layer dimensions, training epochs, batch sizes, starting learning rates, and momentum. Concurrently, we evaluated boosted trees classifiers across seven distinct feature number thresholds. All three evaluated architectural paradigms, including logistic regression, gradient-boosted trees, and neural networks, generate output layer activations or analogous measures of prediction certainty. We leveraged these probabilistic outputs directly within the optimization loop to derive node specific prediction thresholds that optimally segregated true positive from false positive assignments, thereby maximizing local classification accuracy. This optimization protocol was executed on a stratified one fourth fraction of the training data subset, comprising 100,000 cells in total, with individual trial runtimes strictly capped at ten hours. Furthermore, we implemented the previously described cross validation framework, iteratively utilizing discrete datasets from the training corpus as independent holdout folds.

#### Population scale model compilation and external benchmarking

Following this optimization phase, the final pretrained PBMC model was compiled and trained using the empirically derived hyperparameter configurations across the entire corpus of 5.4 million cells in Suco PBMC version 1.3 (10.5281/zenodo.20113753), limited however to a maximum of 1 million samples per local classifier after stratified downsampling by dataset at the local classifier level. This ensured avoiding sample loss at smaller nodes while limiting resource use. We subsequently generated an unbiased external test set to verify that the superior performance of Compocyte was not merely an artifact of our highly curated internal annotations. Specifically, we leveraged an extended PBMC resource containing heterogeneous, author-provided labels and standardized this nomenclature by mapping all external annotations to our defined unified ontology (**Table S5**). We then benchmarked the fully pretrained Compocyte model against the most granular immune cell classifier available in CellTypist (“Immune_All_High” v2^78^, which was trained on approximately 330,000 cells across 19 studies) as well as Azimuth^171^ (which utilizes a PBMC reference of 162,000 cells derived from a single study) across eleven completely independent datasets. To ensure a strictly equitable comparison, all output labels generated by CellTypist and Azimuth were formally mapped back to our standardized cell type ontology. Ultimately, despite the inherently lower cell type resolution native to the pretrained CellTypist and Azimuth architectures, Compocyte robustly achieved vastly superior global classification performance.

### Synthesizing myeloid and T cell classifiers

The modular architecture of Compocyte facilitates the integration of overlapping subtree classifiers trained on entirely independent datasets. For example, classifiers trained independently on distinct cell lineages can be computationally synthesized into a unified model by introducing an upstream parent classifier that segregates these broad lineages. We refer to this process as ‘classifier synthesis’. To empirically demonstrate the non-inferiority of this modular synthesis approach compared to standard combined data training regimes (‘data synthesis’), we independently trained and subsequently merged distinct myeloid and T cell classifiers.

To construct these lineage specific models, we obtained a pan cancer T cell atlas^42^ and a separate pan cancer myeloid cell atlas^61^. For the T cell atlas, the original study’s authors aligned the reads and filtered out cells containing fewer than 200 detected genes or greater than ten percent mitochondrial reads, alongside filtering genes detected in three or fewer cells. The authors additionally utilized Scrublet^120^ to computationally remove doublets, though upstream preprocessing for the previously published datasets in this atlas was not explicitly defined. For the myeloid cell atlas, the original study’s authors filtered the data to retain cells exhibiting between 2,000 and 40,000 unique molecular identifiers, between 500 and 5,000 detected genes, and fewer than ten percent mitochondrial reads. The authors also explicitly removed a contaminating cluster exhibiting both T cell and myeloid lineage markers which Scrublet^120^ identified as having a high doublet content. For the previously published data within the myeloid atlas, the authors reported applying identical preprocessing steps, except for lowering the unique molecular identifier threshold to 300 and raising the mitochondrial read cutoff to twenty percent for the contained inDrop datasets. Similarly, for the contained MARS Seq datasets the authors utilized the 300 unique molecular identifier cutoff alongside an additional twenty percent cutoff for ERCC spike in control genes.

To homogenize the disparate author provided labels across all datasets utilized in this specific experiment, we mapped them to a simplified tumor infiltrating leukocyte ontology referenced in **Table S7**. All raw count data was total count normalized to 10,000 counts per cell and log transformed utilizing the natural logarithm with a pseudocount of one. Following this standardization, we trained two independent Compocyte lineage classifiers comprising one dedicated myeloid model and one dedicated T cell model. To simulate a common real-world scenario where researchers possess abundant coarsely labeled data but lack access to granular lineage specific datasets, we trained a third upstream parent classifier using our independent Suco tumor infiltrating leukocyte atlas to broadly differentiate between myeloid and T cells within the tumor microenvironment. These three distinct classification nodes were then programmatically synthesized into a single cohesive hierarchy utilizing the native export and import features of Compocyte. To establish an independent evaluation cohort, we completely withheld the breast cancer dataset generated by Bassez et al^60^ to serve as our test data.

### Human-in-the-loop annotation

To test our expert-guided annotation strategy we chose the Salcher et al.^179^ lung cancer atlas which included 865,503 immune cells from 324 lung cancer or lung tissue samples from 187 patients in 13 independent datasets from 11 studies^38,62,179-187^. We identified immune cells using the authors’ cell type labels. Data were generated using the 10x Genomics^62,180-187^, InDrop^38^, and BD Rhapsody platforms ^179^ and aligned using the CellRanger v5.0.0, nf-core^188^ and Seven Bridges Genomics cloud pipelines. The authors performed coarse labeling and label transfer with scANVI, followed by doublet removal as well as removing low-quality cells with high mitochondrial genes, low count number as well as the removal of sparsely expressed and mitochondrial genes.

We partitioned this global atlas into two distinct training and validation cohorts alongside one completely independent test dataset. The first training cohort comprised datasets from five studies, specifically Adams and Kaminski 2020^180^, Chen and Zhang 2020^181^, Habermann and Kropski 2020^182^, He and Fan 2021^183^, and Kim and Lee 2020^184^. The second training cohort incorporated data from Lambrechts and Thienpont 2018^185^, Laughney and Massague 2020^186^, alongside three distinct sample partitions from Leader and Merad 2021^62^. The independent test dataset consisted of samples from Madissoon and Meyer 2020^189^, the UKIM cohort^179^, and Zilionis and Klein 2019^38^.

We subsequently derived phenotypic metacells across these cohorts utilizing the SEACells algorithm^75^ according to the methodology detailed above. We then deployed a baseline Compocyte classifier onto the first training dataset. This baseline model was originally trained on the Suco tumor infiltrating leukocyte atlas (v1.1, 10.5281/zenodo.15350418)^20^ and our HTAN CITE seq dataset. It utilized the standard local classifier neural network architecture described in the Compocyte methodology section, with additional memory cell annotations derived from our HTAN CITE seq data appended to the taxonomy.

Following inference on the first training cohort, each SEACell was assigned its most frequently predicted annotation. These metacell level predictions were then subjected to manual expert review and correction. To maximize efficiency, human review was prioritized toward the most uncertain annotations as defined by our composite uncertainty metric. We then assessed the expression of cell type identity markers from Cytopus on SEACells selected for review. Next, the expert validated annotations were integrated back into the training corpus to compile an iteratively improved Compocyte classifier. We subsequently performed inference on the second training dataset using this updated model and repeated the identical manual review process. This iterative workflow yielded a final expert review guided Compocyte model. We ultimately benchmarked this refined model against both the first iteration model and the original baseline Compocyte model using the completely withheld independent test data to rigorously quantify the performance gains achieved through human-in-th-loop curation.

### Integration

To estimate the loss of biological information by common integration approaches, we tested integration with scVI on our Suco resource. We first subset our TIL data to 2,000 highly variable genes using the scanpy implementation of the Seurat v3 algorithm. We then generated 10-, 30-and 100-dimensional scVI embeddings by training an scVI model with 2 layers for encoder and decoder with 128 hidden neurons each and negative binomial gene likelihood with 90/10%, for a number of max epochs set by scVI’s internal get_max_epochs_heuristic without early stopping. After detecting no qualitative differences in the outcome intra-cluster cell type mixing between different numbers of hidden dimensions we decided upon 30 hidden dimensions. We repeated this first part of the experiment with 30 hidden dimensions for a subset of our PBMC Suco resource including four dataset (Oelen_E1, Oelen_E2, Hao_E1, Hao_E2). Based on the extracted embeddings, we computed Phenograph clusterings using k=10 to k=100 in steps of 10. We chose the k parameter based on clustering stability as measured by pairwise rand index and outlined above.

To quantify cell type mixing per cluster we calculated the effective number of lineages per cluster, inspired by species diversity measures from ecology^190^. This measure assigns meaning to both the distribution of labels in different clusters, as well as to their (dis)similarity. To quantify dis(similarity), we first generated a distance matrix by calculating tree-distances for all possible label combinations in the respective TIL/PBMC hierarchy graph. We then min-max scaled these distances to the range 0 to 1, shrank the distance matrix to all combinations between available labels in the cluster (*D*) and calculated the proportion of labels per cluster (p) to address label distribution. Based on this, the effective number of lineages was calculated as follows:

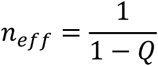

With Rao’s Q being defined as:

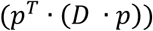

## Data availability statement

The single cell universal classification omnibus resources for PBMC (10.5281/zenodo.13709549) and TIL (10.5281/zenodo.15350417) have been deposited on Zenodo. CITE-seq data from metastatic human tumors will be available on the HTAN data portal shortly and before publication (https://humantumoratlas.org/data-access). PBMC data from the ANTICIPATE trial will be made available upon publication on Zenodo and EGA.

## Code availability statement

Computer code for Cytopus 2 (https://github.com/wallet-maker/cytopus) and Compocyte (https://github.com/WALL-E-Lab/Compocyte) is available on GitHub. Compocyte open-weight models for PBMC (10.5281/zenodo.19708294) and TIL (10.5281/zenodo.19707909) are available on Zenodo. Computer code required for reproducing figures and major analyses from this manuscript has been deposited on GitHub (https://github.com/WALL-E-Lab/Compocyte_reproducibility & https://github.com/WALL-E-Lab/Suco_reproducibility).

## Acknowledgements

We thank our dear coworker and friend Ignas Masilionis who pioneered the CITE-seq data generation for this project but was unable to witness the completion of this manuscript. PBMC samples were processed and provided by Heidelberg Cell and Liquid Biobank, a member of BioMaterialBank Heidelberg. We thank Umair Haroon for helping with the Compocyte readthedocs setup. We thank all patients who participated in this research. This project was supported by the following research grants: Excellence in Cancer Immunotherapy, Spanish Association Against Cancer (AECC, PI049999 to T.W.), Federal Ministry of Research, Technology and Space (BMBFTR, 001001KT2322 to T.W.), Stiftung für Krebs- und Scharlachforschung (to T.W.), German Cancer Aid (70114608, to. C.B.). National Cancer Institute (NCI) grants R35 CA263816, U24 CA213274, and P30 CA008748 (all to C.M.R.), Research Council of Lithuania (P-MIP-24-93 to L.M.).

## Author contributions

Conceptualization: T.W., Main computational methodology (Suco, Compocyte, Cytopus): T.W., C.B., L.S.; other Methodology: T.W., S.A.R, J.M.C., R.C.; Data analysis: C.B., T.W., Z.Q., A.A.; Data generation: C.B., T.W., A.Q.-V., P.M., J.A.K., A.L., S.B., B.M., S.U., L.M., J.N., E.d.S., R.C; Provided patient samples: T.W., J.A.K. K.A.K., C.S.C., S.M.K., K.D., A.W., M.J., T.P., S.K., N.H., D.P.N., N.S.M., E.Z., A.K.B., C.S., S.Z., J.C.H., J.D., D.J., C.A.I.-D., K.G., G.U., G.M.H., C.M.R.; Writing-original draft: T.W.; Writing-review and editing: C.B., T.W.; Visualization: C.B., T.W.; Funding acquisition: D.P., C.A.I.-D., C.M.R., G.U., P.E.H., T.W.; Supervision: D.P., C.A.I.-D., C.M.R., G.U., P.E.H.,T.W.; Read and approved final manuscript: All authors.

## Declaration of generative AI and AI-assisted technologies in the writing process

During the preparation of this work the authors used Google Gemini Pro 3.1 (01.05.2026-20.05.2026) in order to improve readability and English language. After using this tool, the authors reviewed and edited the content as needed and take full responsibility for the content of the published article.

