## Supplementary Figures for "Hierarchical classification of immune cell transcriptomes at population-scale"

### Supplementary Figures | Beltz et. al.

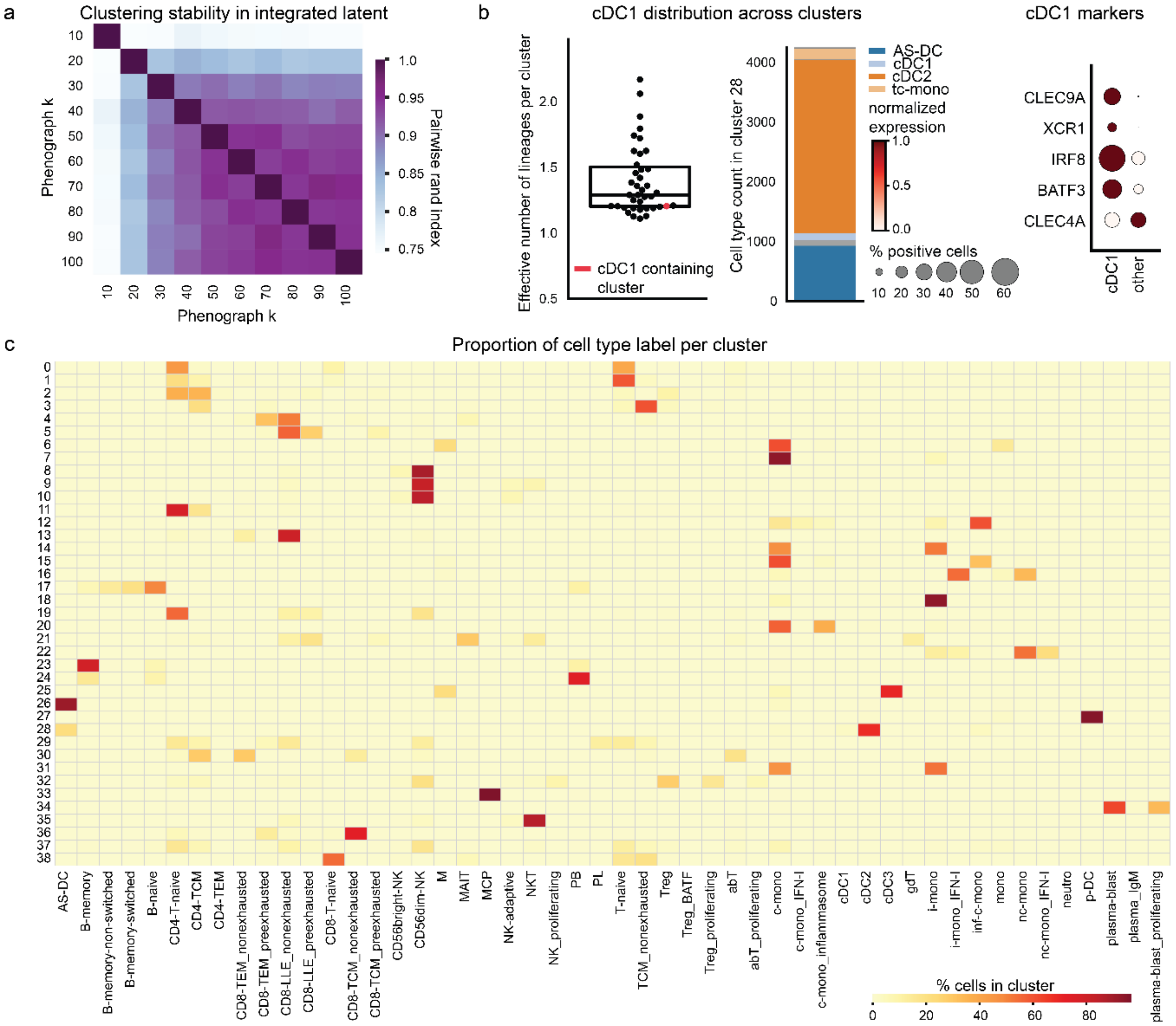


**Fig. S1 | Rare cell type clusters are lost during unsupervised data integration| a**, We integrated a 4-dataset subset of Suco PBMC resource (Methods) using scVI and clustered cells using Phenograph. K=30 resulted in granular, stable clustering as shown by the pairwise rand index. **b**, Conventional dendritic cells type 1 (cDC1) fall into a cluster with a low degree of mixing as determined by the effective lineage number (scatterplot in red), they only represent a minority of the cell types in this cluster (bar graph, middle). Other cells in the cluster did not show cDC1 marker expression confirming that cDC1 would be lost in unsupervised data integration. **c**, Many clusters obtained from clustering in scVI latent space contain multiple cell types. tc: transitional classical, mono: monocyte, NK: Natural Killer, MAIT: mucosa-associated invariant chain T cell, AS DC (AXL & SIGLEC6 positive dendritic cell), baso (basophil), cDC (conventional dendritic cell), HSC (hematopoietic stem cell), MAIT (mucosa associated invariant chain T cell), mono (monocyte), NK (Natural Killer), tc (transitional classical), B (B cell), c mono (classical monocyte), DN (double negative), eosino (eosinophil), ery (erythrocyte), exh. (exhausted), gdT (gamma delta T cell), gran (granulocyte), IFN (interferon), ILC (innate lymphoid cell), i mono (intermediate monocyte), M (myeloid cell), nc mono (non classical monocyte), neutro (neutrophil), PB (plasma or B lineage), prol. (proliferating), T (T cell), TCM (T central memory), tc-mono (transitional classifcal monocyte) TEM (T effector memory), term. (terminally), TNK (T or innate lymphoid cell), Treg (regulatory T cell).

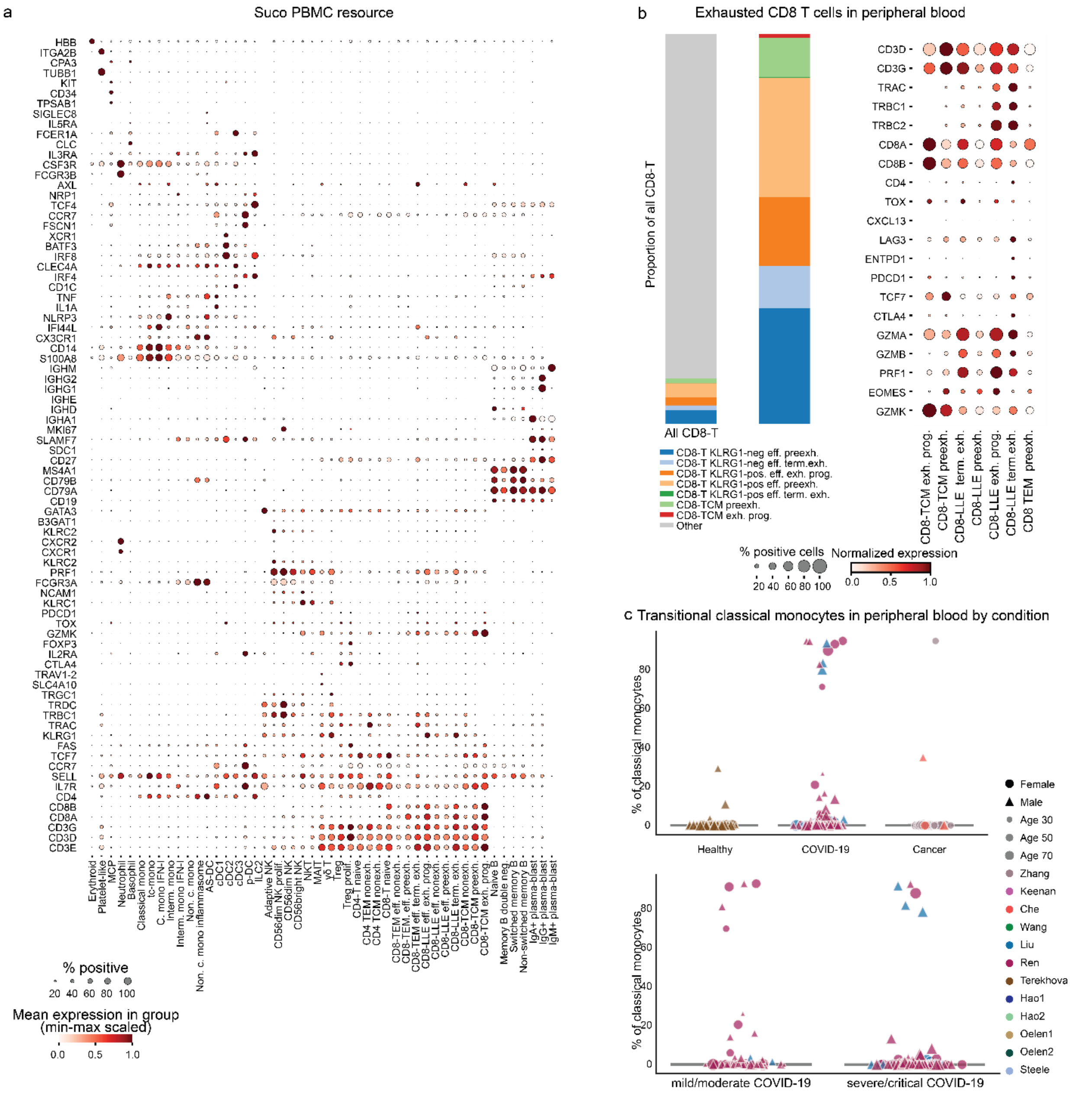


**Fig. S2 | Cell types in the Suco peripheral blood mononuclear cells resource | a**, Dotplot showing marker gene expression leaf-node labels from our Suco peripheral blood mononuclear cell (PBMC) resource (n= 5,389,645 cells). Transitional classical monocytes (tc-mono) express monocytes but no other lineage markers. B (B cell), c mono (classical monocyte), DN (double negative), eosino (eosinophil), ery (erythrocyte), exh. (exhausted), gdT (gamma delta T cell), gran (granulocyte), IFN (interferon), ILC (innate lymphoid cell), i mono (intermediate monocyte), M (myeloid cell), nc mono (non classical monocyte), neutro (neutrophil), PB (plasma or B lineage), prol. (proliferating), T (T cell), TCM (T central memory), tc-mono (transitional classifcal monocyte) TEM (T effector memory), term. (terminally), TNK (T or innate lymphoid cell), Treg (regulatory T cell). **b**, Stacked barplots showing the proportions of exhaustion states in CD8-T cells in Suco PBMC, and dotplot showing their marker gene expression. Only, CD8 LLE span the entire spectrum of exhaustion states exh.: exhausted, term.: terminally, preexh.: preexhausted, prog.: progenitor . c, Proportion per sample (n=1560) of transitional classical monocytes are present across diseases, ages, datasets, sex.

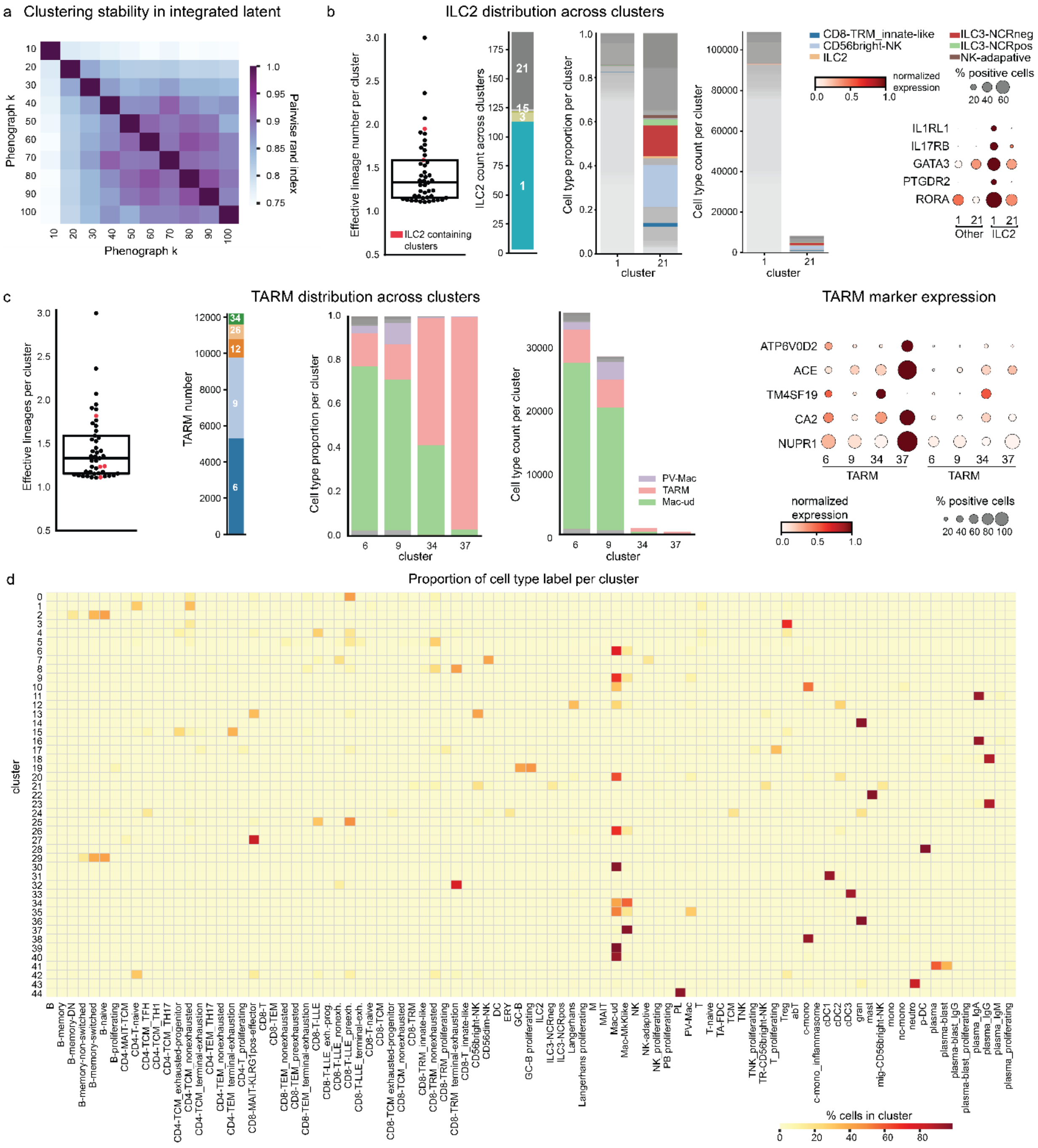

**Fig. S3 | Granular cell types are lost in unsupervised data integration | a**, We integrated our Suco TIL data using scVI and clustered the data using k=60 PhenoGraph clustering as a stable but granular clustering as determined by the pairwise rand index. **b**, Label homogeneity as judged by the effective number of lineages per cluster shows how innate lymphoid cells type 2 (ILC2) are assigned to heterogeneous clusters 1 and 21 (left, in red). ILC2s are distributed across several clusters (2nd plot from the left), and even in the clusters containing the most ILC2, they co-occur with other cell types (3rd and 4th plot from left). Cell type marker gene expression (plot on the right) confirmed that the remaining cells in the clusters were not of ILC2 lineage. **c**, Many tumor-associated resorptive macrophages (TARM) containing clusters (in red) showed strong cell type heterogeneity . Clusters 6 and 9 contained most TARM (2nd plot from the left), but are dominated by other macrophage subtypes (3rd plot from the left). Other cells in the cluster generally lacked TARM marker gene expression confirming that TARM will be confused with other cell type during unsupervised data integration. **d**, Many clusters from integrated data contain multiple cell types.

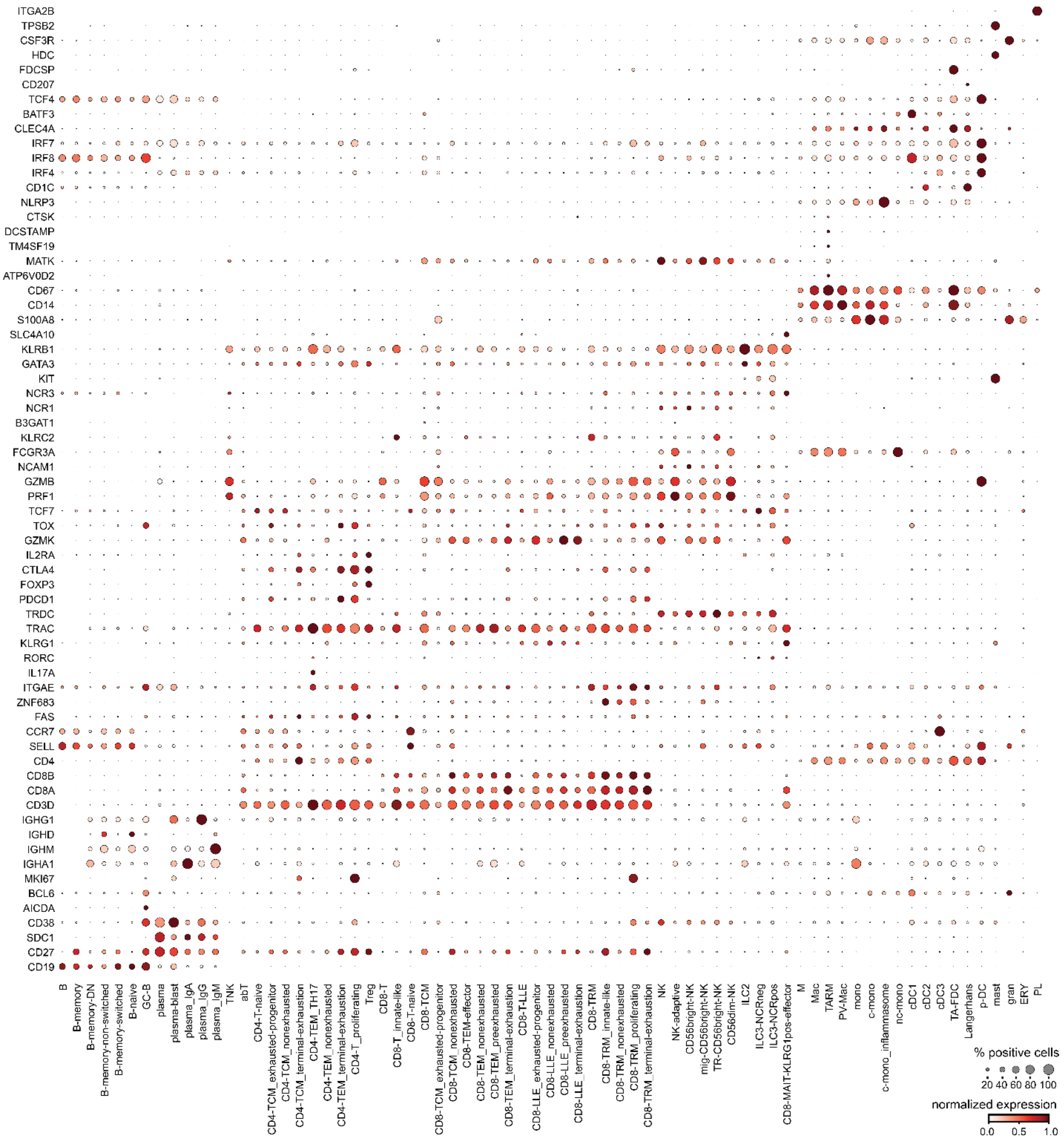


**Fig. S4 | Marker gene expression in tumor infiltrating leukocytes |** Dotplot showing expression patterns of different cell types across in our Suco tumor-infiltrating leukocyte (TIL) resource (n=764,571 cells). CD56bright NK cells frequently show a tissue resident phenotype marked by ITGAE and ZNF683 alongside an absence of SELL. TARM lacked canonical dendritic cell markers (CD1C, CD1A, BATF3, IRF7, IRF4, IRF8). c- (classical), cDC (conventional dendritic cell), DN (double negative), exh (exhausted), GC (germinal center B cells), gran (granulocyte), ILC (innate lymphoid cell), innate like (innate like T cells), LLE (long lived effector), M (myeloid), Mac (Macrophage), MAIT (mucosa associated invariant chain T cells), mem (memory), mono (monocyte), nc (non classical), neutro (neutrophil), NK (Natural Killer cell), ns (non switched), PB (plasma or B cell lineage), plasma (plasma cell), prog (progenitor), prol. (proliferating), PV (perivascular), sw. (switched), T (T cell), TA FDC (tumor associated follicular dendritic cell), TARM (tissue associated resorptive macrophages), TEM (T effector memory), TH17 (T helper 17), TNK (T or ILC lineage).

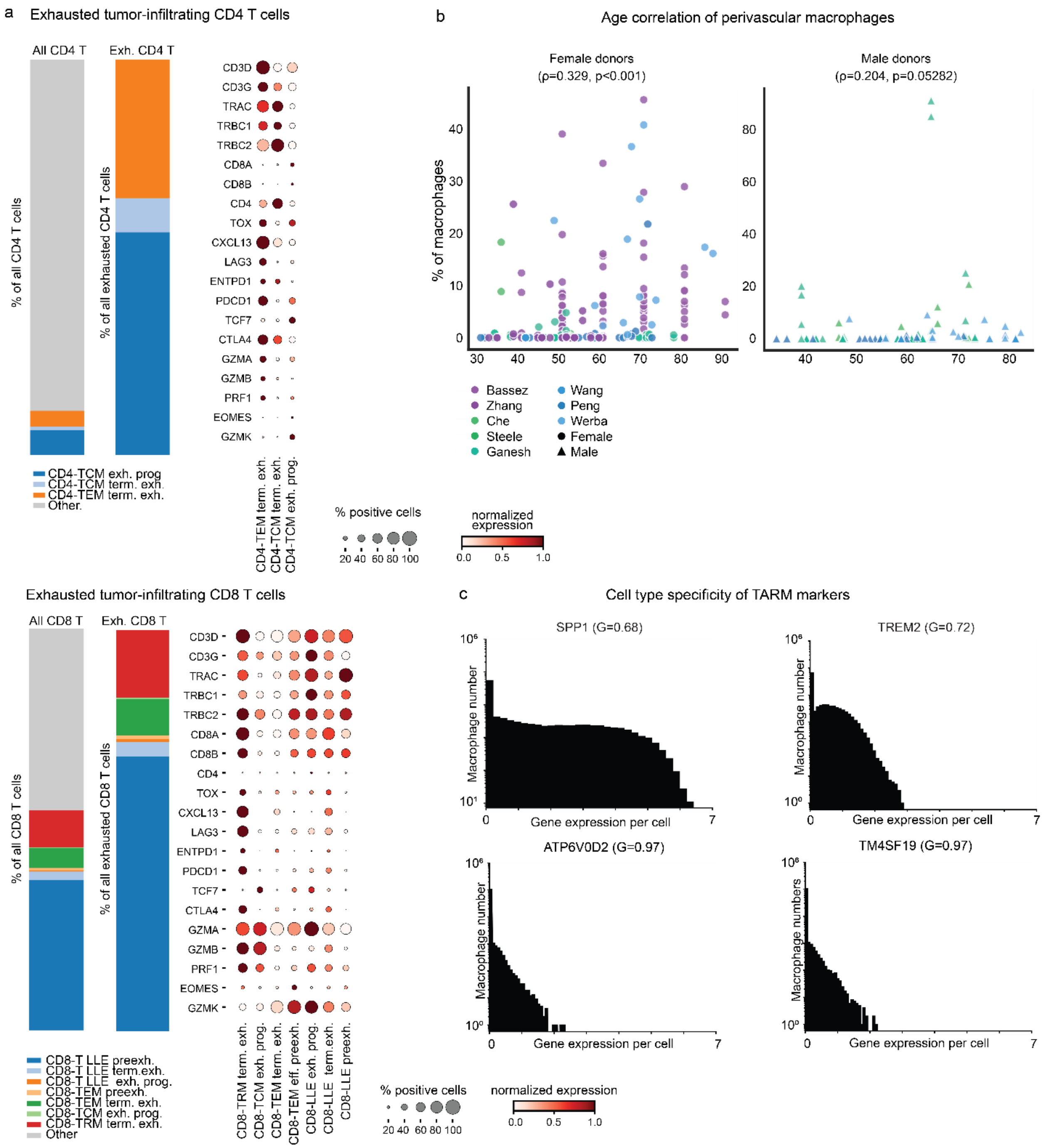


**Fig. S5 | Tumor infiltrating leukocytes in Suco** **a**, Stacked barplots showing the proportions of exhausted subset in CD4-T (upper panel) or CD8 T cells (lower panel) in our Suco TIL atlas. Dotplot showing the normalized exhaustion marker gene expression. TRM: tissue-resident memory T cells, TCM: central memory T cells, LLE: long-lived effector T cells, term.: terminally, exh.: exhausted, prog.: progenitor, preexh.: pre-exhausted. **b** Dotplots showing the correlation between the proportion of perivascular relative to all tumor-infiltrating macrophages by age, dataset and sex with statistically significant positive correlation between age and the PV-Mac proportion in female donors. Correlations were quantified using the Pearson correlation coefficient. **c**, Distributions of normalized, log1p-transformed gene expressions in macrophages show a higher Gini coefficient for TARM-specific genes such as ATP6V0D2 and TM4SF19 as compared to TREM2 and SPP1.

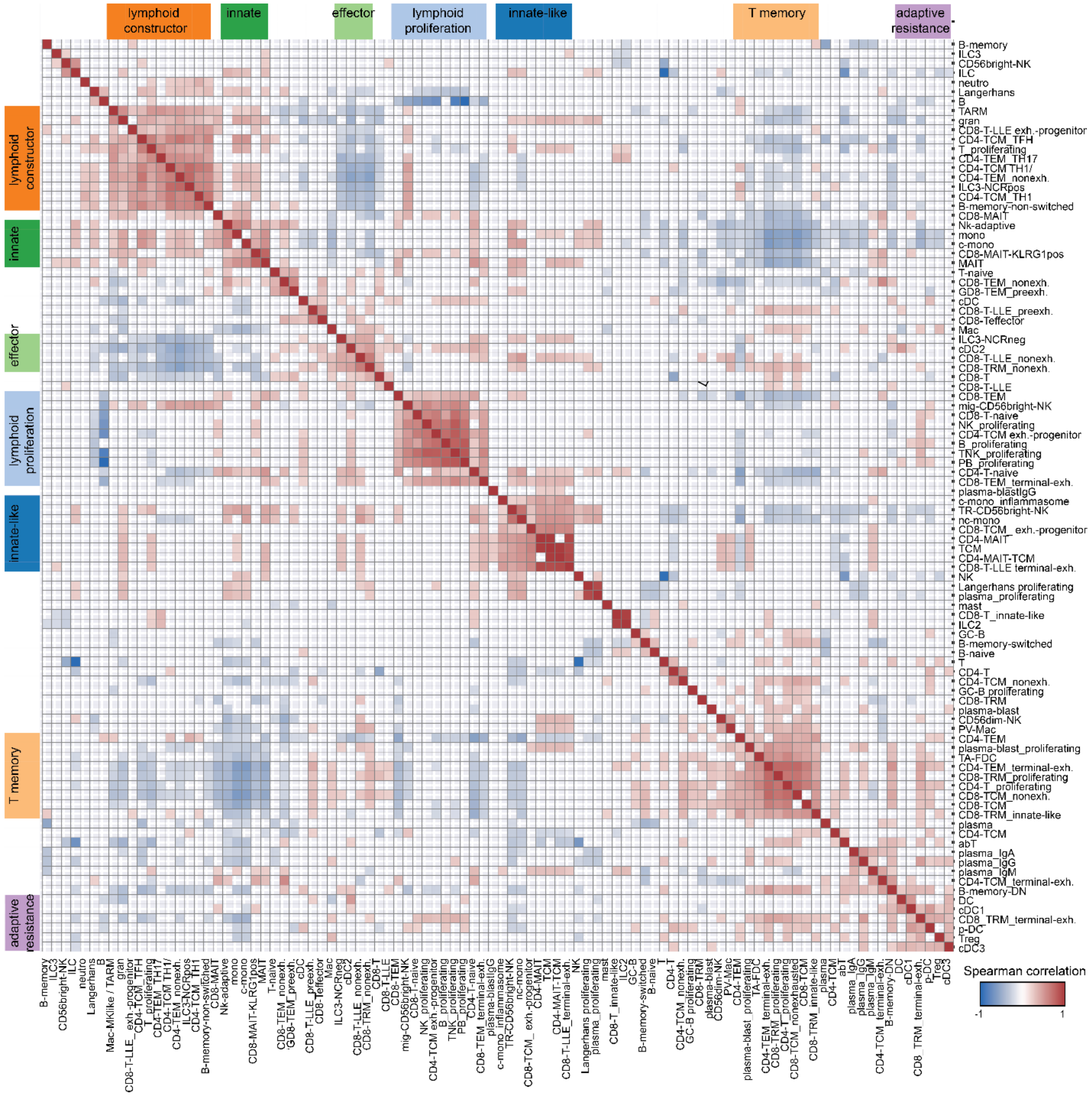


**Fig. S6 | Cell type co-occurence identifies tumor immune microenvironments |** We quantified cell type co-occurence by pairwise spearman rank correlation based on cell type frequency in our Suco TIL atlas (n=310 patient samples, Methods). This uncovered the tumor immune microenvironments (TIME) indicated with a color bar. Non-signficant correlations are crossed out. c- (classical), cDC (conventional dendritic cell), DN (double negative), exh (exhausted), GC (germinal center B cells), gran (granulocyte), ILC (innate lymphoid cell), innate like (innate like T cells), LLE (long lived effector), M (myeloid), Mac (Macrophage), MAIT (mucosa associated invariant chain T cells), mem (memory), mono (monocyte), nc (non classical), neutro (neutrophil), NK (Natural Killer cell), ns (non switched), PB (plasma or B cell lineage), plasma (plasma cell), prog (progenitor), prol. (proliferating), PV (perivascular), sw. (switched), T (T cell), TA FDC (tumor associated follicular dendritic cell), TARM (tissue associated resorptive macrophages), TEM (T effector memory), TH17 (T helper 17), TNK (T or ILC lineage).

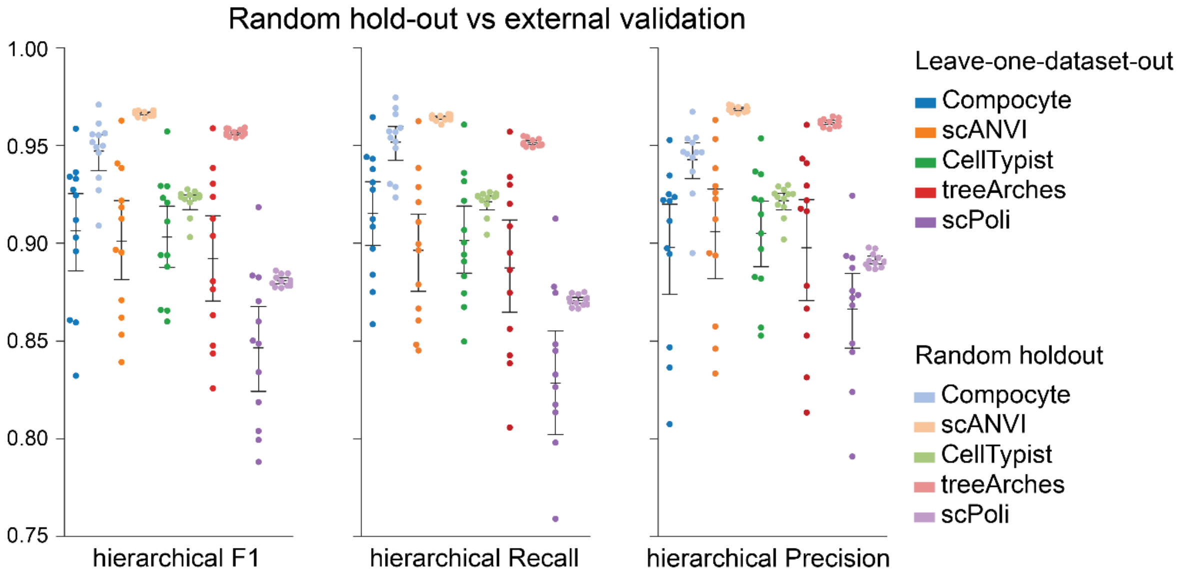


**Fig. S7 | Cell-level holdout overestimates model generalizability |** We simulated equivalent experiments by holding out dataset-sized randomly selected cells (random hold-out, lighter colors) which overestimated the ability of classifiers to generalize, as compared to dataset-level holdouts (darker colors).

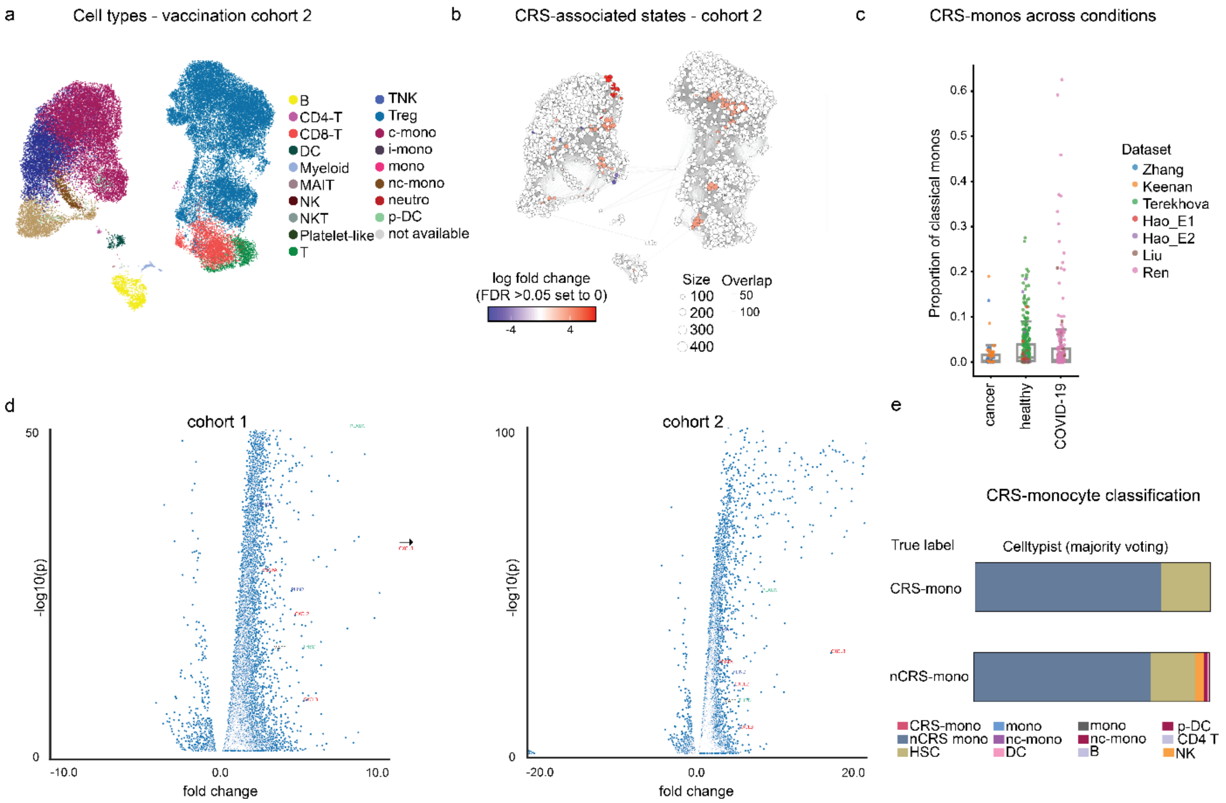


**Fig. S8 | Validation of subclinical cytokine release syndrome-associated monocytes in an independent cohort | a**, UMAP showing cell types in cohort 2 n=6 immune checkpoint therapy (ICT) treated cancer patients undergoing SARS-CoV-2 vaccination. **b**, Specific monocyte neighborhoods (CRS-monos) were significantly associated with high CRS-associated cytokine levels as revealed by Milo (false discovery rate (FDR)=0.05). **c**, Using our Compocyte classifier we quantified CRS-monos in 7 PBMC datasets totaling 775 samples across a total of 389 healthy donors, patients with cancer and active infectious disease. **d**, Volcano plots showing differentially expressed (Mann-Whitney U tests) genes in CRS-monos as compared to other classical monocytes. Genes with fold changes outside of the axis are indicated with an arrow. **e**, Proportion of predicted cell types in the unseen SARS-CoV-2 vaccination cohort 2 by Celltypist with majority voting (n=6 patients) for CRS-monos and nCRS-monos.

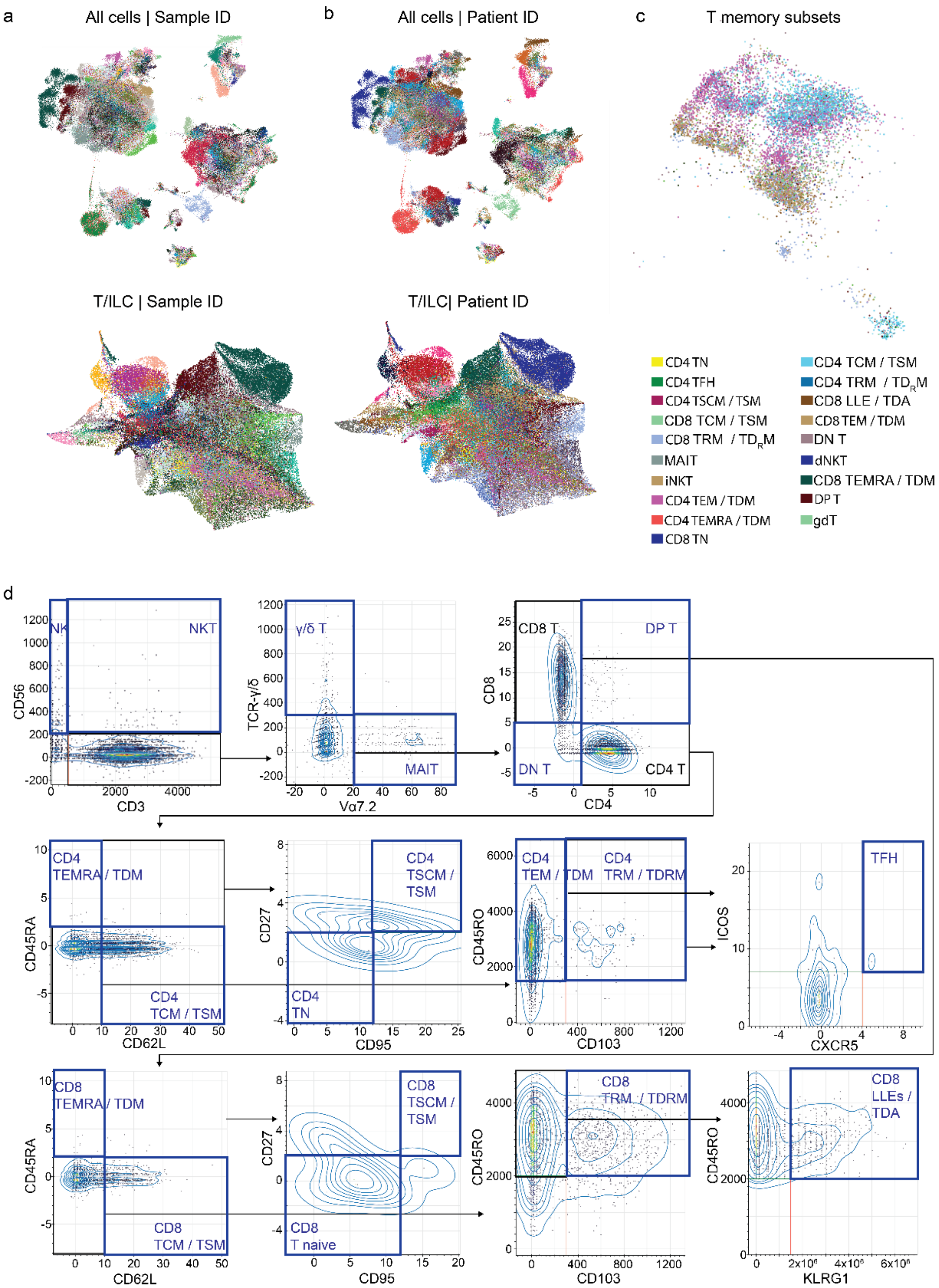


**Fig. S9 | CITEseq analysis of tumor infiltrating leukocytes | a,** UMAP embeddings of tumor infiltrating leukocytes (n=150,059) or tumor infiltrating leukocytes (n=84,222) colored by sample or by **b**, patient. **c**, UMAP of one representative patient calculated on gene expression alone with color code indicating T cell memory subsets manually gated on surface protein CITE-seq data. **d**, T and innate lymphoid cells in CITE-seq samples were manually gated per sample according to the indicated gating strategy on DSB antibody-derived tag data with leaf-node cell types indicated with blue gates and carried-forward populations in black. NK: Natural Killer cell, NKT: Natural Killer T cell, TEMRA: Effector memory RA T cell, TCM: Central memory T cell, TSCM: Stem cell memory T cell, LLE: Long-lived effector cell, TRM: Tissue-resident memory T cell. We also indicate a modular T cell nomenclature: A: activated, D: disseminated, S: Secondary lymphoid organ homing, M: memory, R: resident. Mean Spectra cell score per cell type and sample in metastases versus primary tumors in our pancreatic and lung cancer cohort (n=39 patients).

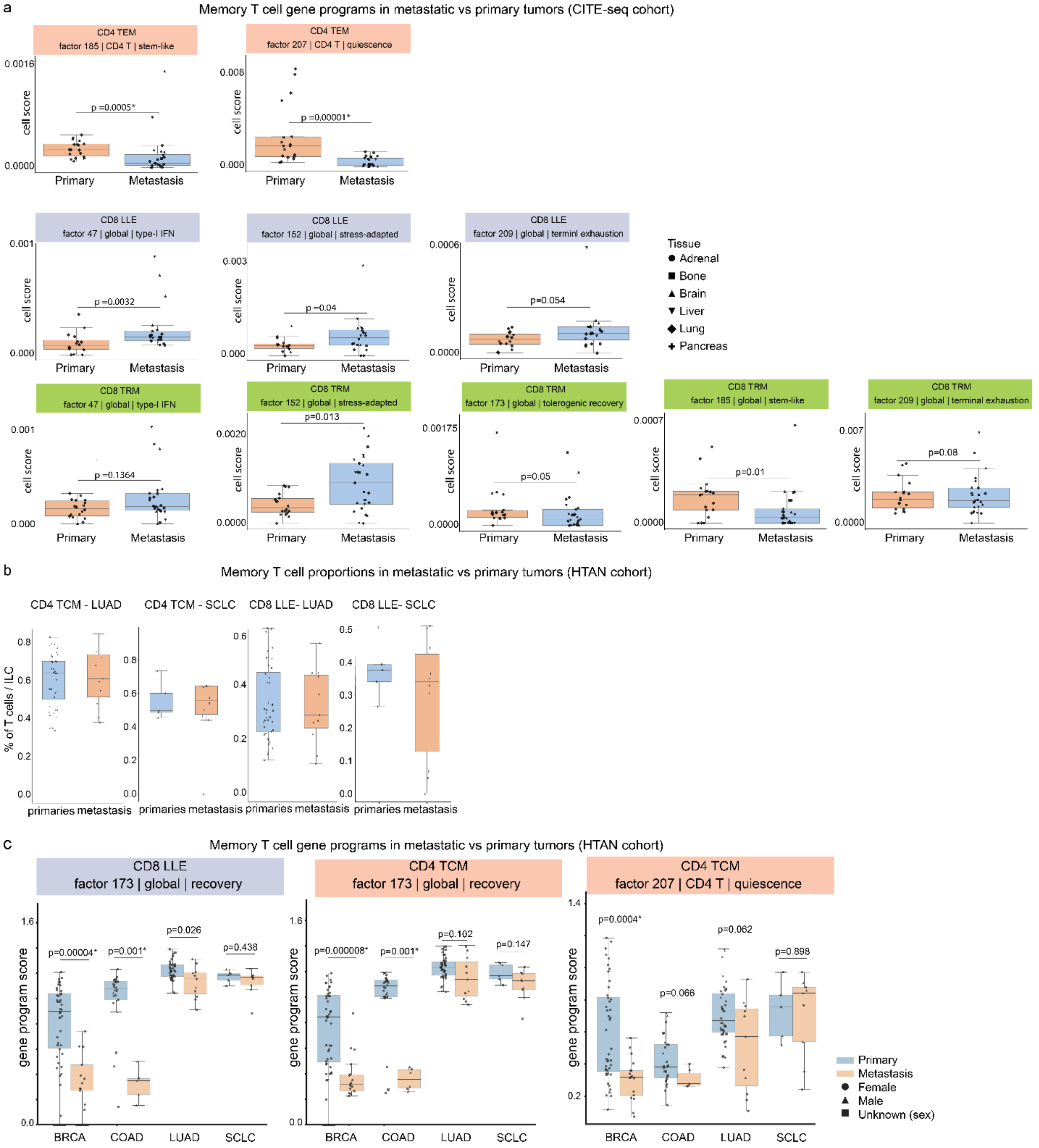


**Fig. S10 | Gene expression programs in memory T cells across tumor types | a,** Mean Spectra cell score per cell type and sample in metastases versus primary tumors in our pancreatic and lung cancer cohort (n=46 samples). **b**, Frequencies of indicated memory subsets in tumor types from the Human Tumor Atlas Network (HTAN) cohort (n=160 patients). **c**, Mean gene program scores in biopsies from metastatic and primary tumors across cancer types (breast cancer (BRCA), colorectal adenocarcinoma (COAD), lung adenocarcinoma (LUAD), small cell lung cancer (SCLC)) in the HTAN cohort (n=160 patients). All p values were calculated using Mann-Whitney U tests with significant comparisons highlighted with an asterisk according to the Benjamini-Hochberg method.

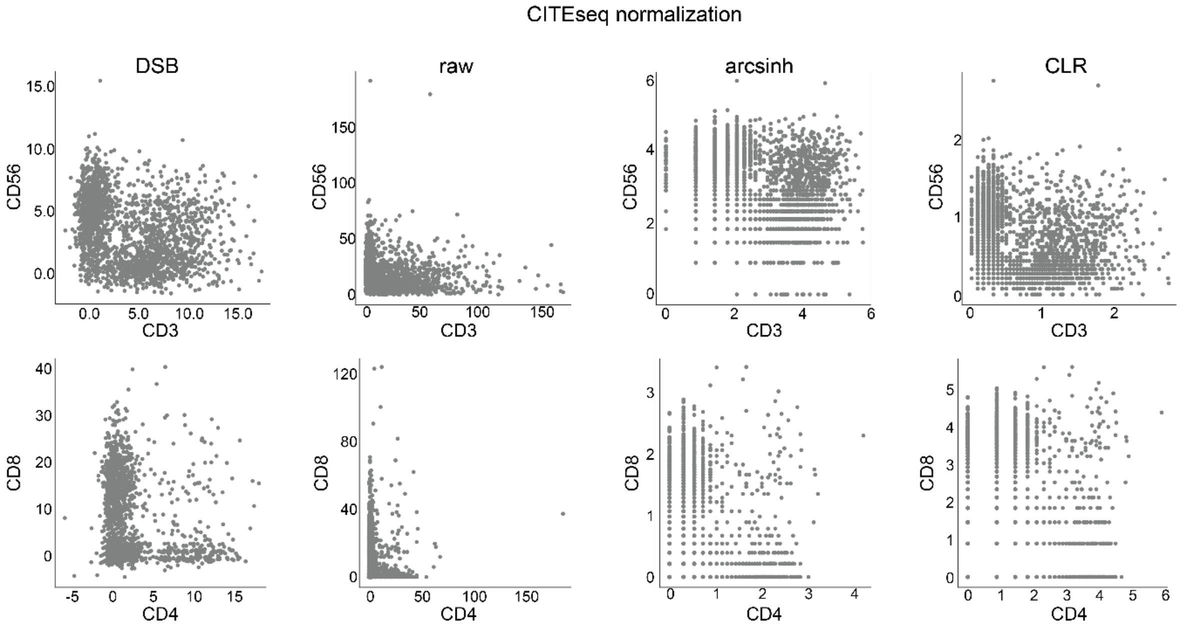


**Fig. S11 | Normalization of CITE-seq data** | Comparison of immune cell subset separation by different CITEseq normalization methods (raw, DSB normalization, arcsinh normalization, CLR normalization) shows DSBs ability to identify major immune cell lineages (n=1 representative patient)
