## Supplementary Note for "Hierarchical classification of immune cell transcriptomes at population-scale"

**Supplementary Note | Beltz. et. al.**

### Gene programs differentially expressed in primary tumors and metastases (Human Tumor Atlas Network CITE-seq dataset)

<u>Global factors – expressed in all cells</u>

**39 – hypoxia response:** This program strongly overlapped with our hypoxia input gene set and was particularly expressed in CD8 TRM and CD8 LLEs in metastases as compared to primary tumors (**Fig. 6c**). It encompasses core glycolysis genes (*ENO, GAPDH, PFKFB3, PFKL, PGK1*), transcriptional master regulators of hypoxia (*VHL, CITED2*) and mediators of cell cycle arrest (*CDKN1A, CDKN1B,* **Table S10**).

**47 – type I IFN signaling:** CD8 LLEs and TRMs showed elevated type I interferon signaling in metastases as compared to primary tumors (**Fig. 6c**, **S10a**). This program captured typical IFN-stimulated genes from our input set, featuring *IFI6, IFI44L, ISG15, IFITM1, IFITM2, IFITM3, IRF1,* and *IFIH1* among its top 10 marker genes (**Table S10**).

**106 – MHC-I presentation:** Across CD4 T cells, this program was more highly expressed in metastases compared to primary tumors (**Fig. 6c**). The factor successfully captured the majority (8 of 10) of genes from our MHC-I antigen presentation input set, specifically including the structural HLA molecules (*HLA-A, HLA-B, HLA-C*), the peptide transporters (*TAP1, TAP2*), and the peptide-loading complex chaperones (*TAPBP, CALR, PDIA3,* **Table S10**).

**131 – ROS response:** Strongly overlapping with our reactive oxygen species (ROS) detoxification input set, this program captures the thioredoxin system (*PRDX1, PRDX2, PRDX4, PRDX6, TXN, TXNRD1, TXNRD2*), the glutathione system (*GCLC, GCLM, GSR, GLRX, GLRX2, GPX4, MGST1*), and direct ROS scavengers (*CAT, SOD1, MPO*). It exhibited higher expression, across most CD4 and CD8 T cell subtypes in metastases compared to primary tumors (**Fig. 6c**, **Table S10**).

**152 – stress-adapted activation:** Enriched in metastasis-associated CD8 long-lived effectors (LLE) and tissue-resident memory cells (TRM), this program merges markers of tumor reactivity and T cell receptor (TCR) activation (*ENTPD1, TRAF1, NFKB2, NFKBIE, KRAS, CRKL, BHLHE40*) with profound metabolic adaptation. It highlights a shift toward beta-oxidation (HIF1A, PPARA, SLC20A2, ME2, HADHA) to navigate hypoxic, low-glucose niches, alongside robust activation of the unfolded protein response (*ATF6, CALR, HSPA9, DNAJB12, AMFR, ERLIN1, QSOX1*) (**Fig. 6c**, **S10a**, **Table S10**).

**165 – Metabolic blast:** Capturing the metabolic reprogramming required for a highly active and proliferating T cell state, this program highlights active lipid remodeling (*PLPP3, MGLL, PIGS*), DNA biosynthesis (*DHFR, RFC4, CKS1B*), and sustained inflammatory signaling (*TRAF6, NFKBIB*). It was predominantly expressed in metastases by CD4 TRM and TEM (**Fig. 6c**, **Table S10**).

**166 – Tissue residence:** Predominantly expressed in tissue-resident memory (TRM) T cells (**Fig. 6c**), this program features tissue-anchoring chemokine receptors and integrins (*CXCR4, CXCR6, ITGA2*) alongside mediators of cell cycle arrest (*BTG1, BTG3*) and metabolic stress defense (*FTH1, TSPO*). Notably, *COMT* expression suggests an adaptation allowing these cells to withstand catecholamine-rich microenvironments. Furthermore, robust upregulation of cytoskeletal remodeling transcripts (*ACTB, PFN1, CFL1, ARPC3, MYL12A, MYL12B, VIM*) supports their physical structural adaptation for tissue retention (**Table S10**).

**173 – Tolerogenic recovery:** Prominent in primary tumor CD8 LLEs and TRMs (**Fig. 6c,d, S10a**,**c**), this program reflects cellular responses to genomic damage (*TP53, TP53INP1*) and ER stress (*EDEM2, MAN2A1, B4GALT1*). Uniquely, it features *SMAD2*, a transducer of TGF-β signaling, and *JAG1*, which has been implicated in driving tolerogenic immune interactions and regulatory T cell generation (**Table S10**).

**185 – Stem-like:** Elevated across primary tumor memory subsets (CD4 TCM/TEM and CD8 LLE/TRM, **Fig. 6c,d, S10a**), this program shows long-term survival markers (*IL7R*) and enforced quiescence, restricting both cell cycle progression (*BTG1*) and glucose uptake (*TXNIP*). High expression of *CXCR4* may aid in retaining these cells within tertiary lymphoid structures or the bone marrow. Alongside *JUNB* and *CREM* which are shared with the quiescence program 207, this state employs mRNA decay (*ZFP36, ZFP36L2)* and signaling regulators (*TNFAIP3, PDE4B*) to dampen TCR-induced activation and maintain a resting, wound-healing-like state (**Table S10**).

**187 – TNF response:** Showing strong overlap with our TNF-alpha input gene set, this program was particularly enriched in primary tumor-infiltrating cells across all CD4 and CD8 memory subtypes (**Fig. 6c**, **Table S10**).

**207 – Quiescence**: In primary tumor CD4 TEMs and TCMs (**Fig. 6c,d, S10a**,**c**), we observed higher expression of this new quiescence program, which shares survival (*IL7R*) and anti-proliferative (*BTG1, JUNB, CREM*) factors with the stem-like program 185. However, it is distinctively enriched for anti-apoptotic mediators (*RTKN2*), internal signaling brakes (*PIK3IP1*), and co-inhibitory receptors (*TIGIT, CH25H*). The presence of *SESN1, SESN3*, and *SOD1* likely endows these cells with enhanced resistance to oxidative stress (**Table S10**).

**211 – Uncommitted:** This program particularly observed in CD8 TCM and TEM in metastases (**Fig. 6c**), is driven by genes that oppose tissue residency (KLF2[1], *GPR171*) and transcription factors that maintain an undifferentiated, poised state (*ETS1, TLE4*), from which cells can eventually acquire effector or TH17 polarization (*CAMK4* [2]). Simultaneously, these cells remain in a metabolically resting profile, reliant on baseline oxidative phosphorylation (*PIK3IP1, COX7C, UQCRB, TXNIP*, **Table S10**).

CD4 T cell specific factors

**194 – CD4 immunological synapse formation:** Enriched in metastasis-associated CD4 TRM and TFH cells (**Fig. 6c**), this program partially overlaps with our IL-12 signaling input set, specifically capturing genes critical for immunological synapse formation (*LCP1, RHOG*). This functional assignment is further supported by the expression of the glutamine transporter *SLC38A2*, the common gamma chain cytokine receptor (*IL2RG*), and an extensive network of actomyosin cytoskeletal regulators (*ACTB, ACTR2, MYL12A, MYH9, FLNA, CORO1B, AHNAK*, **Table S10**).

**196 – Tumor reactivity**: We discovered this novel, metastasis-specific tumor reactivity program in CD4 T cells (**Fig. 6c**). While we previously reported a tumor reactivity program in CD8 T cells[3], this CD4 program shares core activation features with its CD8 counterpart, including the transcription factor *BATF*, effector cytokine *IFNG*, co-stimulatory receptor *TNFRSF18*, cell stress mediator *GADD45G*, and TCR signaling kinase *LCK*. It also includes PDCD1 suggesting the cells expressing this program may be targeted by immune checkpoint therapy. However, it is distinguished by CD4-specific mechanisms, including dendritic cell activators (*CD40LG, CSF2, XCL1*), death-receptor ligands (*TNFSF10, TNFSF14*), lysosomal cathepsins (*CTSC, CTSH, CTSL*, **Table S10**).

CD8 T cell specific factors

**209 – CD8 terminal exhaustion:** Particularly expressed in metastatic CD8 TRMs (**Fig. 6c**, **S10a**), this program maps to our terminal exhaustion input set (capturing 4 of 13 genes). It features the immune checkpoints *LAG3* and *HAVCR2*, alongside exhaustion-driving transcription factors *EOMES* and *IRF4*. Furthermore, Spectra identified additional severe exhaustion markers in this state, including *CTLA4, CD200R1*, and *LAYN*, as well as TCR-attenuating genes *DUSP4* and *SLA*. The presence of *KLRC1, KLRC2,* and *KLRD1* among the top 50 markers also indicates a degree of innate reprogramming (**Table S10**).

**210 – CD8 progenitor exhaustion:** Predominantly expressed in primary tumor CD8 LLEs and TEMs (**Fig. 6c**), this program strongly overlaps with classical exhaustion and stem-like input genes. It captures the exhaustion master regulator *TOX*, the gold-standard progenitor marker *SLAMF6*[4]*, PDCD1*, alongside stemness preservers *BCL6* and *ID3*. Notably, although *TCF7* was absent from the input set, Spectra successfully recovered this definitive progenitor master regulator directly from the data, placing it as a core driver of this program (**Table S10**).

**212 – CD8 tumor reactive** Enriched in metastasis-infiltrating CD8 TRM cells (**Fig. 6c**), this program closely matches our previously published CD8 tumor reactivity factor (sharing 12 of the top 50 genes[3]), with modifications likely reflecting adaptation to the lung cancer microenvironment. This tissue-resident state includes the TLS-promoting chemokine *CXCL13*, the checkpoints *ENTPD1*, *LAG3*, and *TNFRSF9*, the residency marker *ITGAE*, and effector genes comprising *IFNG, GZMA, GZMB, GZMH, PRF1, GNLY,* and *CD8B (***Table S10**).

**219 – Epigenetic remodeling:** This program is defined by chromatin regulators (*H2AFZ, HIST1H4C, ZNF331, ZNF90*) which, alongside the transcriptional repressor *CREM*, likely poise the epigenome while restricting active transcription. The co-expression of *CCR7* and *IL7R* suggests a memory-like phenotype optimized for survival and persistence in the blood and lymphoid organs (**Table S10**). We found this program to be particularly expressed in primary tumors as compared to metastases (**Fig. 6c**).

### Supplementary Note References


1. Fagerberg, E., Attanasio, J., Dien, C., Singh, J., Kessler, E.A., Abdullah, L., Shen, J., Hunt, B.G., Connolly, K.A., De Brouwer, E., et al. (2025). KLF2 maintains lineage fidelity and suppresses CD8 T cell exhaustion during acute LCMV infection. Science *387*, eadn2337. doi:10.1126/science.adn2337.
2. Koga, T., Hedrich, C.M., Mizui, M., Yoshida, N., Otomo, K., Lieberman, L.A., Rauen, T., Crispin, J.C., and Tsokos, G.C. (2014). CaMK4-dependent activation of AKT/mTOR and CREM-alpha underlies autoimmunity-associated Th17 imbalance. J Clin Invest *124*, 2234-2245. 10.1172/JCI73411.
3. Kunes, R.Z., Walle, T., Land, M., Nawy, T., and Pe'er, D. (2023). Supervised discovery of interpretable gene programs from single-cell data. Nature Biotechnology. 10.1038/s41587-023-01940-3.
4. Gago da Graca, C., Sheikh, A.A., Newman, D.M., Wen, L., Li, S., Shen, J., Zhang, Y., Gabriel, S.S., Chisanga, D., Seow, J., et al. (2025). Stem-like memory and precursors of exhausted T cells share a common progenitor defined by ID3 expression. Sci Immunol *10*, eadn1945. 10.1126/sciimmunol.adn1945.
